# Strain resolution and wide relative abundance quantification of bacterial microbiomes with nanopore sequencing of the ribosomal RNA operon

**DOI:** 10.64898/2026.10.18.759976

**Authors:** Christopher J. Woodruff, Terence P. Speed

## Abstract

Identification and quantification of microbiomes at strain level taxonomic resolution is increasingly sought. Longread sequencing of sufficiently high accuracy can provide this using ribosomal RNA operon (rrn) sequencing, subject to strains differing by at least one base in some of their operons. Prior studies of nanopore-sequenced amplicon reads have not established reliable strain level resolution and sensitive quantification. Here we demonstrate that if a sufficient number of rrn operon reads with a mean error rate of better than about 0.008 errors per base are available in a dataset, then strain identification and quantification, mediated by operon-level resolution, can often be achieved. Two multiple dataset sources of real, nanopore-sequenced bacterial mock microbiome were accessed, and multiple datasets of a realistic mock human gut microbiome, using state-of-the-art simulated nanopore-sequenced reads, were simulated. These datasets had read quality distributions lying between those of the real datasets, did not have multiple strains for any species, but, for the higher quality datasets, allowed demonstration of within-strain operon resolution and quantification over 2 orders of magnitude. The simulation-generated datasets demonstrated multiple cases of within-species strain resolution with accurate relative abundance quantification. The analysis of such datasets has been implemented here in a pipeline of 4 multi-parameter command line calls. This allows intra-species resolution and quantification over at least 3 orders of magnitude with more than 50 strains present in the microbiome.

## 2 Introduction

Microbiomes of plants and animals (including humans) are today recognised as powerful mediators of important physiological systems’ states and responses to the environment [1], [2], [3]. They are also known to be manipulable [4], and hence, in the case of the various human microbiomes, potential tools to improve human health. The human gut microbiome is not only central to metabolic processes, but is also a known modulator of innate and adaptive immune systems [5], [6]. Recently evidence has grown of it having an important role in aspects of cognitive function [7], [8], presumably mediated via the vagus nerve [9]. Although much of the earlier published work on modulating the microbiome of plants and animals - and, in particular, humans - dealt with variations at the family or genus taxonomic levels, it is now clear that real targeting requires knowledge of microbiome interactions and manipulations at the strain level [10], [11], [4]. A first step in developing such knowledge is to determine what strains of organisms are present and to be able to monitor their relative abundances consequent on controlled or otherwise observed interventions or external environmental pressures.

Metagenomics approaches to profiling microbiota have gained favour in recent years, quite probably because of the considerable demands in determining what strains of various organisms are present. However the increased power of genomic sequencing consequent on the availability of long read sequencing technologies -specifically Pacific Biosciences (PacBio) HiFi capabilities, and the markedly improved ONT nanopore sequencers, has allowed increased taxonomic resolution in determining the microbiome profile - what organisms are present and in what relative abundance. The PacBio capability provides the leading edge capability in such profiling, but that technology is far less globally accessible (considering cost and location) than nanopore sequencing. Given the well-known variation in microbiome structure with environment, it is important for global health to have available a sufficiently capable technology that is more accessible than the PacBio systems. Nanopore sequencing is well proven as a technology that is accessible globally [12],[13] and beyond [14],[15]. If it can be shown that its capability is such as to be able to provide strain level taxonomic resolution for realistic microbiome structures this would be widely beneficial.

The pioneering work of Woese and colleagues enabled genomic-based microbiome analysis based on ribosomal RNA (rRNA). Analysis using hypervariable regions of the 16S ribosomal RNA gene has dominated subsequent microbiome analysis. The variability of such regions, however, has been inadequate for reliable resolution even at the species level. More recently, with the emergence of various technologies to get longer reads, greater resolution has been achieved using full length 16S rRNA genes, and using most or all of the full ribosomal RNA operon (rrn). Benitez-Paez and colleagues [16], [17], using an early version of the MinION nanopore sequence from Oxford Nanopore Technologies (ONT), demonstrated moderate species level taxonomic identification. Subsequently Cusco et al. [18] analysed dog skin microbiota using full 16S rRNA gene sequences, and also [19] compared results from 16S rRNA and the 16S-ITS-23S rrn operon. These studies showed somewhat inconsistent quantification between 16S rRNA gene and rrn operon datasets, with the rrn operon giving a poor representation of the designed mock microbiome analysed. Kerkhof et al. [20] used an iterative approach to develop a consensus sequence from reads forming OTU clusters and used this to estimate underlying 16S and 23S rRNA gene sequences of constructed microbiomes of greater complexity than standard mock microbiomes. Their analysis showed that species and, to a certain extent, strain level resolution was possible from both 16S and 23S rRNA gene sequences. They also demonstrated a limited quantification capability. These authors noted the limited coverage of databases other than for 16S rRNA genes. The taxonomic resolution achieved by their method was a clear advance on previous studies, but was strongly limited by the error rate of the nanopore sequencing of that time. It is of note that the iterative refinement from OTU clusters is somewhat analogous to the later method used in Kumar et al.’s Robust Amplicon Denoising [21] used extensively in the analysis reported in the Results section of this paper.

Achieving better taxonomic resolution based on sequences of characteristic genomic features, such as the rrn operon or its 16S and 23S rRNA genes, or the internally transcribed spacer (ITS), requires suitable databases of what are the corresponding sequences of organisms that are relevant. If strain level resolution is to be achieved for the human gut microbiome of both healthy and unhealthy people across ages and social groups then the reference database must have information at the strain level across the range of relevant organisms. Currently, this is far from the case. A key part of the problem is isolating organisms from which genomic material can be extracted. This is particularly so for organisms that survive only in specific environments, such as anaerobic bacteria. More recently, improvements in culturing have allowed greater representation of such bacteria. Metagenomic assembly of genomes has also enabled more extensive representation, with resolution at least to the species level. It is not clear, however, that reliable strain resolution can be achieved with this approach at present, particularly where strains differ by only a few bases across the whole rrn operon, or do not differ at all over the rrn operon but only at other genomic locations.

It is likely that quantitative understanding of microbiota interactions within a microbiome, and between a microbiome and its host, will depend on relative abundances of the interacting strains. That is, cellular abundances will be more relevant than rrn operon abundances. Much microbiota profiling gathers counts of operons. Determination of cellular abundances requires that the number of operons per genome is available. This requires that databases for species-level quantification be accurate in terms of the number of rrn operons present per genome of that species (or strain). There is a large difference between the number of species that have incomplete genomes in reference databases and those that have complete genomes. Methods are available for estimating the number of rrn operons, and for most purposes associated with human health it is likely that these are sufficiently reliable given current limitations in other aspects of quantitative modeling.

Here we propose the use of tools and methods that enable amplicon-based microbiome identification at the operon level using ONT nanopore technology, and, by condensing operon-level information across strains, providing both identification and quantification at the strain level. The overall method of analysis of a microbiome consists of i) Deriving exact or close matches to the sequences that have been the origin of the reads (sequences) in the library of reads following sequencing; ii) Aligning each such estimated original sequence (referred to as an Amplicon Sequence Variant, ASV) against a reference database of operons to identify the most likely of the database entries from which the original sequence was derived; iii) Associating as many as possible of the reads of the library with exactly one of the ASVs, and hence, using the alignment of ASVs to database entries, determining how many reads are deemed to have arisen from the various operons, strains, species or genera of the database. Denoising achieves item i) and the association component of item iii).

Our results come from mock microbiomes having either real read data or simulated read data. Briefly, we show the quality of the input data, what strains are found to be present, and how much (proportionately) of each strain is present. The results are presented largely through figures showing, for each dataset, i) the distribution of read mean base error rates for the library of reads presented to the denoising algorithm; ii) a two-dimensional representation of the inter-ASV dis-similarities for the operons deduced to be present based on the output from the denoising; and iii) scatterplots of observed versus actual relative abundances of strains for the mock microbiome.

## 3 Results

Two sources of real sequenced reads were used: extracted 16S and 23S rRNA sequences from a whole genome sequencing study by Sereika et al.[22] (See Methods and Supplementary Materials Section 1), and 4 datasets of RNA amplicons of ribosomal RNA for each of two mock microbiomes from Srinivas et al [23] (see Methods and Supplementary Materials Section 2). Multiple datasets of simulated reads were also generated, based on the healthy human gut microbiome proposed by King et al. [24] (see Methods). Wick’s badread ONT sequence simulator [25] was used to generate reads from which libraries of reads corresponding to those expected for particular mock microbiomes were constructed. Each dataset was denoised using the Robust Amplicon Denoising (RAD) algorithm [21], producing a set of estimated sequences, which we refer to (loosely) as Amplicon Sequence Variants (ASVs). These are aligned using blastn [26] to databases derived from [27], one of which only has entries that are resolved at the strain operon level. Further detail on the databases is in the Methods section.

### 3.1 Data Quality

Our primary measure of data quality is the mean(per-base-error-rate of a read), also represented by the Q-score, defined as

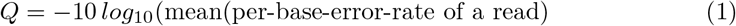

Figure 1 shows distributions of mean per base error rate for indicative datasets from each of the real data sources ([22], [23]) and two of the simulated datasets. We focus on the error rate region that is necessary for the denoising algorithm to give useful results (error rate ≤ 0.01, or *Q* ≥ 20).

**Fig 1.**
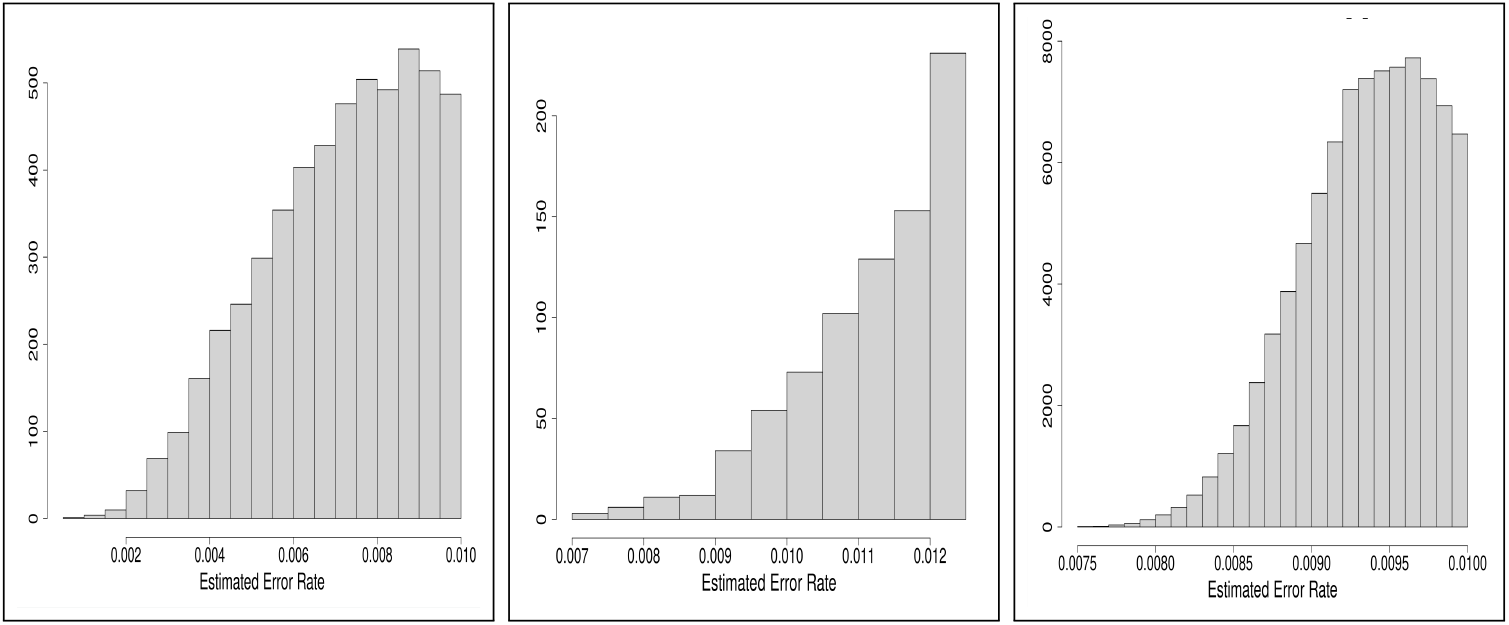
Error rate distributions indicative of real read datasets from (left) Sereika etal. and (middle) Srinivas et al, and (right) simulated reads dataset based on a healthy human gut microbiome [24] with mean Q=30.Note that the range of mean error rate for the middle plot is from a maximum of 0.0125, while the others are from a maximum of 0.10.

Figure 1 shows two important features of the datasets - firstly, our simulated data has a distribution that lies between that of the two sources of real read datasets. Secondly, the Srinivas et al. dataset has very few reads of error rate less than 0.008 (Q=20.97). It was found in this work that adequate quality of estimated sequences from the denoising required that the error rate distribution be such that it was likely that each ASV was based on reads that included one read that had quality better than about 0.008. Note that the pre-filtered dataset from Srinivas et al. was markedly larger than from Sereika et al. but gave far fewer reads with error rate below 0.01. Because of the limited quality of the Srinivas et al. data no further analysis from that source is presented in this section, but is provided in the Supplementary Material Section 2. Our results from simulated reads (section 3.3.2, Figure 9) provide a brief examination of the effect of shifts in the distribution of mean read quality down towards (but not reaching) the quality of those Srinivas et al. data.

### 3.2 Identification Performance

Identification is based on achieving high quality alignment of ASVs to entries in the reference database. For rrn alignments this corresponds to having alignments of length approximately 4800 - 5000 bases and with fewer than 4 mis-matches plus indels. The 23S rRNA gene alignment lengths are approximately 2200 bases and have fewer than 3 mis-matches plus indels. Because some different strains have 23S rRNA gene sequences, and even rrn operon, that are identical or differ by a very small number of bases there may be tied best matches to operons in the database that result in unresolved strains or even species. Our simulated data allows some insight into such cases. That is not the case for the real read data as the mock microbiomes on which these were based differed at the genus level.

In our analysis of identification performance we use both visualisation and some simple quantification to indicate performance. RAD returns both the ASV sequence and the identities of the reads best associated with that ASV for all ASVs extracted. Visualisations of the ASVs estimated by denoising and the reads associated with these ASVs using 2-dimensional Uniform Manifold Approximation [28] was implemented through code from the R package, uwot [29]. Following alignment to the selected reference database using blastn [26], labeling was implemented that links ASVs to the identified species or strains. A simple quantification of identification comes from the number of genera, species or strains known to be in the microbiome that were not identified, together with the number of genera, species or strains that the identification process erroneously reports as present.

The identification process involves labeling an ASV as being from a particular operon in the database and that operon is one from a particular strain, species and genus. If the ASV label corresponds to one of the strains in the mock microbiome we refer to it as a valid label, otherwise it is an invalid label. An observation of a strain is labelled valid or invalid according to whether the strain was part of the mock microbiome or not. Strains that are in the mock microbiome of interest, but not identified by our analysis, are referred to as missed strains

#### 3.2.1 Sereika-derived Dataset

For the 23S data there were 23,613 fastq sequences before length and quality filtering, and of these 5,334 were processed by the RAD code. Figure 2 provides 2D UMAP visualisations of the relationships between ASVs and their associated reads for the 23S rRNA gene amplicons.

**Fig 2.**
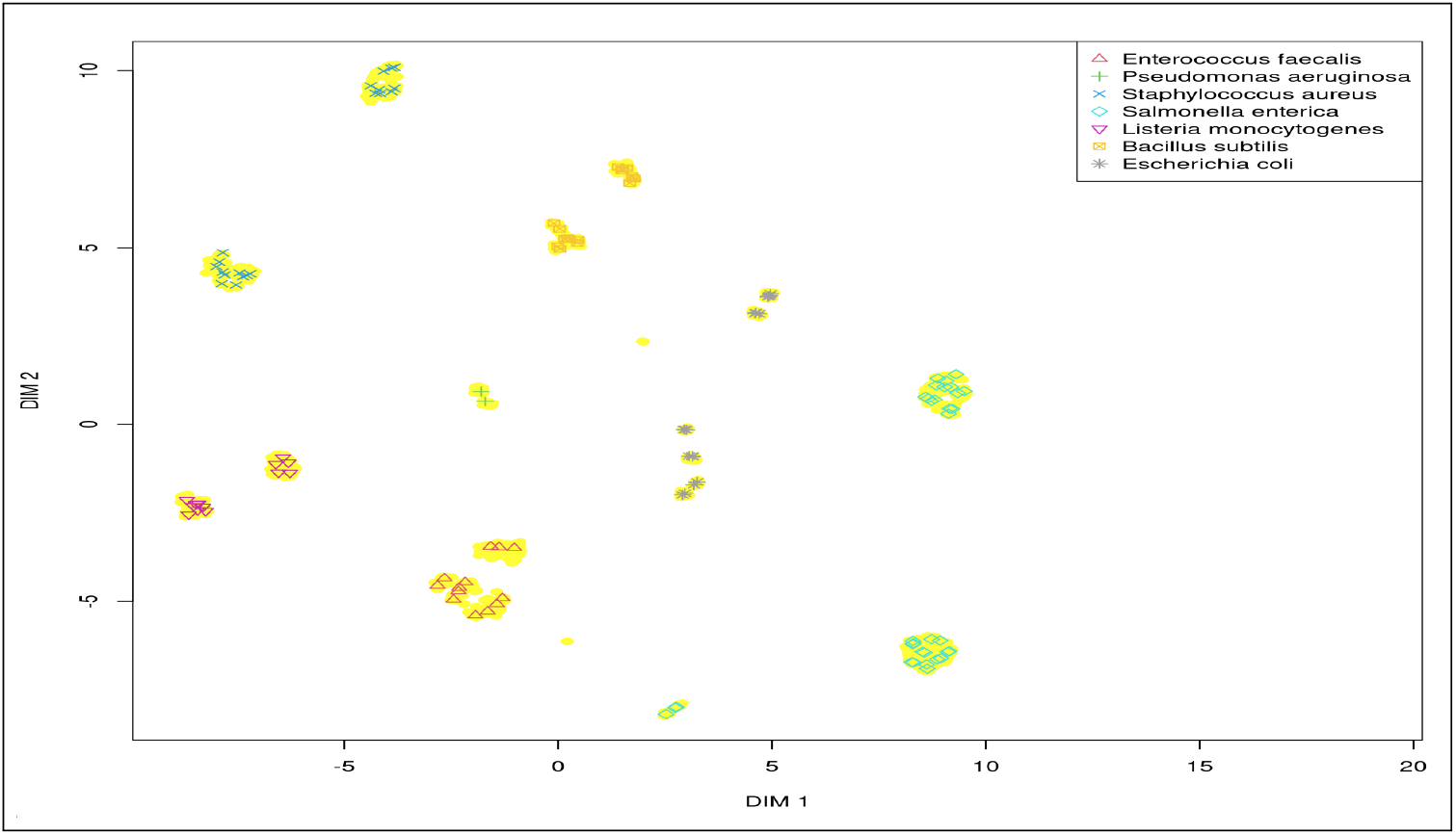
2D UMAP representation of the relationship between ASVs and their associated read clusters for the 23S rRNA gene sequences extracted from the Sereika et al. D6322 dataset.

This shows clear resolution of the species, and ASVs associated with most read clouds. Note that the default setting in RAD for ASV formation is that there be at least 5 reads close to the ASV of interest. Hence some small read clouds would not have any ASV associated with them. As noted earlier, the reference database has operons of each strain identified. Typically a strain has multiple operons, some of which are identical to each other. Also, reads may be reverse complements of one another and hence ASVs reflecting this arise. Hence it is not surprising that each species has multiple ASVs, particularly given the number and quality of reads.

The apparently clean resolution from the analysis on this Sereika-derived data allowed consideration of a sub-sampling process to investigate sensitivity over a larger dynamic range than the full dataset actually has. This was originally described in [30]. Relevant detail is provided in the Methods section, and in Supplementary Materials Section 1.2. Briefly, on the assumption that the reads associated with each ASV from the relevant rRNA gene are correctly identified, a sub-sampling of reads of any selected strain is possible. Four sets of such sub-samplings were selected (Table 1), and the reads from each of the sub-samplings constituted a library for a new mock microbiome that was then processed by the methods used for the full Sereika 16S and 23S datasets.

**Table 1.**
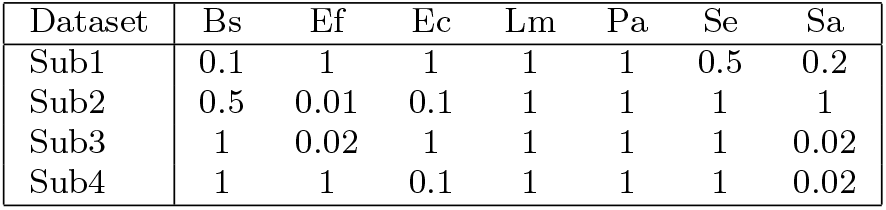
Sub-sampling factors on reads from species in the Sereika-derived 16S and 23S rRNA gene datasets used to generate four mock microbiomes that cover a range of either 50:1 or 100:1 in relative cellular abundance of the species. Species key: Bs=*Bacillus subtilis* , Ef=*Enterococcus faecalis* , Ec=*Escherichia coli*, Lm=*Listeria monocytogenes* , Pa= *Pseudomonas aeruginosa* , Se=*Salmonella enterica* , Sa=*Staphylococcus aureus* .

Figure 3 presents a UMAP representation of Sub3 from Table 1. It is seen that the two species, *Enteroccous faecalis* and *Staphylococcus aereus*, that are sub-sampled to 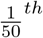 of the full dataset are represented by markedly fewer ASVs than in Figure 2. Despite the sub-sampling, species are still correctly identified and clearly resolved from other species.

**Fig 3.**
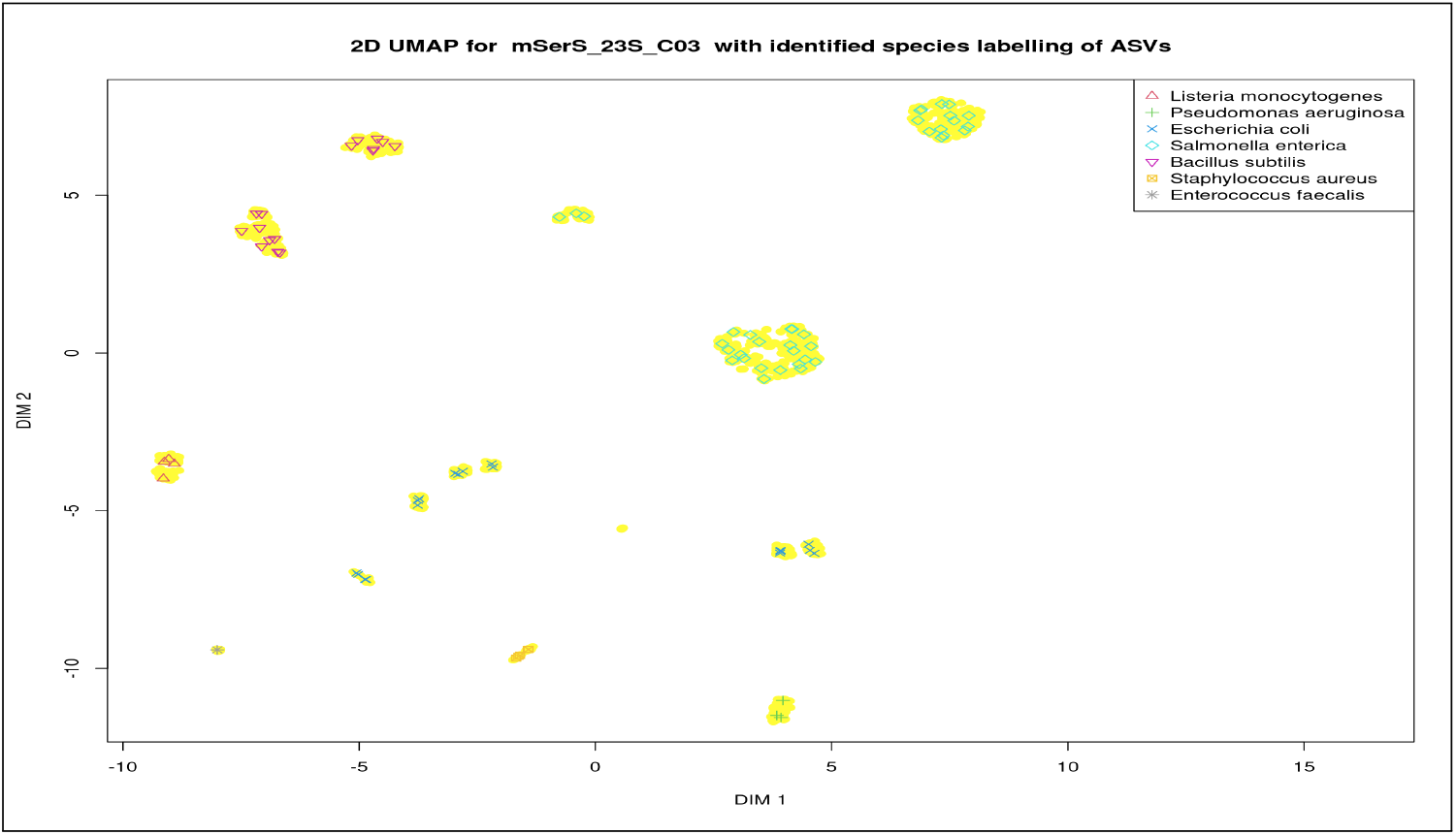
Visualisation of case Sub3 of the sub-sampling (Table 1) of the full Sereika et al.-derived 23S rRNA gene dataset.

#### 3.2.2 A realistic human gut mock microbiome, mKB_rrn_C11, with simulated nanopore-sequenced reads

Now consider a mock microbiome for which datasets of simulated 16S, 23S and rrn operon datasets were generated. We present data here only on the rrn operon dataset. This dataset is identified as `mKB_rrn_C11`. It has 59 strains with relative abundances matching the average values given by King et al. [24] (but re-calculated due to removal of the no-strain entries). Supplementary Materials Section 7 presents corresponding 16S and 23S rRNA gene datasets analysis.

Dataset `mKB_rrn_C11`has far more reads (98,762) than any of the real datasets considered above, and the mock microbiome has many more operons (291) than the earlier mock microbiomes (e.g.≈ 50). Not surprisingly, therefore, there are more ASVs formed-640. Figure 4 shows the extent of ASV coverage of the clusters of reads. This is in stark contrast to the results with the Srinivas data, and more akin to the simpler Sereika dataset.

**Fig 4.**
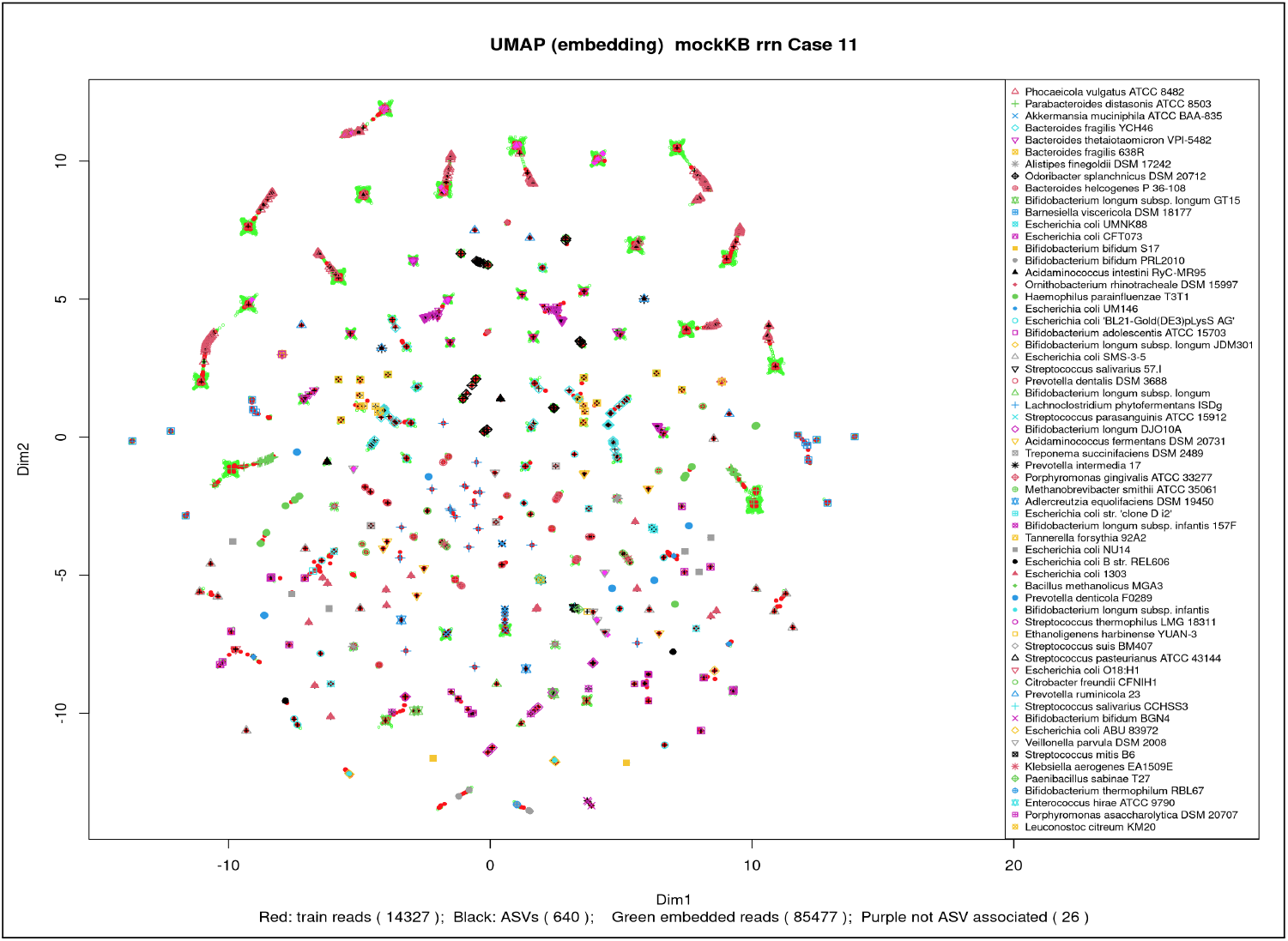
2D UMAP for `mKB_rrn_C11`dataset. The 2D UMAP projections computed for this figure involved training a model with a subset of the data followed by an embedding of the remainder of the data. Generated with R package uwot. ASVs are shown by characters according to the legend. Reads associated with a given ASV that are used in the training of the model are in red, embedded reads in green.

What is clear from this figure is that there is no strong separation of strains from each other. However there is strong localisation of the reads associated with each ASV to that ASV. ASVs are themselves aligned to operons, and most of these alignments are for close to the full length of the operon, and overwhelmingly have 100% identity of reference operon sequence with ASV sequence. Our reference database names each entry with the (genus, species, strain, operon index), and so our ASV alignments are usually highly specific to a particular operon, and that specificity is reliable.

In Figure 5 UMAP 2D projection plots for four species with multiple strains are displayed. The mapping is computed independently for the 4 species. The *Bacillus fragilis* plot has 2 strains both of which are in the design. The *Escherichia coli* plot shows 11 strains, but 6 of those are not in the actual mock microbiome. One strain that was in the design was not identified. The *Bifidobacterium longum* plot shows 6 entries, 2 of which are not true strain entries. Excluding those 2, the 4 strain entries are definitely in the design and the design does not have further strains of this species. Finally, there are 3 *Bifidobacterium bifidum* strains each of which is in the design, with no strains missed. This data, viewed from the operon level, strongly supports strain-level resolution in identification.

**Fig 5.**
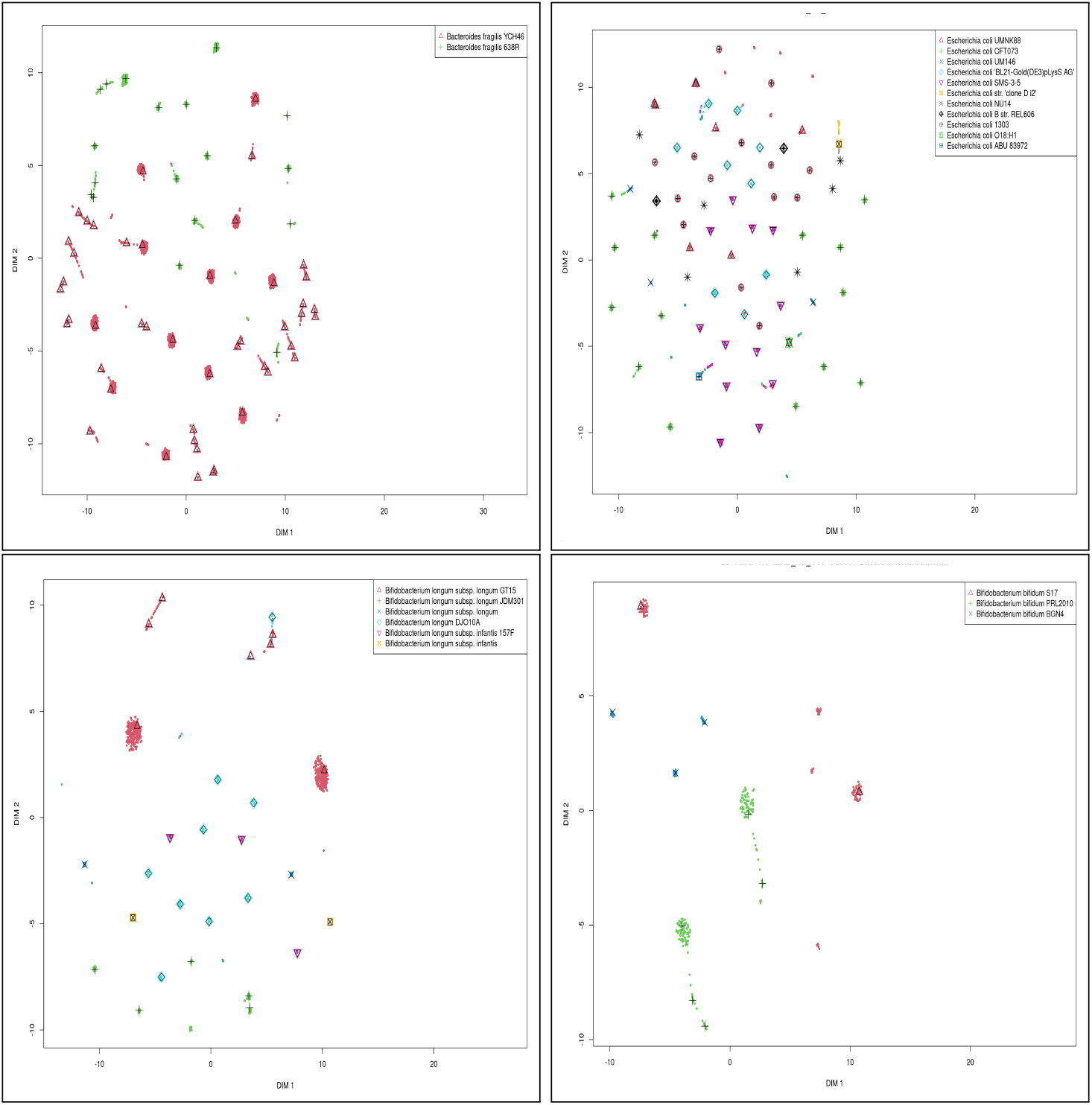
2D UMAP representation of the relationship between ASVs and their associated read clusters for the simulated rrn sequences for strain-specific subsets of the full KB dataset of Figure 4. Both *Bacteroides fragilis* strains are valid and none are missed. For *Escherichia coli* 5 of 11 strains are valid, 6 invalid, 1 missed. For *Bifidobacterium longum* 4 strains are valid, none are missed. For *Bifidobacterium bifidum* all 3 strains are valid, none are missed. ASVs are shown as characters, while reads associated with an ASV are filled circles of colour corresponding to the legend colour for that ASV.

In Supplementary Material Sections 4 and 5 a more detailed analysis for these four species is provided that looks at all optimal alignments for all ASVs of each of these species. That includes a discussion of how the operon data - both its completeness and the matching or otherwise of indices - can assist in distinguishing highly similar strains (Supplementary Materials Section 5). This assumes that operon indices for any strain are ordered consistently with location along the genome of each operon. For the *Eshcerichia coli* species this analysis led to the following being the identified strains - CFT073, SMS-3-5, 1303, UMNK88, one of (B, BL21, BL21(DEL)), one of (NU14, UTI89). The actual strains present are the same first 4 listed, together with BL21(DE3) and UTI89.

### 3.3 Quantification Performance

Quantification performance is indicated by the extent of agreement between the relative abundance (RA) of species (or genera, or strains) in the known microbiome (the design RA) and the corresponding values of relative abundance determined by analysis of the read datasets (the observed RA). This can be indicated in scatterplots and in correlation coefficients and regression line slopes.

#### 3.3.1 The Sereika-derived 23S rRNA gene datasets

Figure 6 provides species scatterplots for the 23 rRNA gene analysis of the real sequenced data from the Sereika et al. full microbiome and subsample Sub3. The accuracy is not high, but this microbiome only has a small dynamic range and there may be errors arising prior to sequencing that distort the results.

**Fig 6.**
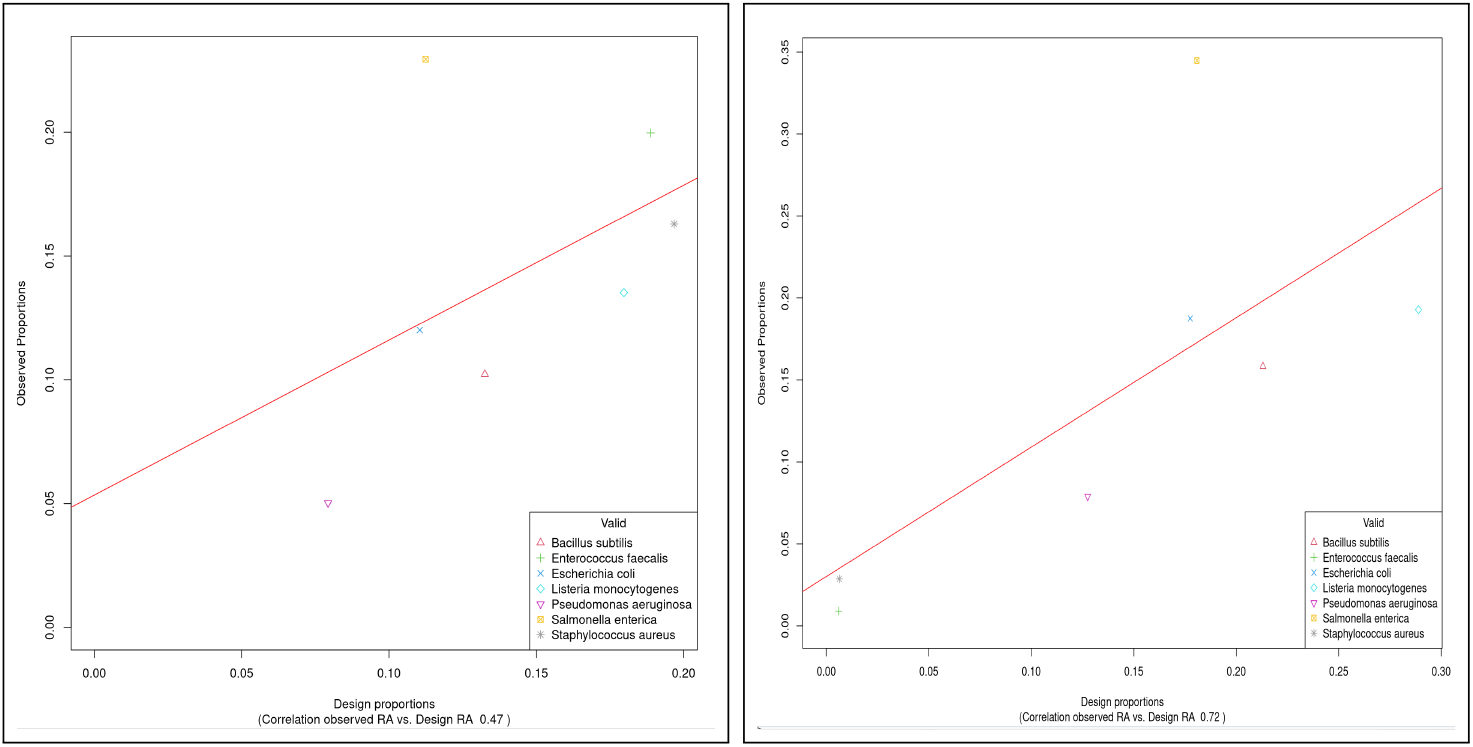
Relation between the relative abundances of the observed species and species of the designed microbiome as determined from 23S rRNA genes. The full Sereika et al.-derived dataset is on the left, while sub-sample Sub3 is on the right. Note that the order of observed relative abundances flips for the two species being down-sampled between the full and the sub-sampled datasets, indicating some inaccuracy in the observed values.

The observed increase in correlation between the full and sub-sampled datasets as the RA in the designed microbiome covers a greater range is what one would expect if the quantification is moderately accurate. This result is supported by the results from the other sub-samples (Supplementary Material). Note that the RA range here is not 50:1 but a bit less, as the two species with highest RA in the full dataset were the ones sub-sampled in Sub3 at 50:1.

#### 3.3.2 The mKB_rrn_C11dataset

A more substantive and informative analysis is now provided based on the simulated reads mock microbiome, `mKB_rrn_C11`. Figure 7 presents log plots of observed cellular abundances vs. designed relative cellular abundances at both strain and species level of taxonomic resolution. This ranges over nearly 4 orders of magnitude in relative abundance, and the observed data is very accurate as indicated by the Pearson correlation value of greater than 0.995 over this range of the observed abundances and design abundances. Note that the design abundances vary over a range of approximately 4500:1. The missed and invalid strains all have abundances that are less than about 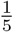 of the most abundant strain.

**Fig 7.**
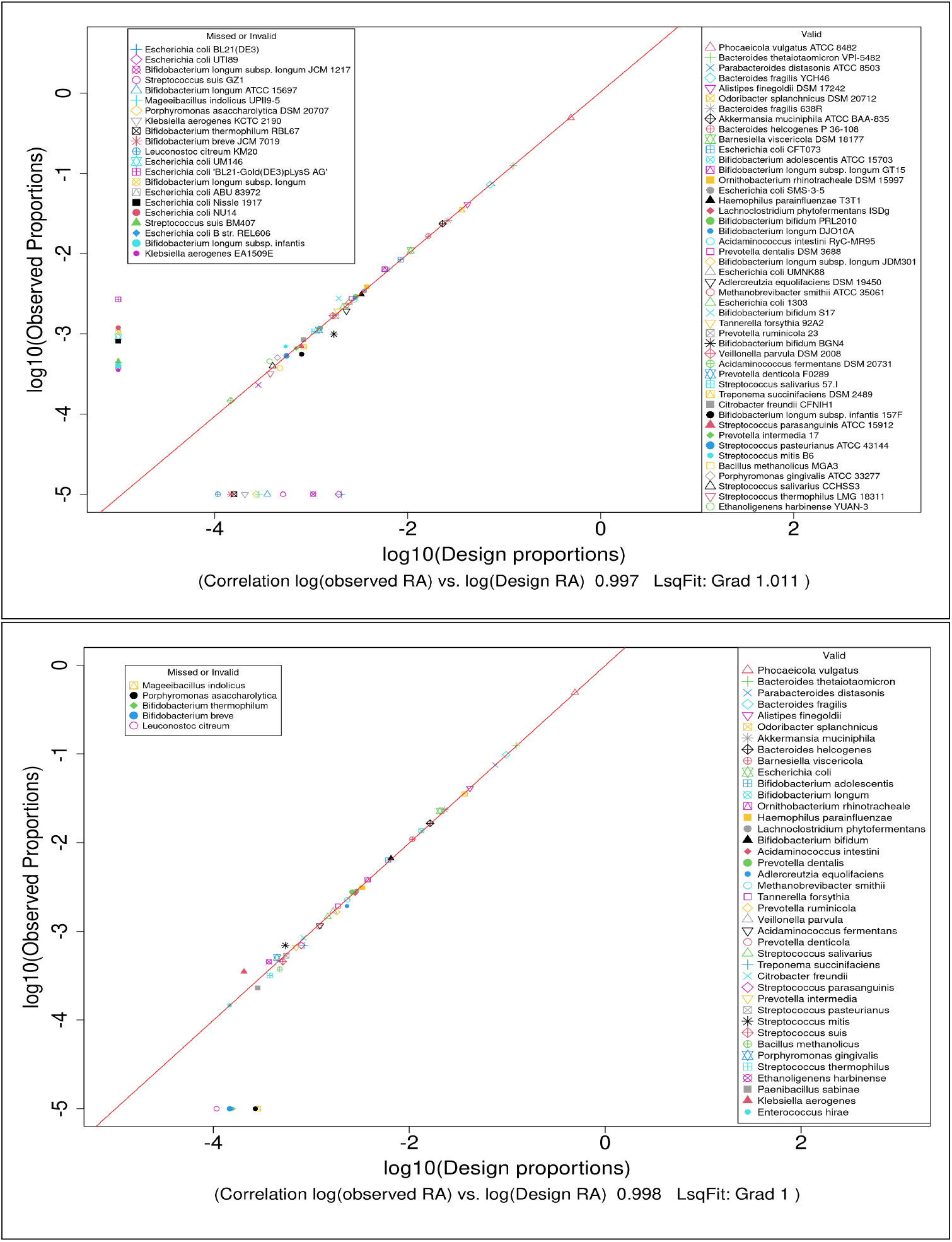
Relation between the (log_10_) observed relative abundances of strains (top) and species (bottom) and that of the designed microbiome as determined from badread-simulated full rrn sequences. Missed strains are plotted along the -5 Y-coordinate, invalid strains on the -5 X-coordinate. Correlation coefficient calculations do not include missed or invalid data.

Figure 8 shows the shift in read mean error rate, and in the number of reads available for denoising, as the simulated reads quality distribution has its mean shifted from *Q* = 28 to *Q* = 22. It also shows the clear gain with strain-level resolution (left hand sub-figure) from using the rrn compared to the shorter 23S and 16S rRNA genes. As the taxonomic level of resolution sought decreases (left to right sub-figures) it is seen that the 16S becomes progressively more effective. It is worth noting that the 16S rRNA gene is better for genus-level than is indicated for the 23S rRNA gene and, possibly,for the rrn operon datasets.

**Fig 8.**
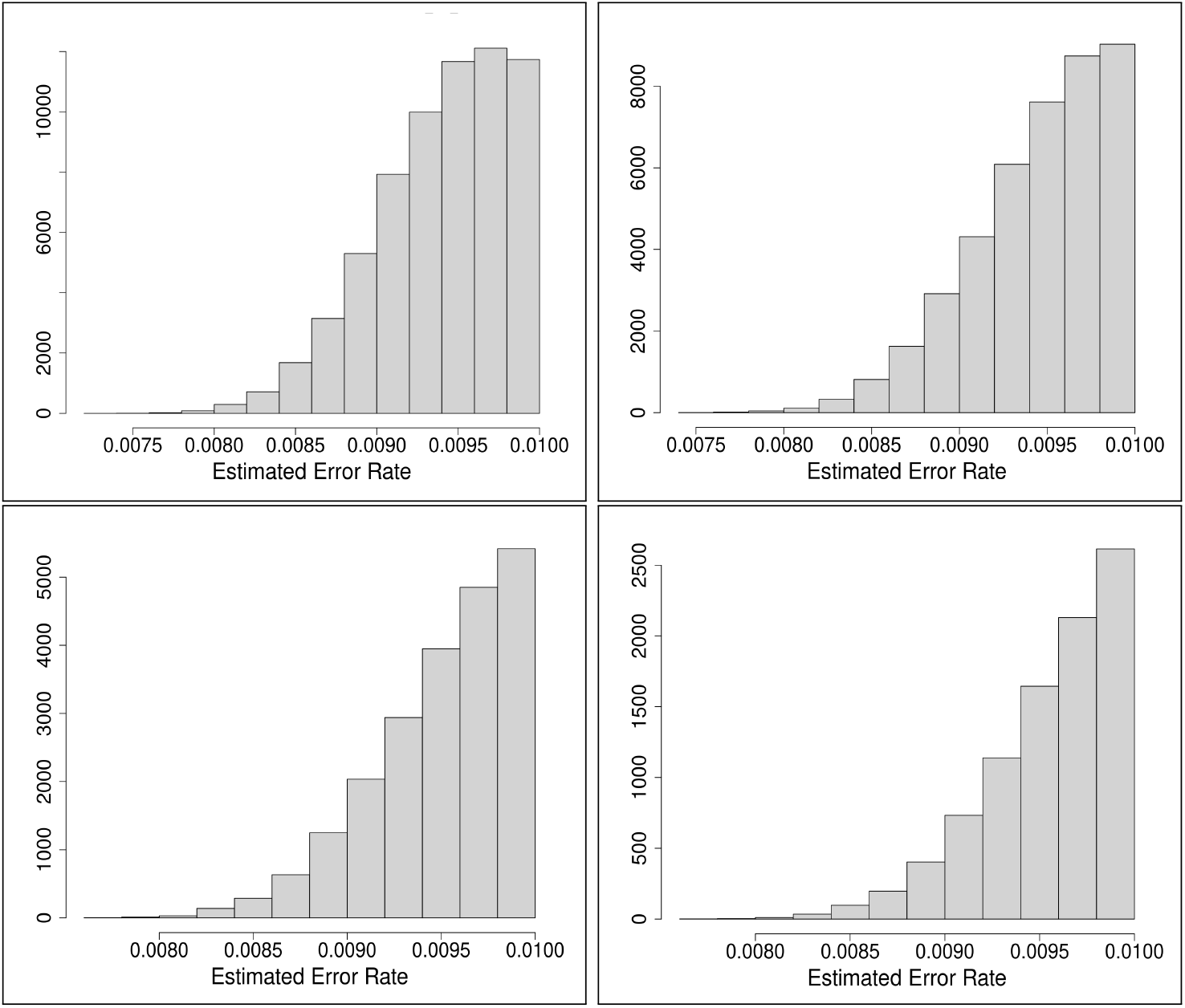
Error rate distributions indicative of datasets with mean Q of 28, 26, 24 and 22 from top left to bottom right row-wise.

**Fig 9.**
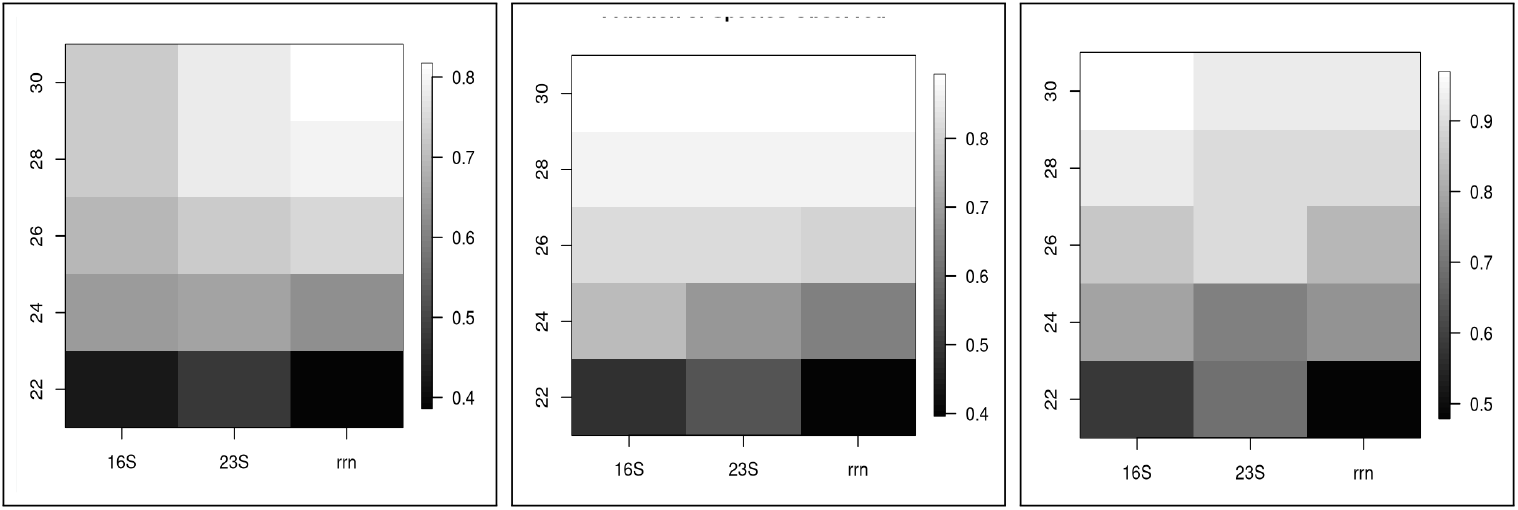
Fraction of strains (left), species(centre) or genera (right) present in the mock microbiome with datasets from simulated reads that were actually observed. Each plot shows 5 different mean quality score datasets for each of 16S and 23S rRNA genes and the rrn operon.

Figure 9 shows the marked improvement in valid strain labelling from Q=22 to Q=30, and - at strain and species level, but not genus level - from using either the 16S or 23S rRNA genes compared to the full rrn operon.

## 4 Discussion

These results, built on the excellent denoising of sequences ∼5000 bases long by the Kumar et al. RAD code, clearly demonstrate that accurate identification at operon-level of the strains present in complex bacterial microbiomes is now possible with ONT nanopore sequencing. Our results also show that quantification is possible over at least 4 orders of magnitude in relative abundance, subject to sequencing of adequate depth. These results, of course, are reliant on an adequate operon-level reference database, and there is more work needed to generate this infrastructure. We have highlighted that such databases should be derived from complete genome sequences, and have operons indexed according to their order along the genome.

Because by far the majority of ASVs gave excellent alignments with database operons over all, or almost all, the length of those operons, the performance achieved would be close to what would be achieved with state-of-the-art PacBio consensus sequencing. The analysis methods here are directly applicable to such consensus sequences, simply by treating these consensus sequences as ASVs. It would be possible, if necessary, to apply denoising to a library of consensus sequences. The ONT nanopore sequencer has application advantages over PacBio technology of field deployability, as well as capital and running costs. The PacBio technology, however, would be the preferred technology to underpin populating of the appropriate databases.

The use of alignment of denoised sequences provides a considerable reduction in computational demand, as there are far fewer ASVs than there are reads in most datasets. This also reduces downstream computational demands. We have used the default lower threshold, 5, in the RAD denoising code for ASV formation. As a result it can happen that some reads are either lost or inappropriately assigned to ASVs. Our Supplementary Material shows how these can be identified and some useful additional identification and more accurate quantification obtained. The effect is small, and would not normally be worth the effort.

The reliance on simulated nanopore sequenced reads for the major results is a limitation. However, the choice of Wick’s badread simulator was made following experience with a range of simulators. Badread incorporates a large empirical base of kmer probabilities of correctness, or of any one of many possible errors, given the context of each base. It also considers the effect of insertions and deletions in a substantial way. Related methods - [31], [32] - seek to generate the actual electric signal. While it is not difficult to move from signal to fastq file libraries, it was found that delivering simulated reads at the quality sought in this work was apparently not yet possible. Thus we almost certainly have used the best publicly available simulator for this study.

It is recognised that the rrn operon is itself not adequate for some strain resolution requirements, including some important discriminations that need to be made in commercial probiotics. Multi-locus studies based on NGS sequencing such as [33] are addressing that. It may be that the methods developed here could be usefully applied to such studies, particularly as they seek to use information on genes whose sequences are much longer than can be processed with NGS sequencing.

## 5 Methods

This section addresses four key steps in implementing our method of analysis - micro-biome specification and characterisation, denoising, reference database generation, and analysis of the results from denoising.

### 5.1 Specification and Characterisation of Microbiomes Analysed

Two sources of real nanopore-sequenced microbiome genomic DNA - Sereika et al. [22] and Srinivas et al. [23] - and one source of simulated microbiome data based on a proposed standard healthy human gut microbiome, King et al. [24], were used.

#### 5.1.1 Real Sequenced Data

The Sereika et al. data was from the Zymo Research D6322 microbiome standard [34] which has equal masses of genomic DNA of 7 bacterial organisms, and 2% yeast, *Saccharomyces cerevisiae* . Whole genome sequencing of this “even” mock microbiome by Sereika et al. [22] with Oxford Nanotechnologies (ONT) R10.4.1 pores on a Prome-thION sequencer resulted in 52.3GB of reads with an N50 of 5,992 bases and estimated modal read accuracy of 98.13%, based on Q-score for reads, and consisted of 22.3 million reads. It was estimated that a useful number of 16S and 23S rRNA gene sequences could be extracted from this dataset to allow an analysis of the feasibility of subspecies resolution of microbiomes. The computationally expensive extraction process is described in Supplementary Materials S1. Two datasets were obtained - one of the 16S ribosomal gene, the other the 23S ribosomal gene. These are identified as mSerF-16S, and mSerF-23S respectively, where F (Full) indicates there has been no sub-sampling (see Supplementary Materials 1). To extend the analysis of these real sequenced reads a sub-sampling process, Table 1, was used to give insight into performance of the analysis methods for more realistic relative abundance structures of microbiomes (noting that the D6322 mock microbiome is an “even” distribution of genomic DNA mass). These four sub-sampled datasets were identified as mSerSn, n=1,2,3,4.

The Srinivas et al. data was from two mock microbiomes. One was the Zymo Research D6311 microbiome standard [35] that has 8 bacterial strains, with a logarithmic distribution of relative abundance of genomic DNA mass over 7 orders of magnitude (0.89 to 0.0000009), deliberately contaminated with 2% of a yeast. The other was an ATCC mock microbiome of 10 microbial strains, with equal masses of genomic DNA for each strain. Further details are provided in Supplementary Materials Section 1.2.

#### 5.1.2 Datasets from Simulated Reads of a mock gut microbiome

Merging information from King et al’s Gut Feeling Knowledge Base (GFKB) [24] and the GROND-derived strain-level blast database allowed selection of 59 strains common to both. From [24] the rank order of average relative abundance in the proposed standard health human gut was also available. We specified a mock microbiome consisting of all 59 strains and with the relative abundances of these strains as reported by King et al, scaled to add to 1, giving a range of approximately 5000:1. The rrn operon library of reads for this microbiome is labelled `mKB_rrn_C11`. Correspondingly labelled 16S and 23S rRNA gene versions were also generated. Datasets of 16S and 23S rRNA gene and rrn operon sequences were constructed, with simulated ONT nanopore-sequenced reads generated for two different read quality settings. The simulator was Wick’s badread simulator [25]. In the case of `mKB_rrn_C11`reads were simulated for each of 291 operons separately, in numbers larger than would be used, and then randomly sampled for each operon to give the required composition in the dataset, and likewise for the 16S and 23S rRNA gene datasets.

Two reference databases were used, both based on the Walsh et al. rRNA data base, GROND Refseq [27]. Because our study was focused on sub-species profiling of microbiota and the healthy human gut microbiome, a strains subset of the GROND Refseq database was formed. This subset consisted of all ribosomal DNA (rDNA) sequences in the GROND Refseq entries that were from complete genomes and that identified the strain from which the sequence was derived. There were 25,285 such entries. All entries had their GROND sequence name modified to specify the genus, species and strain (if available) from which the sequence was derived, together with an operon index. For many strains this was realistic in that the strain had the correct number of different indices. In other cases there were clearly multiple sub-strains (or repeats of the same strain) contiguous in the GROND database - e.g. Escherichia coli K-12 strain had hundreds of contiguous entries. For the Srinivas data, where the focus had been on resolution to the species level, a second, larger variant of the GROND database was created containing all GROND Refseq entries that had a complete genome. This database had 147,291 entries.

### 5.2 Denoising

Denoising in this context involves processing a relatively large number of noisy reads that originated from a markedly smaller number of original sequences to estimate what those original sequences were. We used the Robust Amplicon Denoising (RAD) algorithm of Kumar et al. [21], executed in the Julia code from the authors with the default parameters. The input is a single fastq file with all the reads of the relevant dataset. This process returned 3 key types of information, i.the set of estimated underlying sequences - which we refer to as Amplicon Sequences Variants (ASVs) - for each ASV, ii. the indices into the input fastq sequence set of the reads associated with that ASV (Note that each read is associated with only 1 ASV); iii. the new fastq sequence headers of the sequences that passed length and quality filtering that were actually used by the denoising process. Each new header gives the mean error rate for the retained fastq sequence, together with an index to the fastq record in the input fastq file.

### 5.3 Database Generation

NCBI’s blastn was used for alignment of query sequences against our databases. This was run locally with two bespoke databases, derived from Walsh et al’s GROND Refseq database [27]. The key modification we used was to incorporate directly informative names into the sequences from which the databases were built. By informative is meant that the sequence names in the fasta file from which the database is built incorporated genus, species, strain and operon information.

The two GROND ribosomal RNA sequence databases consist of sequences extracted from genome sequences in Refseq (grondR), the other covering sequences in GenBank (grondG). While Genbank has a larger coverage than Refseq it is doubtful that metagenome-assembled genomes - which are now part of Genbank - have reliable differentiation of different *rrn* operons within any strain. For this work the focus was on sub-species taxonomic resolution and relative abundance quantification at strain, species and genus level. This requires strain entries that have sequences of all operons of the particular strain within the database. The GROND Refseq database partitions the entries by name to identify those from complete genomes and those from incomplete genomes. The Refseq data is more fully curated than Genbank and hence tends to be more accurate. Therefore we chose to use the Refseq data derived from complete genomes.

An alternative database that was considered was the rOPDB database [36]. This was derived from the same sources as the GROND databases but was less suitable for several reasons:-

- It provided no differentiation between entries derived from Genbank and those from Refseq,
- It did not allow identification of entries based solely on complete genomes,
- It did not have some entries which are significant in the healthy human gut (e.g. *Phocaeicola vulgatus*). For others, such as *Bacteroides thetaiotaomicron VPI 582, Parabacteroides distasonis ATCC 8053, Bacteroides fragilis* strains YCH46, 638R and NC9343, and *Alistipes finegoldii DSM17242* - amongst the highest relative abundances in King et al’s reference human gut microbiome - the number of entries was fewer (in multiple cases only a single entry when there are multiple operons) than what is recorded in Refseq listings.

Hence the rOPDB was not fit for the purpose of this study.

Two versions of the GROND-based database were constructed. The first consisted of all the grondR entries that had both strain-level data according to the taxonomic listing for the Refseq207full entries, and also were from complete genomes. The result was a database with 25,285 rRNA operon sequences - not all unique - from 1,474 species. The second database consisted of all Refseq207 sequences from complete genomes, without any restriction on whether they had strain-level information. This had 147,291 entries for 7,345 species entries and 3,269 strain entries.

### 5.4 Analysis of the results from denoising

This involves alignment of the ASVs against the reference database followed by interpretation of the results from alignment in terms of microbiota identification and profiling for the microbiome.

#### 5.4.1 Alignment of ASVs to the Reference Database

NCBI’s blastn v.2.16.0+ was used for alignment of query sequences against our databases. The primary alignment work involved alignment of ASVs against the reference database. Cross-referencing to the taxonomy file of GROND then allows determination of the best-matching database entry (or entries). The taxonomy files do not provide information below the strain level in many cases - for example, there are hundreds of entries for *Escherichia coli K-12* but any such strain’s genome will only have (typically) 7 operons. The operons themselves are often not identical, but there is no facility to distinguish between them. The maximal amount of lower level (genus, species, strain) information, as well as operon indexing, was incorporated into the names of sequences used to create the blastn reference databases being used. Then, ensuring naming of reads generated by simulation is consistent with the template sequences from which such reads are generated, sub-strain, and operon-level discrimination is enabled.

NCBI’s blastn code was run locally using the databases described above, which were constructed from a fasta file of reference sequences using NCBI’s makeblastdb code. As mentioned earlier, the sequence names were constructed to ensure a common format for genus and species names, to include strain names if available, and to have an operon index attached. They also needed to be no more than 50 characters long, which often required trimming of the strain name detail . Textual output is controlled by multiple options in the call of blastn, and this was tailored to give multiple quality scores on either 10 or - for diagnostic purposes - 50 alignments.

#### 5.4.2 Primary processing of alignment data

For a single ASV being aligned to the database the set of alignments returned is ordered by a quality measure labelled the bitscore. Alignments of equal bitscore are ordered lexicographically based on the name of the operon entry in the database. Length of the alignment is also returned and we have used this to remove short alignments (e.g. less than 4000 for rrn operons). It is common for multiple alignments of a single ASV to give identical best quality measures. Such cases are resolved in our primary analysis by simply taking the first listed, which happens to be alphabetically sorted - e.g. both *Bacillus intestinalis* and *Bacillus subtilis* are sometimes equal best alignments for an ASV, but our method only returns *Bacillus intestinalis* . This obviously can introduce systematic errors, though our experience is that by far the majority of alignments are unambiguous at the strain level - but would, of course, change with much more comprehensive reference databases. In Supplementary Material section 5 we provide a more detailed method for dealing with tied optimal alignments that better resolves that ambiguity.

#### 5.4.3 Identification and Quantification

Having linked each ASV to a single operon, we now identify all ASVs linked to each operon. Summing the read counts for the ASVs linked to an operon provides a read count for each operon. This allows all identified operons to be given a relative abundance.

The strain, species, and genus to which an operon belongs is part of the identity of that operon. By condensing operon profiling across strain, and knowing how many operons that strain has, we can convert operon read counts for a strain to estimates of strain cellular counts and hence obtain strain relative abundances for the microbiome. Further condensation gives species and genus level resolution data.

## Acknowledgments

Valuable support in using the RAD denoising code and the visualisation code, seqUMAP, was provided by Benjamin Murrell, Department of Microbiology, Tumor and Cell Biology, Karolinska Institutet, Sweden.

Advice and support from the WEHI IT and Research Computing personnel to the first author has been greatly appreciated.

## Declarations

- CJW acknowledges the provision of computing resources by the Walter and Eliza Hall Institute.
- The authors declare there are no competing interests.
- The authors consent to publication.
- Code to run all 21 datasets for which analysis results are presented in this paper, and its Supplementary Materials, is available at [37]. Supplementary Materials section 8 provides detail on the pipeline and its code elements.
- CJW conceived the project, developed methods and code, and wrote the draft paper. TPS provided ongoing critical review throughout the project, access to computing and literature resources, and reviewed written material. Both authors finalised the paper.

## 6 Data Availability

Real nanopore-sequenced data came from [22] and [23]. The 8 datasets from [23] had been part of a European Nucleotide Archive (ENA) deposition with project accession number PRJEB7254. The 10 datasets derived from [22] were from an ENA deposition with accession number PRJEB 48692. All 18 of these datasets are deposited in Figshare, [38]. Also deposited in Figshare at [39] are various files that allow the analyses presented here to be conveniently replicated or extended.

## References

[1] Ling-chao Ma et al. “Impact of the microbiome on human, animal, and environmental health from a One Health perspective”. In: Science in One Health (2023). doi: 10.1016/j.soh.2023.100037. url: https://doi.org/10.1016/j.soh.2023.100037.

[2] M. Nazmul Huda, Myungsuk Kim, and Brian J. Bennett. “Modulating the Microbiota as a Therapeutic Intervention for Type 2 Diabetes”. In: Frontiers in Endocrinology 12 (Apr. 2021). doi: 10.3389/fendo.2021.632335.

[3] F. Indrio and F. Marchese. “Microbiota in the first 1000 days: Effect on health and diseases”. In: Global Pediatrics 8 (Feb. 2024), p. 100162. doi: 10.1016/j.gpeds.2024.100162. url: https://doi.org/10.1016/j.gpeds.2024.100162.

[4] Jun Chen, Xinyi Chen, and Chun Loong Ho. “Recent Development of Probiotic Bifidobacteria for Treating Human Diseases: Systematic review of gut microbiota and major depression”. In: Frontiers in Bioengineering and Biotechnology 9 (Dec. 2021), p. 770248. doi: 10.3389/fbioe.2021.770248.

[5] K Arunachalam, HS Gill, and RK Chandra. “Enhancement of natural immune function by dietary consumption of Bifidobacterium lactis (HN019)”. In: European Journal of Clinical Nutrition 54.3 (Mar. 2000), pp. 263–267. doi: 10.1038/sj.ejcn.1600938.

[6] Alberto Finamore et al. “Supplementation with Bifidobacterium longum Bar33 and Lactobacillus helveticus Bar13 mixture improves immunity in elderly humans (over 75 years) and aged mice”. In: Nutrition 63-64 (July 2019), pp. 184– 192. doi: 10.1016/j.nut.2019.02.005. url: https://doi.org/10.1016/j.nut.2019.02.005.

[7] Emily R. Murray, Mylon Kemp, and Tanya T. Nguyen. “The Microbiota–Gut–Brain Axis in Alzheimer’s Disease: A Review of Taxonomic Alterations and Potential Avenues for Interventions”. In: Archives of Clinical Neuropsychology 37 (2022), pp. 595–607. doi: 10.1093/arclin/acac008.

[8] A. Kolobaric et al. “Gut microbiome predicts cognitive function and depressive symptoms in late life”. In: Nature Molecular Psychiatry 29 (Apr. 2024), pp. 3064–3075. doi: 10.1101/0155522307/2308930.

[9] Kate L Tooley. “Effects of the human gut microbiota on cognitive performance, brain structure and function : a narrative review”. In: Nutrients 12.10 (Sept. 2020), p. 3009. doi: 10.3390/nu12103009.

[10] Benjamin D. Anderson and Jordan E. Bisanz. “Challenges and opportunities of strain diversity in gut microbiome research”. In: Frontiers in Microbiology 14 (Feb. 2023), p. 1117122. doi: 10.3389/fmicb.2023.1117122.

[11] Mariarosario Matera. “Bidfidobacteria , Lactobacilli … when, how and why to use them”. In: Global Pediatrics 8 (Jan. 2024), p. 100139. doi: 10.1016/j.gpeds.2024.100139.

[12] Thomas Hoenen et al. “Nanopore Sequencing as a Rapidly Deployable Ebola Outbreak Tool”. In: Emerging Infectious Diseases 22.2 (Feb. 2016), pp. 331– 334. doi: 10.3201/eid2202.151796.

[13] Tiana C. Schwab et al. “Field evaluation of nanopore targeted next-generation sequencing to predict drug-resistant tuberculosis from native sputum in South Africa and Zambia”. In: Journal of Clinical Microbiology 63.3 (Mar. 2025), pp. 1–14. doi: 10.1128/jcm.01390-24.

[14] Sarah L. Castro-Wallace et al. “Nanopore DNA Sequencing and Genome Assembly on the International Space Station”. In: Scientific Reports 7.18022 (Dec. 2017). doi: 10.1038/s41598-017-18364-0.

[15] Christopher E. Carr et al. “Nanopore sequencing at Mars, Europa, and microgravity conditions”. In: npj Microgravity 6.24 (2020). doi: 10.1038/s41526-020-00113-9.

[16] Benitéz-Páez Kevin J Portune, and Yolanda Sanz. “Species-level resolution of 16S rRNA gene amplicons sequenced through the MinION™ portable nanopore sequencer”. In: GigaScience 5.4 (2016), pp. 1–9. doi: 10.1186/s13742-016-0111-z.

[17] Benitéz-Páez and Yolanda Sanz. “Multi-locus and long amplicon sequencing approach to study microbial diversity at species level using the MinION™ portable nanopore sequencer”. In: GigaScience 6 (June 2017), pp. 1– 12. doi: 10.1093/gigascience/gix043.

[18] Anna Cuscó et al. “Using MinION™ to characterize dog skin microbiota through full-length 16S rRNA gene sequencing approach”. In: GigaScience 6 (July 2017), pp. 1–10. doi: 10.1093/gigascience/gix007.

[19] Anna Cuscó et al. “Microbiota profiling with long amplicons using Nanopore sequencing: full-length 16S rRNA gene and the 16S-ITS-23S of the rrn operon”. In: F1000 Research 7 (Aug. 2019), pp. 1755–1781. doi: 10.12688/f1000research.16817.2.

[20] Lee J. Kerkhof et al. “Profiling bacterial communities by MinION sequencing of ribosomal operons”. In: Microbiome 5.116 (Sept. 2017). doi: 10.1186/s40168-017-0336-9.

[21] Venkatesh Kumar et al. “Long-read amplicon denoising”. In: Nucleic Acids Research 47.18 (Aug. 2019), e104. url: 10.1093/nar/gkz657.

[22] Mantas Sereika et al. “Oxford Nanopore R10.4 long-read sequencing enables the generation of near-finished bacterial genomes from pure cultures and metagenomes without short-read or reference polishing”. In: Nature Methods 19 (July 2022), pp. 823–826. doi: 10.1038/s41592-022-01539-7.

[23] Meghana Srinivas et al. “Evaluating the efficiency of 16S-ITS-23S operon sequencing for species level resolution in microbial communities”. In: Scientific Reports 15 (Jan. 2025), p. 2822. doi: 10.1038/s41598-024-8310-7.

[24] Charles H. King et al. “Baseline human gut microbiota profile in healthy people and standard reporting template”. In: PLoS One 14.9 (Sept. 2019), e0206484. doi: 10.1371/journal.pone.0206484.

[25] Ryan R. Wick. “Badread: simulation of error-prone long reads”. In: Journal of Open Source Software 4.36 (2019), p. 1316. doi: 10.21105/joss.01316.

[26] C Camacho et al. “BLAST+: architecture and applications”. In: BMC Bioinformatics 10 (2009), pp. 421–429. doi: 10.1186/1471-2105-10-421.

[27] Calum J. Walsh et al. “GROND: a quality-checked and publicly available database of full-length 16S-ITS-23S rRNA operon sequences”. In: Microbial Genomics 10.001255 (June 2024). doi: 10.1099/mgen.0.001255.

[28] Leland McInnes et al. “UMAP: Uniform Manifold Approximation and Projection”. In: Journal of Open Source Software 3.29 (Sept. 2018), pp. 861–862. doi: 10.21105/joss.00861.

[29] James Melville. uwot: The Uniform Manifold Approximation and Projection (UMAP) Method for Dimensionality Reduction. R package version 0.2.3. 2025. doi: 10.32614/CRAN.package.uwot. url: https://CRAN.R-project.org/package=uwot.

[30] Christopher J. Woodruff, Zhengming Zhang, and Terence P. Speed. “Full 16S and 23S rRNA gene-based,strain-level resolution of the microbiota of a mock bacterial microbiome using ONT nanopore sequencing”. In: bioRxiv (2024). doi: 10.1101/2024.02.07.579414. eprint: https://www.biorxiv.org/content/early/2024/02/09/2024.02.07.579414.full.pdf.

[31] Hasindu Gamaarachchi et al. “Simulation of nanopore sequencing signal data with tunable parameters”. In: Genome Research 34 (May 202), pp. 778–783. doi: 10.1101/gr.278730.123. url: https://www.genome.org/cgi/doi/10.1101/gr.278730.123.

[32] Denis Beslic et al. “End-to-end simulation of nanopore sequencing signals with feed-forward transformers”. In: Bioinformatics 41.1 (Jan. 2025). url: 10.1093/bioinformatics/btae744.

[33] Stephen Nayfach et al. “An integrated metagenomics pipeline for strain profiling reveals novel patterns of transmission and global biogeography of bacteria”. In: Genome Research 26 (Oct. 2016), pp. 1–14. doi: 10.1101/gr.201863.115.

[34] Zymobiomics HMW DNA Standard D6322. (Accessed 25 January 2026). 2026. url: https://zymoresearch.eu/products/zymobiomics-hmw-dna-standard.

[35] Zymobiomics HMW DNA Standard D6311. (Accessed 25 January 2026). 2026. url: https://zymoresearch.eu/products/zymobiomics-microbial-community-dna-standard-ii-log-distribution.

[36] Lee J. Kerkhof et al. “A ribosomal operon database and MegaBLAST settings for strain-level resolution of microbiomes”. In: FEMS Microbes 3 (Jan. 2022), pp. 1–9. doi: DOI:10.1093/femsmc/xtac002.

[37] Chris Woodruff. ASVpaper Rcode and processing pipeline scripts. (Accessed 26 May 2026). 2026. url: https://github.com/cjwoodruff50/ASVpaper.

[38] Chris Woodruff. 10 Sereika datasets and 8 Srinivas datasets. (Accessed 24 May 2026). url: 10.6084/m9.figshare.31052200.v1.

[39] Chris Woodruff. Compressed files providing blastn databases used for the ASV paper analyses, together with gut microbiome definition files. (Accessed 24 May 2026). url: 10.6084/m9.figshare.31041886.v1.

