## Supplementary Material for "Strain resolution and wide relative abundance quantification of bacterial microbiomes with nanopore sequencing of the ribosomal RNA operon"

### Supplementary Material to Microbiome sub-species identification and abundance estimation with Nanopore-sequenced ribosomal DNA Amplicons

Christopher J. Woodruff and Terence P. Speed

#### 1 Datasets derived from the Sereika et al. Whole Genome Sequencing dataset for the Zymo D6322 even microbiome

Here we provide a description of the method of deriving 16S and 23S ribosomal RNA gene datasets from the whole genome sequencing D6322 dataset of Sereika et al. [1]. This includes the sub-sampling process briefly introduced in the primary document. We also present more detail on the identification and relative abundance results from both the full (mSerF) 16S and 23S datasets, and the sub-sampled datasets.

#### 1.1 Deriving 16S and 23S rRNA gene sequences from whole genome sequencing datasets

The D6322 mock microbiome R10.4 dataset from [1] was reported to have an estimated read accuracy (based on reported Q-score for reads) of 98.13%, an observed modal read accuracy of 99.14% (based on read-mapping to reference genomes), and an N50 of 5,992 bp, and consisted of approximately 22.5 million reads. These statistics lead to an expectation that extraction of 16S rRNA genes of length approximately 1500 bases, and 23S sequences of length approximately 2500 bases, would provide a sufficient number of ONT PromethION sequenced reads for amplicon-based analyses of the D6322 mock microbiome. Also, these reads would include a moderate proportion that had a mean read error rate of less than 0.01 errors per base which is the recommended quality of reads for use in the RAD denoiser code. ([2]).

Multiple primer sets exist for laboratory-based extraction of regions of the 16S rDNA. In particular there are primers for near the start and towards the end of that gene. These are the obvious primers to be used for in-silico extraction of these genes. However, when applied to reads that have the error rate of nanopore-sequenced reads these primers have reduced reliability.

The algorithm for extracting 16S and 23S rDNA sequences - described for the 16S case - is as follows:-

- Filter out all reads whose length is less than a threshold - e.g. 1100 bases for the 16S
- Using a set of primers for the 16S 5' location (see Appendix for primers used) align all primers to each retained read, returning an alignments score, the percent identity (PID) between the primer and the read over the alignment length, and the

Levenshtein distance (Lv) between the primer sequence and the aligned section of the read.

- Let $L_p$ = primer length, cost of match ($c_M$) = -1, cost of mismatch ($c_{MM}$) = 2, cost of gap creation ($c_G$) = 2, cost of gap extension ($c_E$)=2, $n_M$ = number of matches, $n_{MM}$ =number of mis-matches, $n_G$ =number of gaps, $n_E$ = total extension of gaps beyond lngth of 1. For every alignment compute the following 3 boolean quantities :-
  - $b_1$ = score $> L_p - 3 * 2$.
  - $b_2$ = PID $> 100(1 - 2/L_p)$.
  - $b_3 = \begin{cases} Lv < 3, \text{ if } L_p > 15 \\ Lv < 2, \text{ otherwise} \end{cases}$
- The value of 3*2 in the definition of $b_1$ reflects a term of the form $(n_{MM} + n_G + n_E)(c_M + c_{MM})$ that would allow various combinations of mis-matches, gaps and extensions in the alignment.
- Combine these three conditions into $b = b_3 \cap (b_1 \cup b_2)$.
- Repeat the previous steps for any primer for this boundary.
- If any of the primers returns TRUE for $b$ consider the boundary found and record its location.
- Repeat the previous steps for the 16S 3' boundary.
- If both 16S 5' and 16S 3' boundaries are found, and their separation falls within a pre-specified length range, consider the 16S sequence found.

Note that, for lower quality reads, a prior method was developed. This used primer sets for locations within the 16S or 23S rDNA sequences corresponding to a reference set of locations for the *Escherichia coli K-12 MG1655* strain to get a set of location pairs, the first element of a pair being the location for the read of interest, the second being the location in the reference E.coli sequence. If the read of interest was a 16S sequence then a linear fit to the E.coli 16S locations would be a good fit, and of gradient close to 1. Analogously for the 23S. The current method is computationally less demanding than such a method and adequate for the dataset from the Sereika et al. study.

Extraction of the 23S rDNA sequences followed the same procedure as was used for the 16S rDNA sequences. The primers used for the 23S extraction, apart from two, were derived by Zhang [3] using an entropy approach and an extensive set of bacterial genomes from the Refseq complete genomes. All primers are listed in section 7 of this document.

Given effective extraction of the 16S and 23 rRNA genes, it is a simple step to extract the complete 16S-ITS-23S, if it exists. Some bacteria do not have all 16S and 23S sequences in the conventional rrn operon configuration and hence before extracting a putative rrn operon the separation of the 16S 3' and the 23S 5' should be checked to see if it falls below an upper threshold for the length of the ITS. In fact, such a process was used to provide a dataset of rrn operons. However, fewer than 1000 were identified and this was considered too low to provide a useful analysis set for this D6322 mock microbiome.

#### 1.2 Sub-sampled datasets from the full Sereika et al. 16S and 23S rRNA gene datasets

The 16S and 23S from the full Sereika et al. D6322 mock microbiome data were presented in the main document. The quality of the identification, coupled with the mostly clear separation of species in the UMAP plots, allow us to reasonably assume that the reads associated with ASVs assigned to a particular species are themselves truly of that species. On this assumption, we can proceed with sub-sampling. Table 1 specifies the 4 different sub-samplings considered. These allow illustration of the sensitivity of the denoising, noting that the D6322 mock microbiome only has 7 bacterial strains all of different genera. First, the UMAP visualisations of 16S and 23S rRNA gene data for each sub-sampled dataset are presented, followed by the species and gene scatterplots for each rRNA gene for each sub-sampling.

**Table 1** Sub-sampling factors on reads from species in the Sereika-derived 16S and 23S rRNA gene datasets used to generate four mock microbiomes for each such gene that cover a range of either 50:1 or 100:1 in relative cellular abundance of the species. Species key: Bs=*Bacillus subtilis* , Ef=*Enterococcus faecalis* , Ec=*Escherichia coli* , Lm=*Listeria monocytogenes* , Pa= *Pseudomonas aeruginosa* , Se=*Salmonella enterica* , Sa=*Staphylococcus aureus* .

| Dataset | Bs | Ef | Ec | Lm | Pa | Se | Sa |
|---|---|---|---|---|---|---|---|
| Sub1 | 0.1 | 1 | 1 | 1 | 1 | 0.5 | 0.2 |
| Sub2 | 0.5 | 0.01 | 0.1 | 1 | 1 | 1 | 1 |
| Sub3 | 1 | 0.02 | 1 | 1 | 1 | 1 | 0.02 |
| Sub4 | 1 | 1 | 0.1 | 1 | 1 | 1 | 0.02 |

##### 1.2.1 UMAP Visualisations of Sub-sampled datasets

For each sub-sampling a set of 4 2D UMAP plots is presented. On the left, labelling of ASVs is at the species level, while on the right it is at the strain level. The upper row of the plots is for the 16S dataset and the lower is for the 23S dataset. For large datasets - e.g. 100,000 reads - the computation of pairwise distances between reads, which is the standard approach with UMAP calculations, is very demanding. However an alternative approach [4] involves subsampling the reads, computing a 2D UMAP model from the subsample and then embedding the remaining reads into the space characterised by that model. This has been implemented in R in the uwot package, and is used here for the large datasets from simulated reads, and also in this sub-section of visualisations, to simply demonstrate its performance compared to the full computation. In this sub-section of Supplementary Material all species-level UMAP plots (the left-hand columns) are computed with the full set of distance calculations, while the strain-level labeled plots are computed by the embedding method. Note that the labeling itself is unaffected by the method of UMAP calculation.

Figures 1 to 4 follow the same pattern, covering sub-samplings 1 to 4 of Table 1. Consider Figure 1. The pattern of read clusters and ASV locations are visually identical for the two 16S plots and for the two 23S plots. The 16S species plot shows 8 species, the *Shigella flexneri* being a mis-identification. The other species comprise the full bacterial mock microbiome. Apart from the *Shigella flexneri* being located close to a cluster of *Escherichia coli* ASVs, all other ASV clusters are single species clusters. The 23S plot has only species that are in the mock microbiome, and all ASV clusters are single species clusters.

The strain-labelled plots on the right-hand side have all ASVs marked by a black cross, and, additionally, each ASV has centred on it a larger coloured symbol as specified in the legend. Careful inspection of the cluster at approximately (-5, -5) shows 3 different symbols, each of which corresponds to a different strain of Listeria monocytogenes. Similarly, a cluster at approximately (-2.5,0) indicates 3 different Staphylococcus strains. There is only a single strain of both of these species in the mock microbiome. The mis-identification almost certainly arises from tied alignment qualities coupled with the selection of the label for an ASV being that strain occurring first in the blastn output - which is alphabetically ordered within ties. The variation would be due to small differences in ASVs or in the underlying operons of the strain favouring one strain over the other.

The number of ASVs for each species is moderately correlated with the number of reads of the different species. For Figure 1 *Bacillus subtilis* has a sampling weight of 0.1 and it is seen that there is only one ASV for this species in both the 16S and 23S plots. A similar result will be seen in other figures where the weighting range is greater than the 10:1 range of Figure 1.

Figures 2, 3 and 4 provide corresponding plots for the other subsampled datasets.

Analysis of these sub-sampled datasets shows very good identification being achieved, even at relative abundances of about 0.01. The read count associated with some of the ASVs formed is less than 10 - e.g. *Escherichia coli* in Sub3 16S dataset analysis.

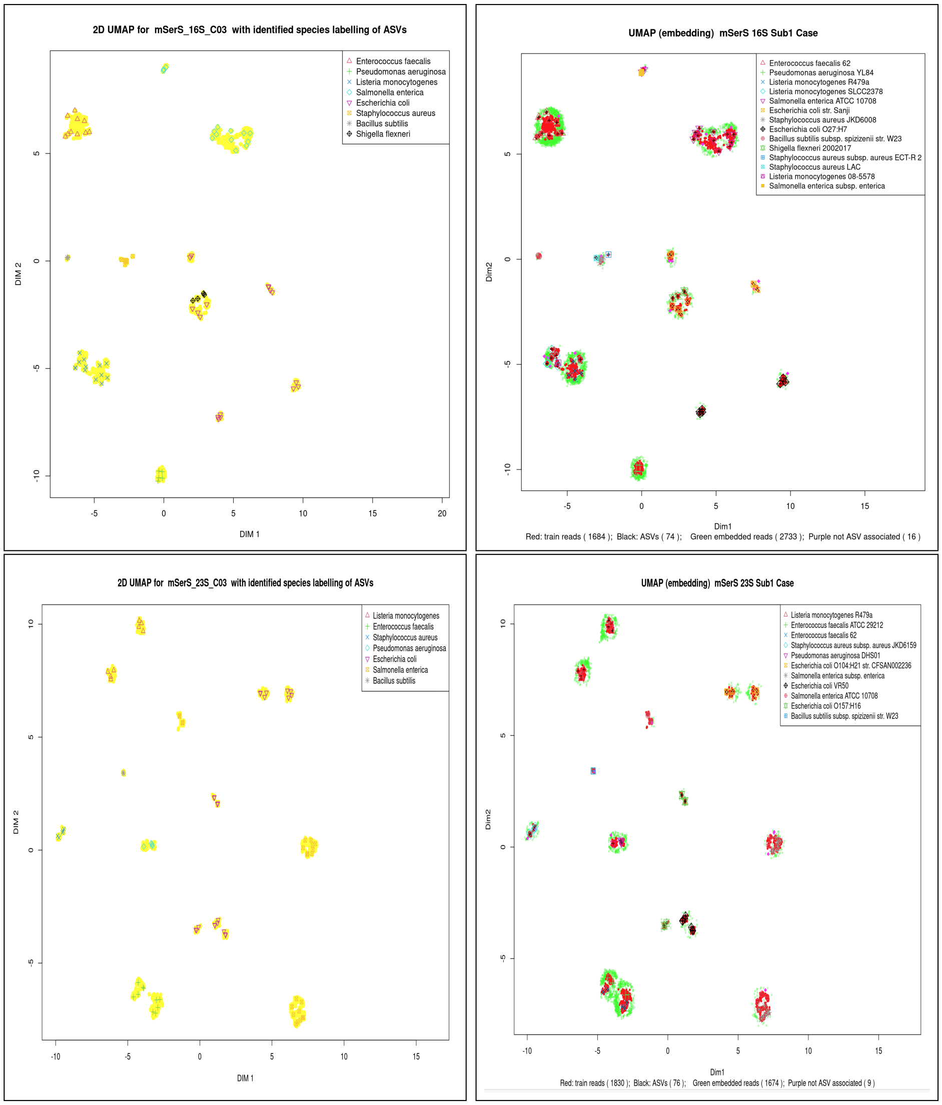


**Fig. 1** UMAP-derived 2D representation of 16S (upper) and 23S (lower) rRNA gene ASVs (black symbols) and reads (yellow dots) for the subsampled dataset, mSerS Sub1, with species labelling (left) and strain (right) labelling to which these ASVs are classified.Weights of less than 1 in the subsampling are 0.1 for *Bacillus subtilis,* 0.2 for *Staphylococcus aureus* and 0.5 for *Salmonella enterica.*

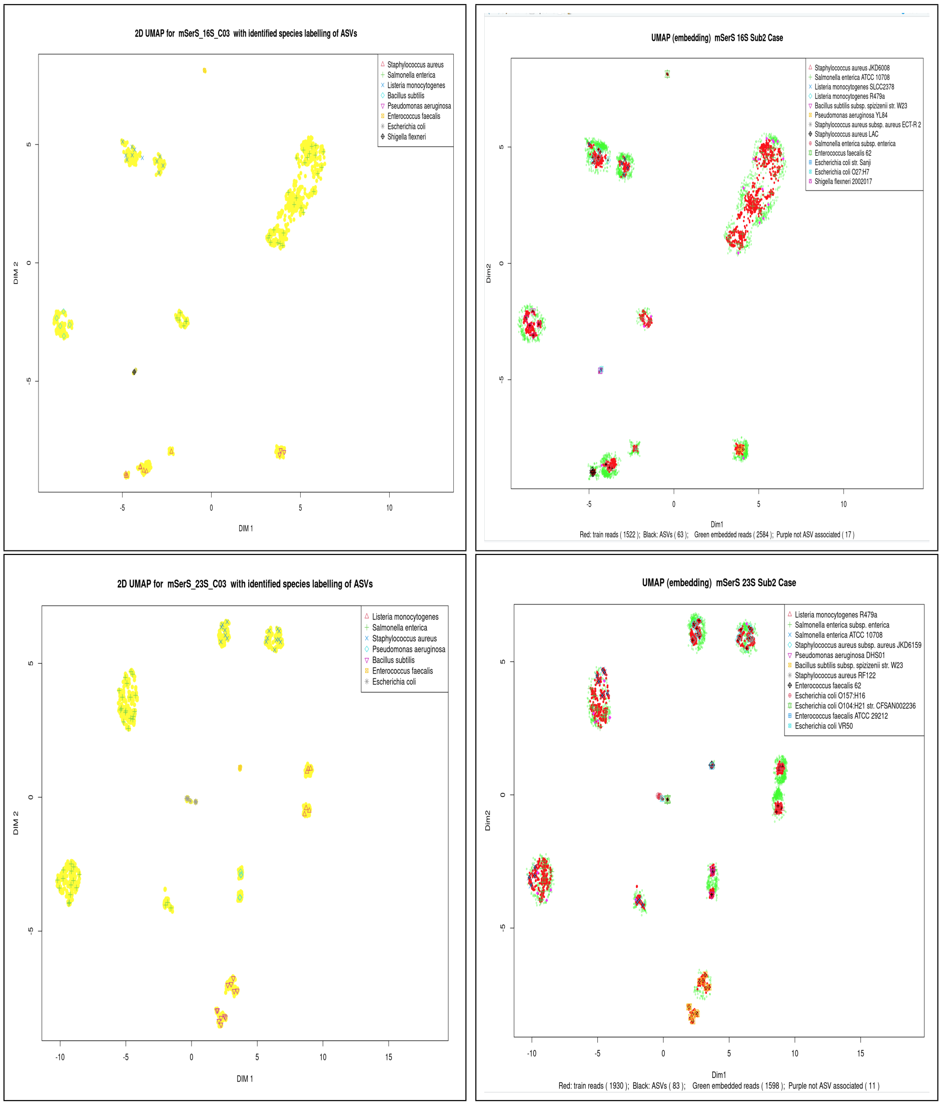


**Fig. 2** UMAP-derived 2D representation of 16S (upper) and 23S (lower) rRNA gene ASVs (black symbols) and reads (yellow dots) for the subsampled dataset, mSerS Sub2. Weights of less than 1 in the sub-sampling are 0.01 for *Enterococcus faecalis*, 0.1 for *Escherichia coli* and 0.5 for *Bacillus subtilis*.

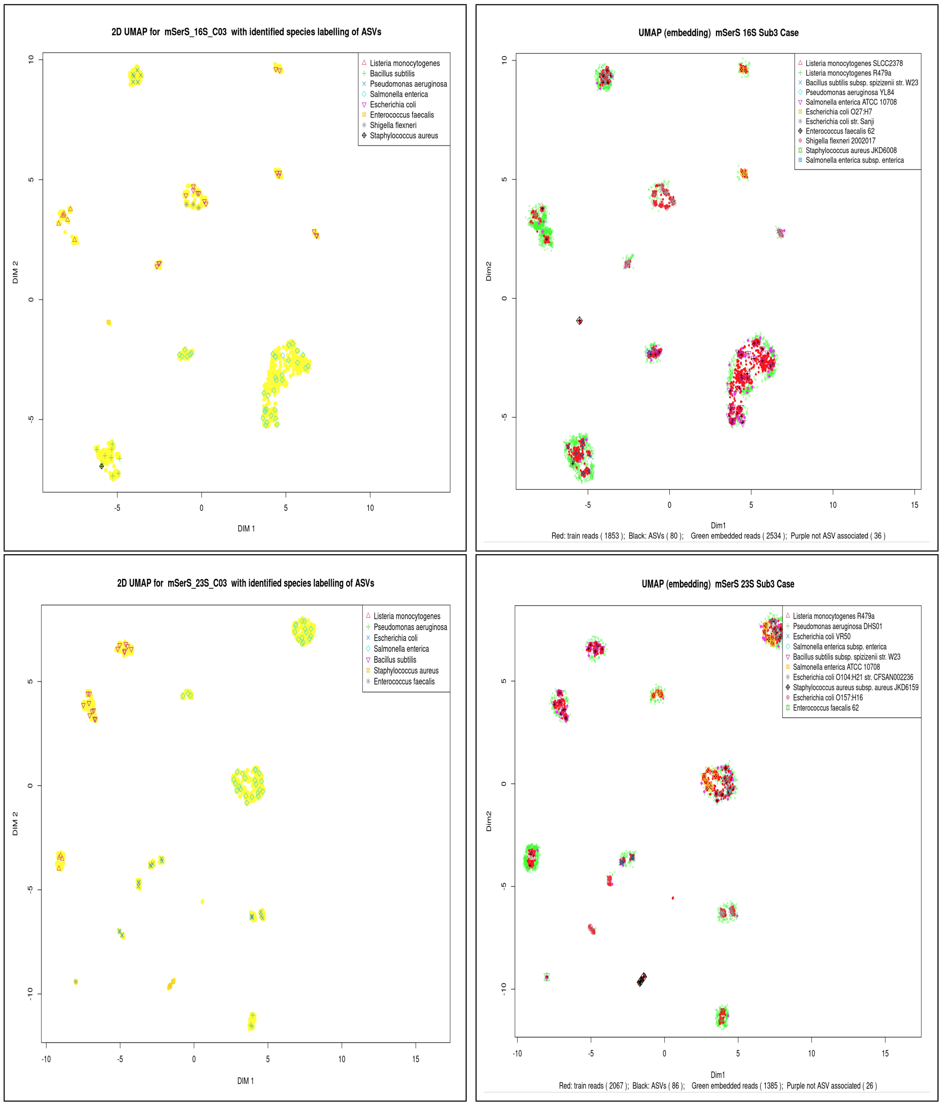


**Fig. 3** UMAP-derived 2D representation of 16S (upper) and 23S (lower) rRNA gene ASVs (black symbols) and reads (yellow dots) for the subsampled dataset, mSerS Sub3. Weights of less than 1 in the sub-sampling are 0.02 for both *Enterococcus faecalis* and *Staphylococcus aureus*.

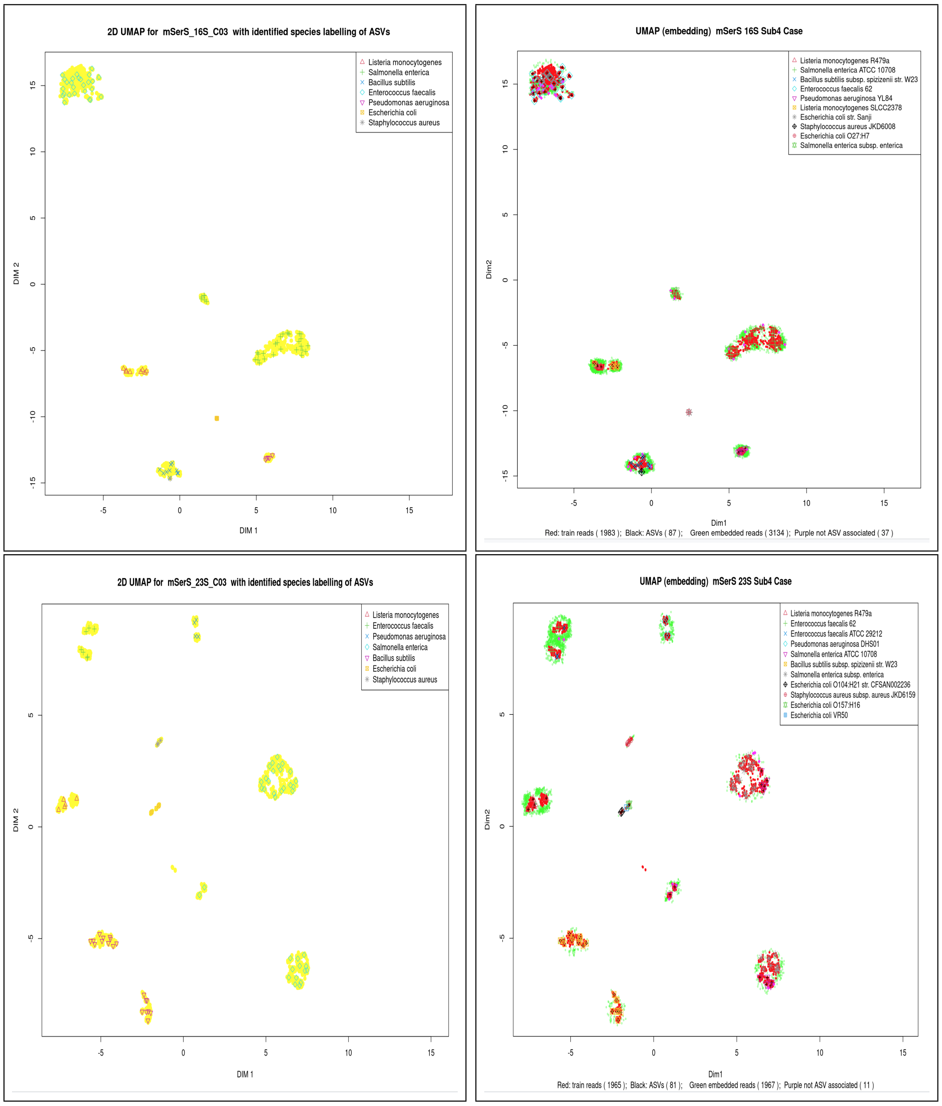


**Fig. 4** UMAP-derived 2D representation of 16S (upper) and 23S (lower) rRNA gene ASVs (black symbols) and reads (yellow dots) for the subsampled dataset, mSerS Sub4. Weights of less than 1 in the sub-sampling are 0.02 for *Staphylococcus aureus*, and 0.1 for *Escherichia coli*.

##### 1.2.2 Observed vs. designed relative abundances from the sub-sampled datasets

Figure 5 and Figure 6 present cellular relative abundances at the species level for the 4 sub-samplings from 16S rRNA gene and 23SrRNA gene analysis, respectively. Note that we are deriving the expected relative abundances based on an assumption that the analyses of the full 16S and 23S datasets are exact, whereas they will almost certainly have some mis-assignment of reads to ASVs. Additionally, a small number of ASVs are probably incorrectly identified. Hence there will probably be some error in the expected abundances, which we ignore .

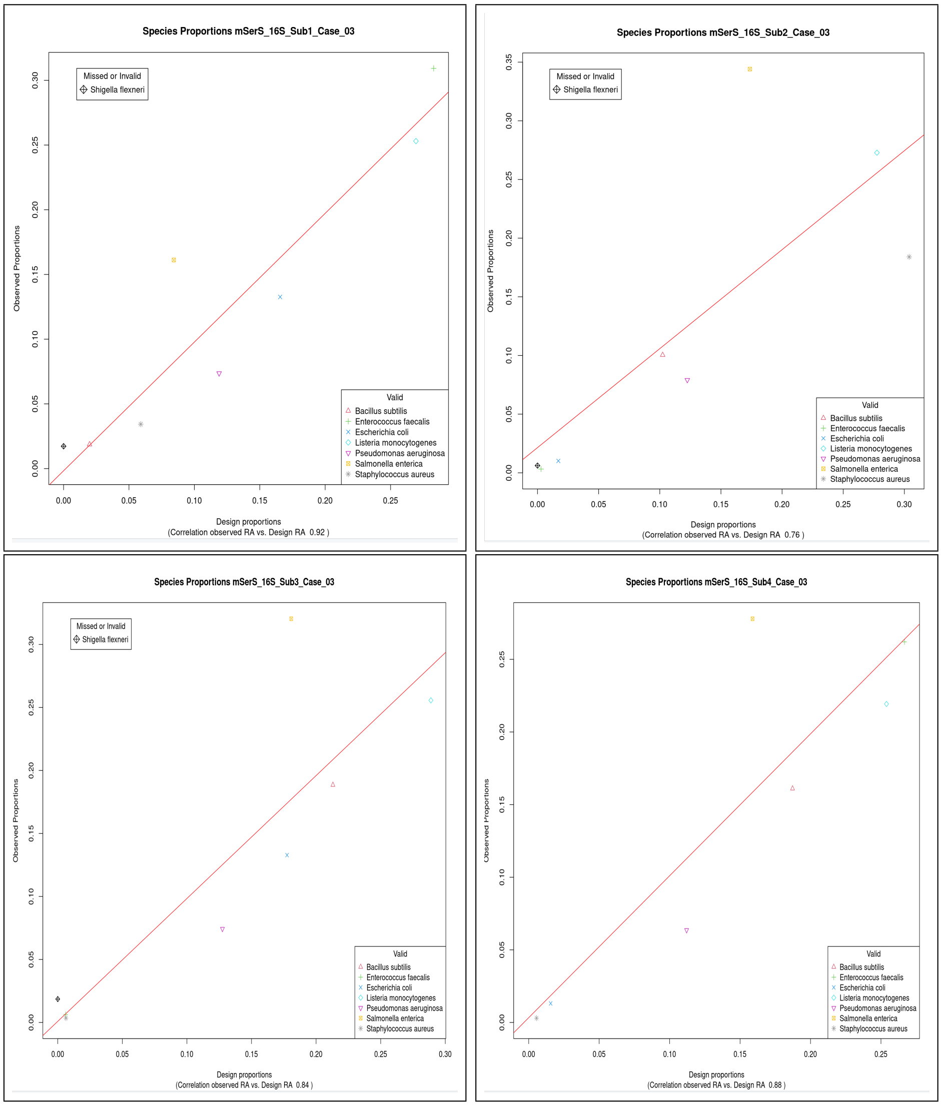


**Fig. 5** Relation between the observed relative abundances of species and those of the designed microbiomes for sub-samples Sub1 to Sub4 as determined from 16S rRNA genes. The red line is a linear fit to the data - excluding the mis-identification *Shigella flexnerii*.

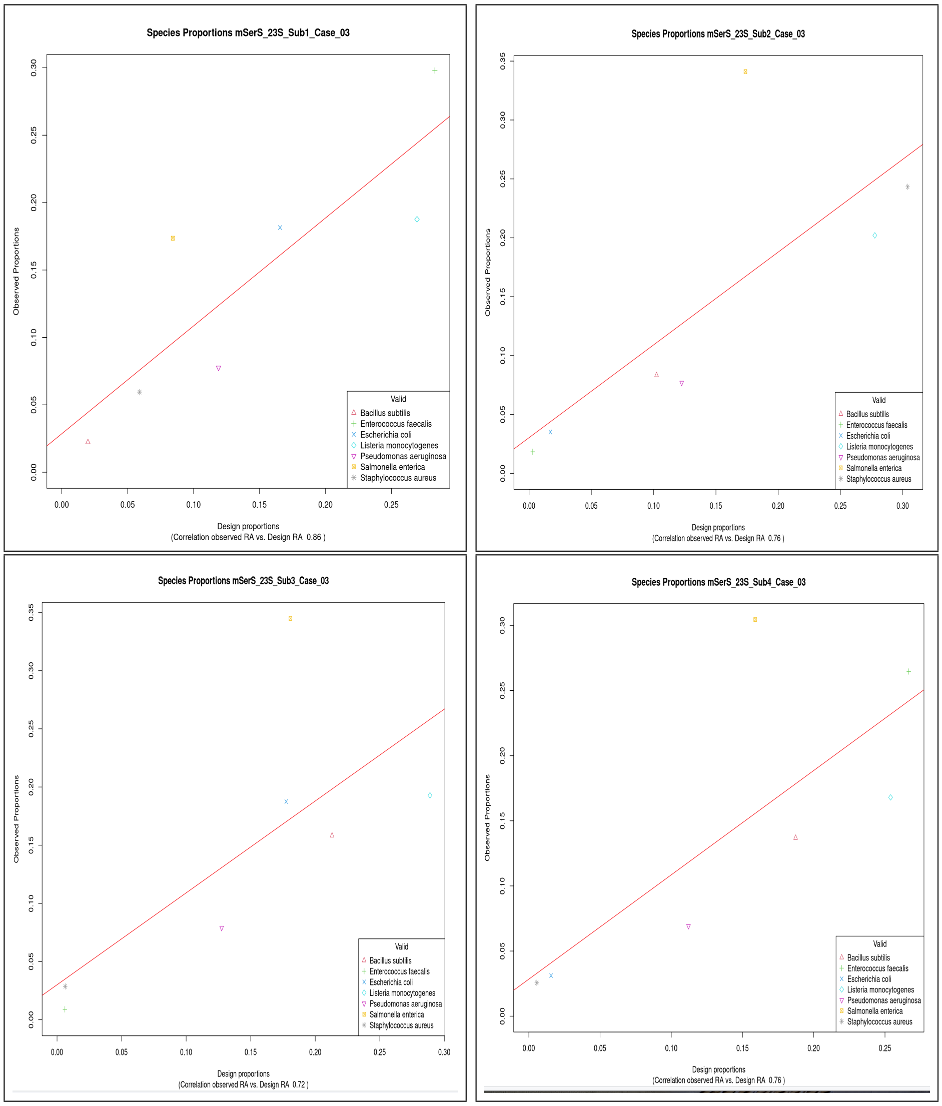


**Fig. 6** Relation between the observed relative abundances of species and those of the designed microbiomes for sub-samples Sub1 to Sub4 as determined from 23S rRNA genes.

### 2 Datasets from the Srinivas et al. 16S-ITS-23S ATCC and Zymo D6322 mock microbiomes

This section presents data from primer pairs 3 (519F, 2241R) and 4 (519F, 2428R), but only where it is not trivially different to results in the main document. So no data on the error rate distribution is presented, for instance. However in figure 7 the variation of error rate with read length is shown for primer pairs 3 and 4.

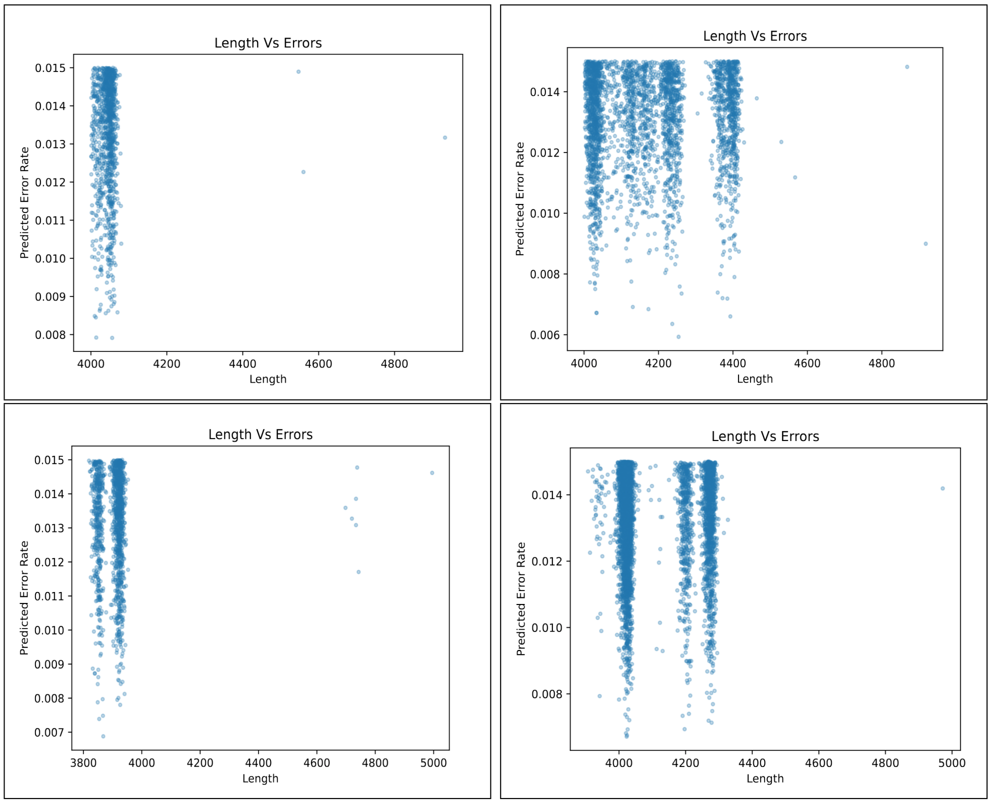


**Fig. 7** Error rate variation with read length for reads processed by the RAD denoiser from the 16S-ITS-23S rRNA operons of the ATCC (upper) and Zymo D6311 mock microbiomes using primer pairs 3 (left) and 4 (right).

Visualisation of species-level-labelled ASV data using UMAP is shown in Figure 8 for primer pairs 3 and 4 for the ATCC datasets, and, in Figure 9, for the Zymo datasets.

Scatterplots of observed relative abundance vs. expected relative abundance are provided for all primer pairs with the ATCC even microbiome here. Figure 10 gives species data, Figure 11 the corresponding genus data. There is no value in other scatterplots that are not presented in the main document. There is clearly no basis for confidence in quantification here.

The various scatterplot of Figures 10 and 11 are all results of relative abundance measurement at either species or genus level on the same microbiome - the difference between SA1 through to SA4 being in what primer set defined the reads processed. The considerable variation between plots indicates the unsuitability of these datasets for processing by the method presented in this paper. These methods require data of higher quality, as exemplified in the Sereika et al. data and in the simulated reads datasets.

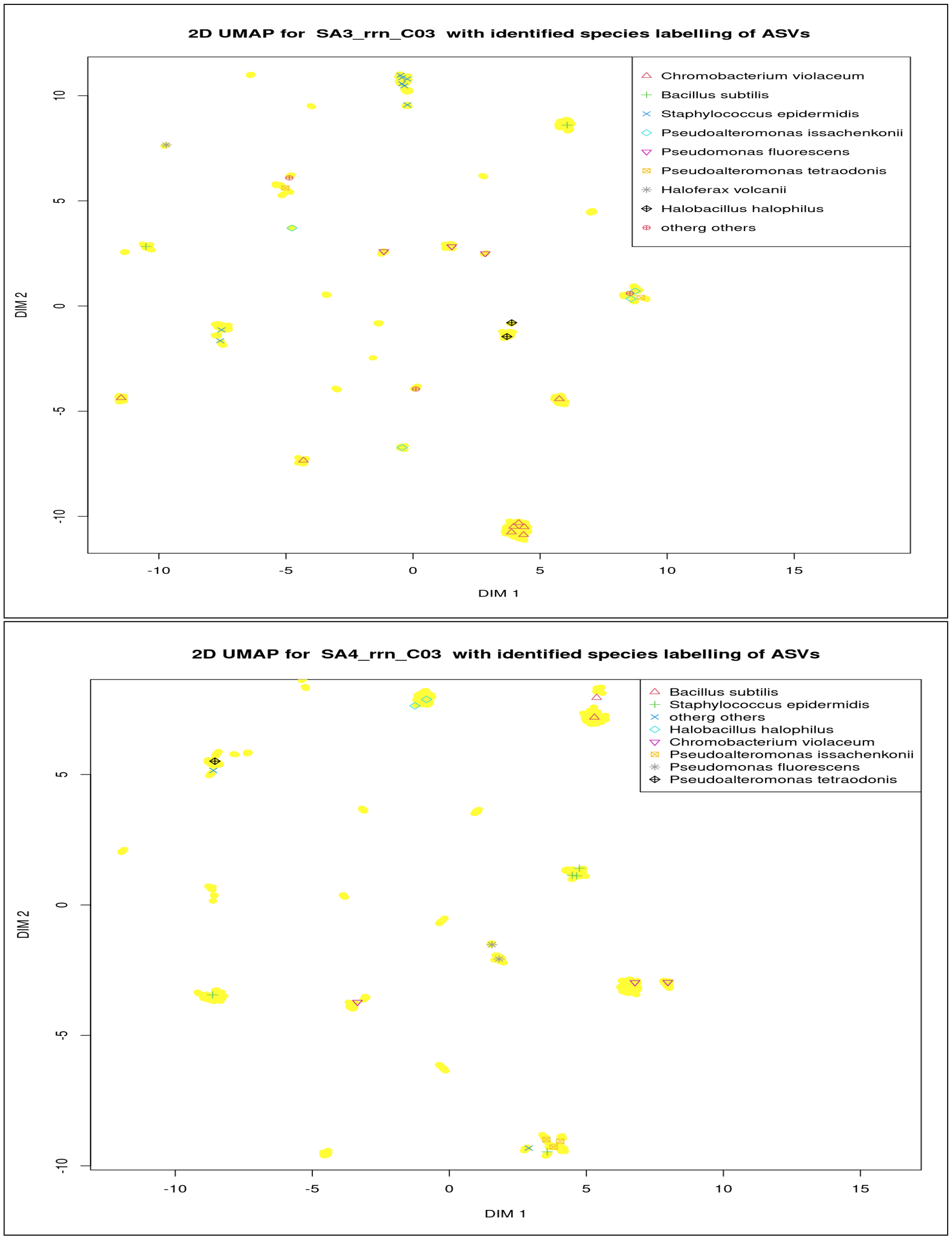


**Fig. 8** 2D UMAP representation of the relationship between ASVs and their associated read clusters for the 16S-ITS-23S sequences extracted using primer pair 3 and 4 from the Srinivas et al. study for the ATCC even microbiome. Note that 6 of the 8 listed identified species of the upper plot are valid and 2 are missing, while 5 of 7 are valid and 3 are missing in the lower plot.

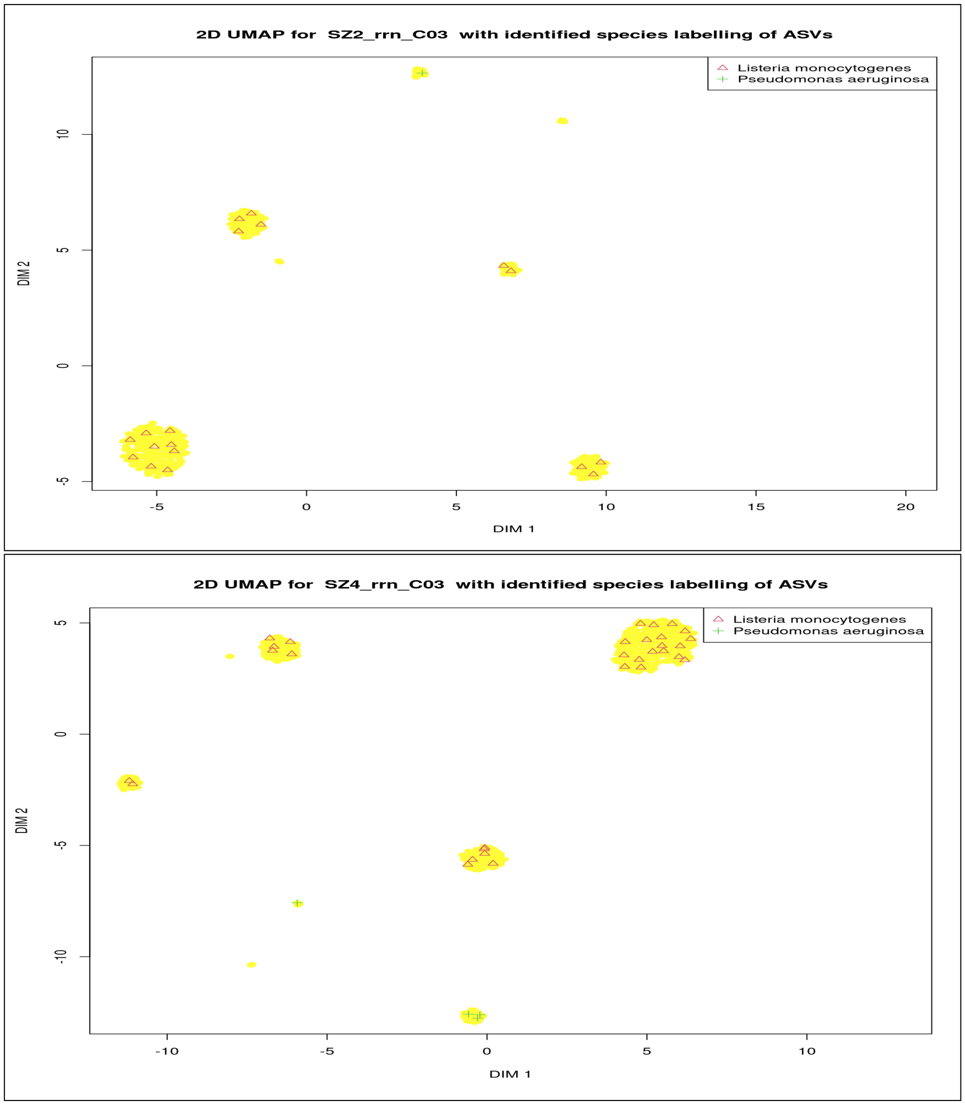


**Fig. 9** 2D UMAP representation of the relationship between ASVs and their associated read clusters for the 16S-ITS-23S sequences extracted using primer pairs 2 (upper) and 4 (lower) from the Srinivas et al. study of the Zymo D6311 logarithmic microbiome.Two species are correctly identified in both these plots, 5 are missing.

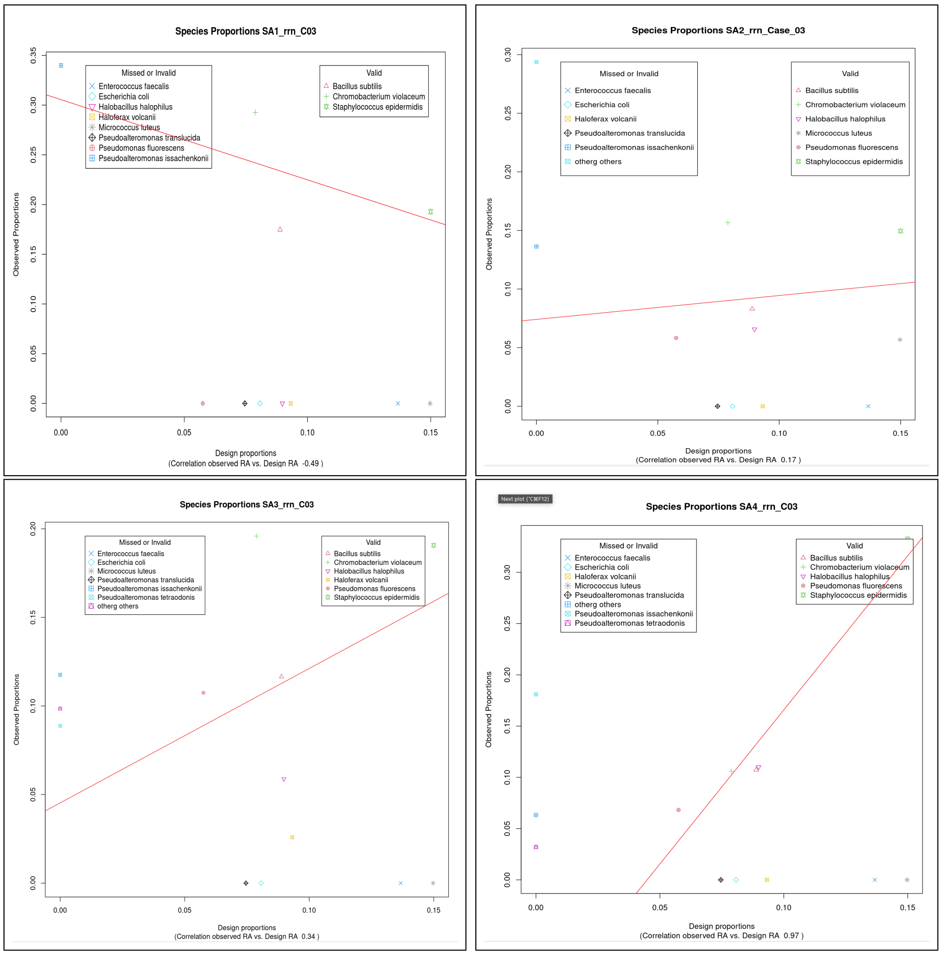


**Fig. 10** Relation between the observed species relative abundance and that of the designed microbiome as determined from the 16S-ITS-23S sequences extracted using primer pairs 1 to 4 for the ATCC mock microbiome. Missed species (or genera) appear with zero observed proportions and are identified in decreasing proportion value in the legend. Invalid species appear at zero design proportions, and follow the missed species (or genera) in the legend.

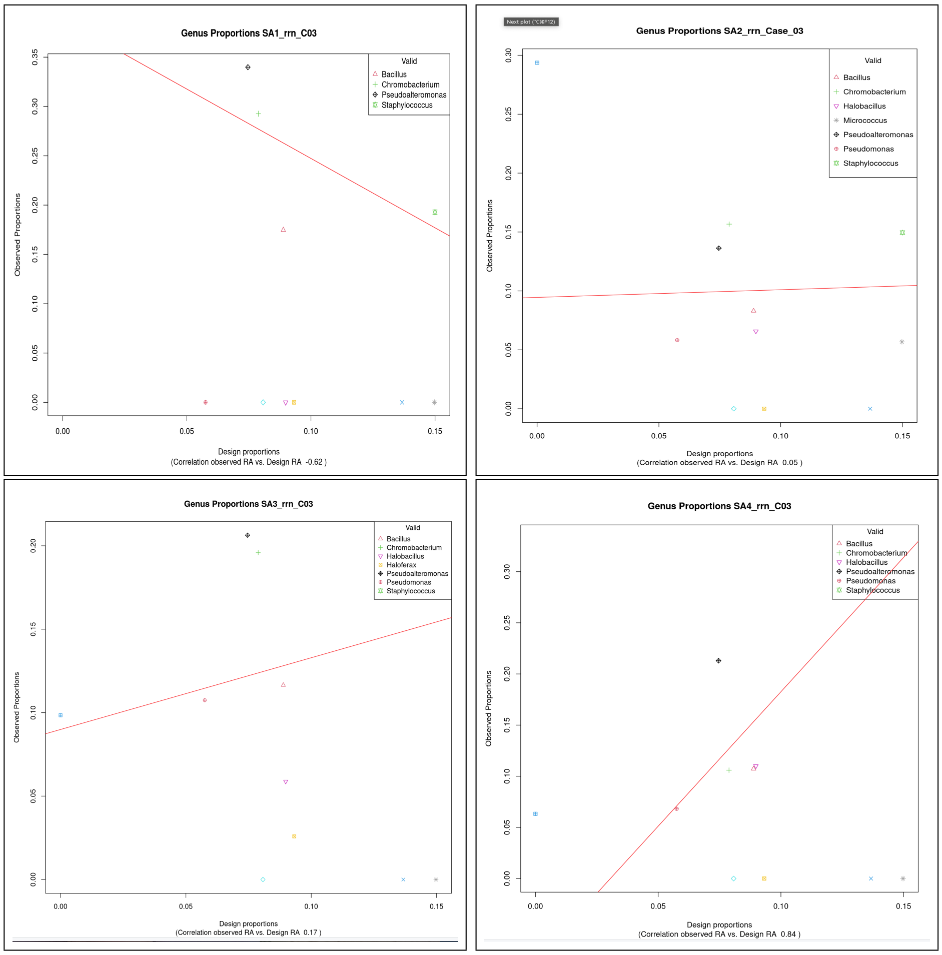


**Fig. 11** Relation between the observed genus relative abundance and that of the designed microbiome as determined from the 16S-ITS-23S sequences extracted using primer pairs 1 to 4 for the ATCC mock microbiome. Missed species (or genera) appear with zero observed proportions and are identified in decreasing proportion value in the legend. Invalid species appear at zero design proportions, and follow the missed species (or genera) in the legend.

### 3 Simulating ONT reads

To create the mKB mock microbiomes based on the Gut Feeling Knowledge Base of [5] 291 *rrn* operons from 59 bacterial strains were identified, and simulated reads generated from each of these using Wick's badread code [6]. That code and its use is well documented on Github [7]. Here we briefly document the specific manner of use in this work. For each operon a number of simulated reads greater than actually required for the mock microbiome are generated and stored. For each operon the simulation is run independently of other operons. This study was not concerned with the process of the preparation of DNA for sequencing, hence the various parameters dealing with sample preparation were not considered - no adapters nor chimeras. Also we chose not to consider any junk or random sequences. Each operon has its particular length in the reference database. This length was used as the mean fragment length with a standard deviation of 5. Read quality is controlled through the identity quantity, which is parameterised either via the distribution of percent identity or by the distribution of the phred-related Qscore. Wick recommends the latter for higher quality simulations, and this is what was used here. The mean Qscore was set to 30 with a standard deviation of 4. This resulted in reads whose mean error rate distribution in the range below 0.01 errors per base fell between the distributions found for the two sources of real data.

### 4 Observed operons, ASVs, and operon- and strain-level resolution analysis for species having multiple strains

Four species having multiple strains in the mock microbiome are considered in the following. For each species the analysis of blastn alignment of ASVs gives a set of operons, with it commonly being the case that multiple ASVs have the same best-aligned operon from the reference DB. In this section, for each species we present a table giving some detail on the alignment of ASVs and operons, a graphical representation of the distance matrix between all the operons and all the associated ASVs, and a UMAP2D plot illustrating the relation between the operons and the ASVs.

#### 4.1 *Bifidobacterium longum*

There are 12 unique operons observed, with 4, 2, 2, 2, 1, 1, 2, 2, 2, 2, 1, and 1 associated ASVs, respectively. The identities of the observed operons are given in Table 2, together with the length of the alignment, Alen, and the number of mismatches (MM), gaps, and the percent identity of bases of the aligned sequences. It should be noted that some less specific entries in the database - e.g. entries such as *Bifidobacterium longum subsp. longum* that have no further detail in the strain except for the operon number may have a large number of operons - e.g we see *Bifidobacterium longum subsp. infantis* with operon 87 appearing. This tends not to be the case for entries with a proper strain name such as *Bifidobacterium longum subsp. longum GT15*.

**Table 2** Observed operons for *Bifidobacterium longum* , together with alignment details for the first-listed ASV for each operon

| Operon | Alen | MM | Gap | PID |
|---|---|---|---|---|
| subsp. longum GT15 op5 | 5023 | 0 | 0 | 100 |
| subsp. longum op13 | 5023 | 0 | 0 | 100 |
| DJO10A op1 | 5022 | 0 | 0 | 100 |
| DJO10A op2 | 5024 | 0 | 0 | 100 |
| DJO10A op3 | 5023 | 0 | 0 | 100 |
| DJO10A op4 | 5023 | 0 | 0 | 100 |
| subsp. longum JDM301 op3 | 5021 | 0 | 0 | 100 |
| subsp. longum JDM301 op2 | 5021 | 0 | 0 | 100 |
| subsp. longum JDM301 op1 | 5020 | 0 | 0 | 100 |
| subsp. infantis op87 | 5023 | 0 | 0 | 100 |
| subsp. infantis 157F op4 | 5023 | 0 | 0 | 100 |
| subsp. infantis 157F op2 | 5023 | 0 | 0 | 100 |

This species gives alignments that are all 5020 bases or slightly more long, and these are perfect alignments. This illustrates the effectiveness of the denoising algorithm used, RAD.

Figure 12 provides a visualisation of the distance matrix between the 12 reference operons that are identified as being best aligned to the ASVs and those ASVs. In viewing this figure consider the 12 x 12 sub-matrix in the lower left corner is the distance matrix for the operons alone. It can be seen that the diagonal elements have the smallest, or equal smallest values of distance. Above row 12 we have the distances between operons and ASVs represented. Careful viewing allows one to see that - apart from operon 1 - there are good matches to exactly those ASVs that are associated with those operons. This is exactly what is expected given the excellent alignments seen in Table 2. Operon 1 has 2 perfect alignments and 2 poorer alignments for the 4 ASVs associated with it. The poorer alignments for operon1 to its associated ASVs are due to the ASVs being markedly shorter than the operon.

Figure 13 provide a visualisation of the relationship between the observed operons - labelled with uppercase letters - and their associated ASVs - labelled with lower case

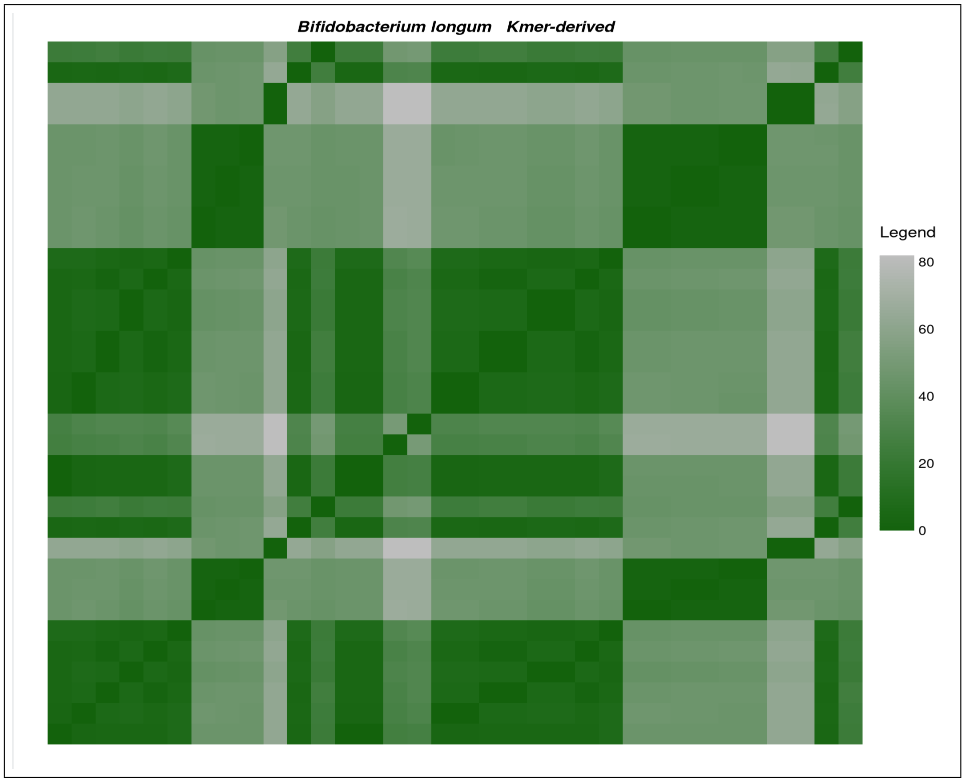


**Fig. 12** A distance matrix between reference operons of those operons observed and the ASVs for which these operons give optimal alignment for *Bifidobacterium longum* in the mock microbiome. The distance measure is the approximate kmer measure.

letters. We have used a single colour for an operon and its associated ASVs. From this figure we see that, for *Bifidobacterium longum*, an operon and its associated ASVs are all quite closely located. We see that operons G,H,I,J are quite close to each other.

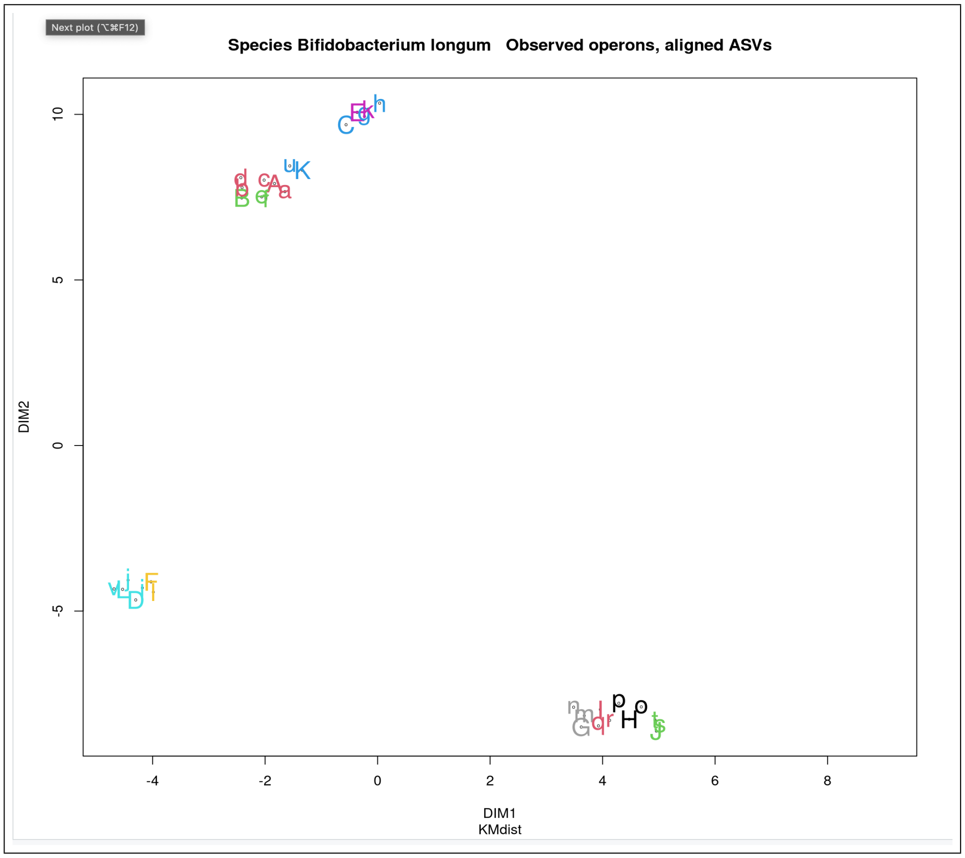


**Fig. 13** A UMAP2D projection plot for a distance matrix between those reference operons observed and the ASVs for which these operons give optimal alignment for *Bifidobacterium longum* in the mock microbiome. The distance measure is the approximate kmer measure. Operon A here is operon 1, and the 4 associated ASVs, a, b, c, d, are labelled the same colour. Note that colouring cycles after 9 operons.

#### 4.2 *Bacteroides fragilis*

For *Bacteroides fragilis*there are 11 operons and a total of 40 ASV associated with them. Table 3 gives the identities of the observed operons and some details on their ASV alignments. Note that the multiple ASVs aligned to a particular operon can be giving rather different alignment lengths. Our table only gives the length of the lowest index ASV (which tends to have the largest number of associated reads).

**Table 3** Observed operons for *Bacteroides fragilis* , together with alignment details for the first-listed ASV for each operon

| Operon | Alen | MM | Gap | PID |
|---|---|---|---|---|
| YCH46 Op5 | 4894 | 0 | 0 | 100 |
| YCH46 Op3 | 4971 | 0 | 0 | 100 |
| YCH46 Op4 | 4896 | 0 | 0 | 100 |
| YCH46 Op2 | 4902 | 0 | 0 | 100 |
| YCH46 Op1 | 4974 | 0 | 0 | 100 |
| YCH46 Op6 | 4894 | 0 | 0 | 100 |
| 638R Ops4,5 | 4905 | 0 | 0 | 100 |
| 638R Op1 | 4896 | 0 | 0 | 100 |
| 638R Op2 | 4897 | 0 | 0 | 100 |
| 638R Op3 | 4897 | 0 | 0 | 100 |
| 638R Op6 | 4904 | 0 | 0 | 100 |

We see that there are 2 strains, each having 6 operons identified, with only operons 4 and 5 of strain 638R being duplicates. There is clear strain discrimination.

Figure 14 gives a visualisation of the distance matrix for the observed operons and the set of ASVs for this species. We note that there is generally less variation for the inter-operon distances (11 x 11 matrix at the lower left of the figure) than for the inter-ASV distances. We also see the pair of approximate curved line of (operon, matched ASV) pairs appearing as darker blocks symmetrically placed about the diagonal - visible with greater difficulty in the corresponding plot for *Bifidobacterium longum*.

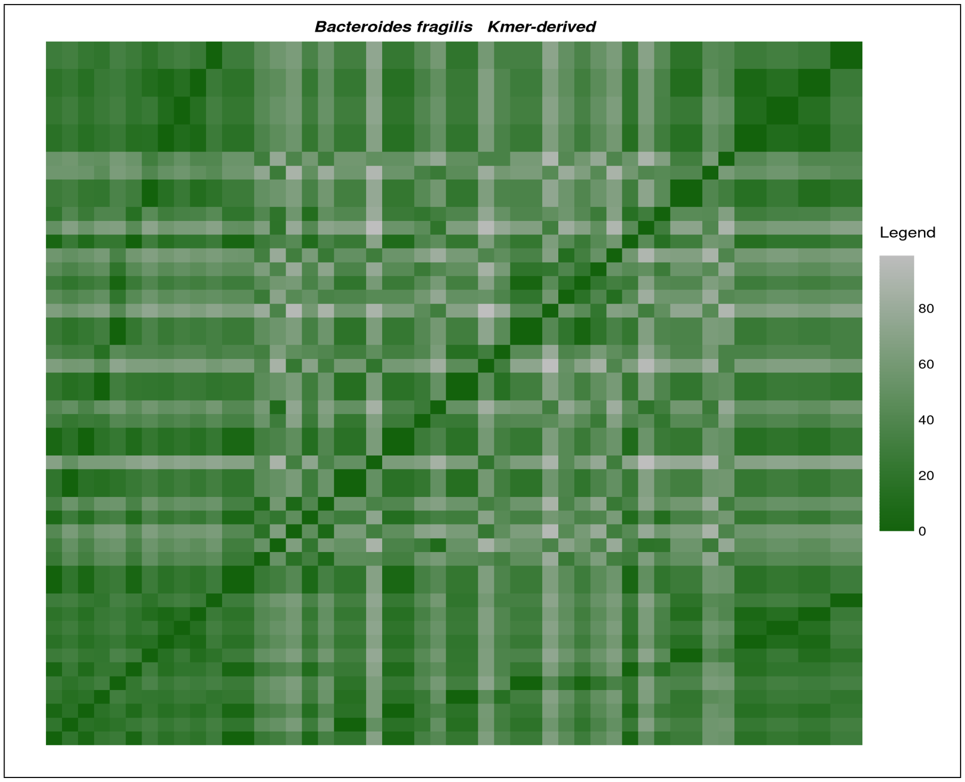


**Fig. 14** A distance matrix between reference operons of those operons observed and the ASVs for which these operons give optimal alignment for *Bacteroides fragilis* in the mock microbiome. The distance measure is the approximate kmer measure.

Figure 15 is a UMAP2D visualisation of the relationship between the operons and the associated ASVs. The 638R operons and their associated ASVs are all located in the lower left corner. The YCH46 data are relatively highly dispersed and we see ASVs , such as c, e, g, y that are quite remote from their operons. These are ASVs that are markedly shorter than the operons - e.g. lengths of around 4400 bases rather than 4900 bases.

This data provides clear resolution of the two strains present. This is achieved despite the significant dispersion of the YCH46 operons and their associated ASVs. It is the fact that the analysis is done from the operon level that allows this resolution.

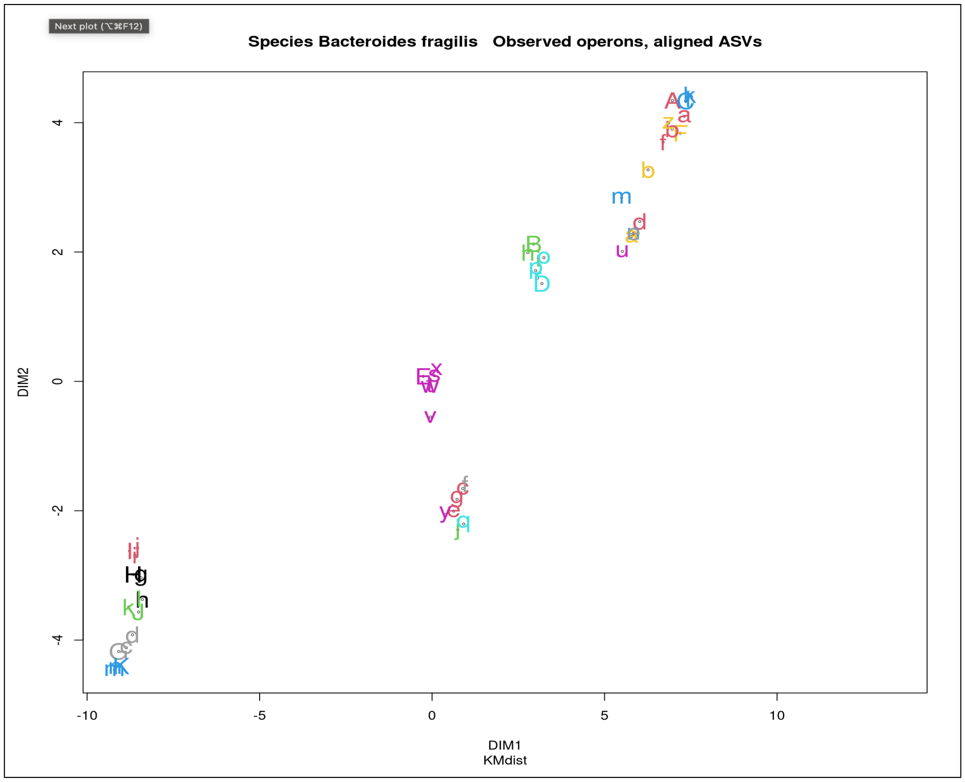


**Fig. 15** A UMAP2D projection plot for a distance matrix between those reference operons observed and the ASVs for which these operons give optimal alignment for *Bacteroides fragilis* in the mock microbiome. The distance measure is the approximate kmer measure. Operons are labelled with uppercase letters, ASVs with lower case letters. Colour differentiates one set of an operon and its associated ASVs from other sets.

#### 4.3 *Escherichia coli*

For *Escherichia coli* there are 35 operons and 61 associated ASVs. The greater complexity of this species due to the number of strains and, for some observed strains, their degree of similarity, means that the table below has limitations. Table 4 is complementary to this one in that it gives the other strains and operons in the tied optimal best alignments.

From Table 4 there are 10 observed strains. However the more thorough analysis of the operon data available from Table 4 reduces the number of strains by dealing with the ambiguity from tied optimal alignments.

**Table 4** Observed operons for *Escherichia coli* , together with alignment details for the first-listed ASV for each operon

| Operon | Alen | MM | Gap | PID |
|---|---|---|---|---|
| CFT073 Op4 | 4800 | 0 | 0 | 100 |
| CFT073 Op7 | 4892 | 0 | 0 | 100 |
| CFT073 Op1 | 4892 | 0 | 0 | 100 |
| CFT073 Op5 | 4800 | 0 | 0 | 100 |
| CFT073 Op2 | 4800 | 0 | 0 | 100 |
| CFT073 Op6 | 4800 | 0 | 0 | 100 |
| CFT073 Op3 | 4892 | 0 | 0 | 100 |
| UMI46 Op6 | 4800 | 0 | 0 | 100 |
| SMS-3-35 Ops2,6 | 4801 | 0 | 0 | 100 |
| SMS-3-35 Op5 | 4801 | 0 | 0 | 100 |
| SMS-3-35 Op3 | 4890 | 0 | 0 | 100 |
| SMS-3-35 Op1 | 4883 | 0 | 0 | 100 |
| SMS-3-35 Op3 | 4800 | 0 | 0 | 100 |
| SMS-3-35 Op4 | 4790 | 0 | 0 | 100 |
| BL21...AG Op7 | 4793 | 0 | 0 | 100 |
| BL21...AG Op3 | 4887 | 0 | 0 | 100 |
| BL21...AG Op6 | 4892 | 0 | 0 | 100 |
| BL21...AG Op1 | 4886 | 0 | 0 | 100 |
| 1303 Op4 | 4799 | 0 | 0 | 100 |
| 1303 Op2 | 4877 | 0 | 0 | 100 |
| 1303 Op5 | 4892 | 0 | 0 | 100 |
| 1303 Op6 | 4885 | 0 | 0 | 100 |
| 1303 Op3 | 4892 | 0 | 0 | 100 |
| 1303 Op1 | 4883 | 0 | 0 | 100 |
| UMNK88 Op3 | 4894 | 0 | 0 | 100 |
| UMNK88 Op5 | 4894 | 0 | 0 | 100 |
| UMNK88 Op6 | 4894 | 0 | 0 | 100 |
| UMNK88 Op7 | 4894 | 0 | 0 | 100 |
| UMNK88 Op1 | 4894 | 0 | 0 | 100 |
| UMNK88 Op4 | 4800 | 0 | 0 | 100 |
| Nissle 1917 Ops1,2,5,6 | 4583 | 0 | 0 | 100 |
| Nissle 1917 Op6 | 4488 | 0 | 0 | 100 |
| REL606 Op6 | 4804 | 0 | 0 | 100 |
| NU14 Op3 | 4892 | 0 | 0 | 100 |
| O18:H1 Op5 | 4486 | 0 | 0 | 100 |

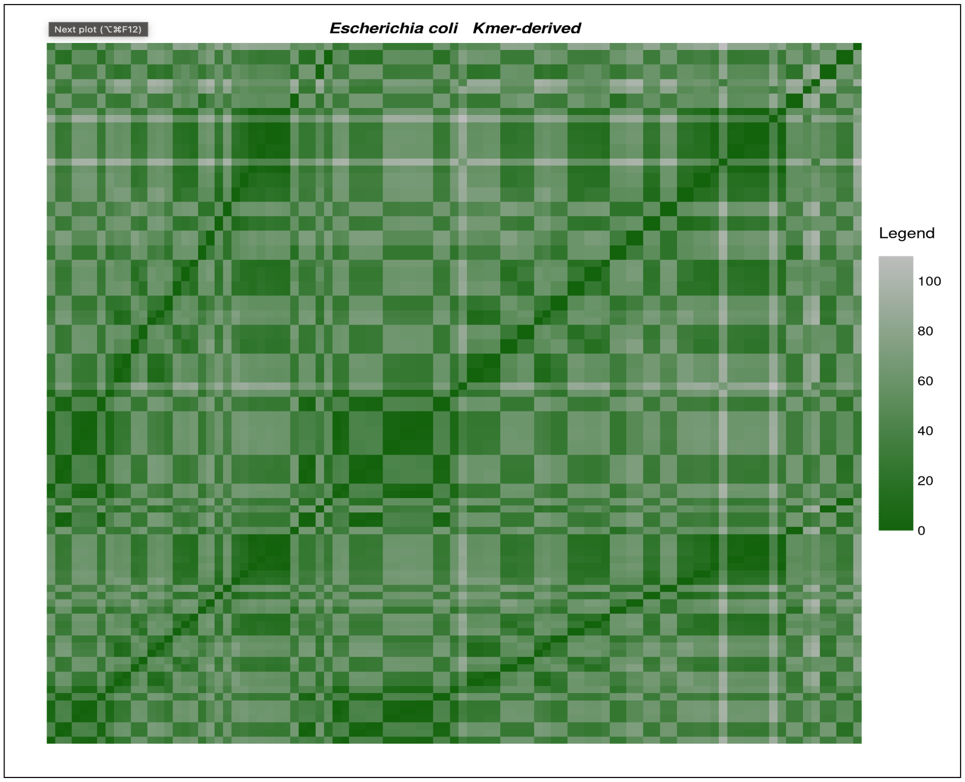


**Fig. 16** A distance matrix between reference operons of those operons observed and the ASVs for which these operons give optimal alignment for *Escherichia coli* in the mock microbiome. The distance measure is the approximate kmer measure.

Figure 17, perhaps fortuitously, identifies 6 clusters of operons and ASVs. Unfortunately the degree of clumping does not allow determination of which operons and ASVs are in most of those clumps.

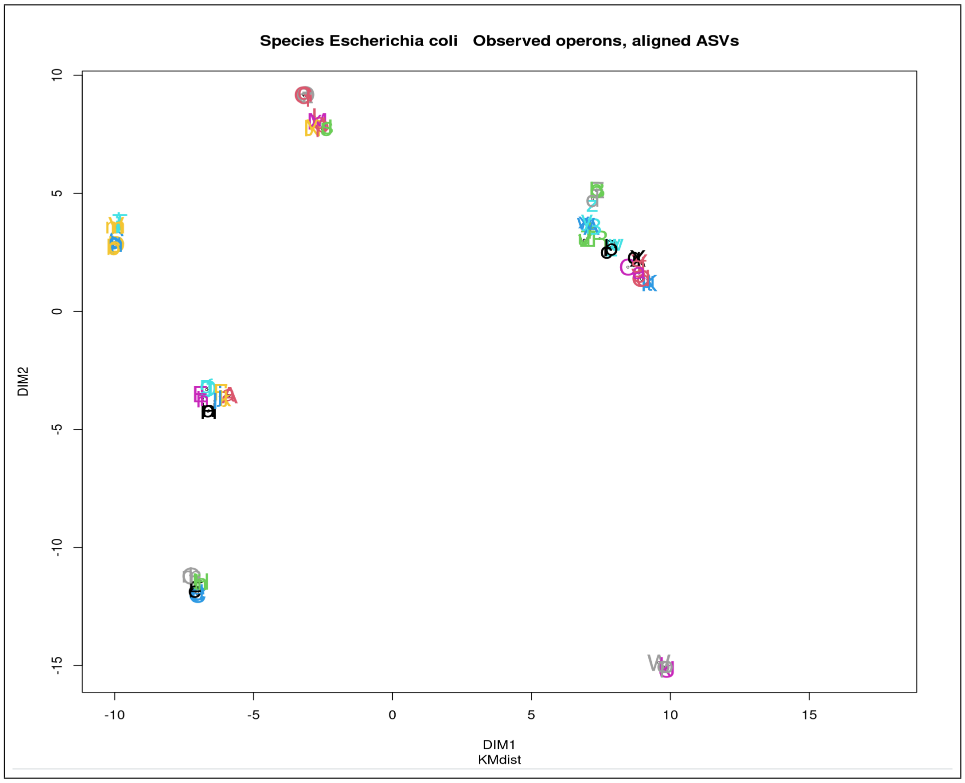


**Fig. 17** A UMAP2D projection plot for a distance matrix between those reference operons observed and the ASVs for which these operons give optimal alignment for *Escherichia coli* in the mock microbiome. The distance measure is the approximate kmer measure.

#### 4.4 *Bifidobacterium bifidum*

There are 3 operons and 8 (2,4,2, respectively, by strain) associated ASVs for *Bifidobacterium bifidum*. Table 5 gives the identities of the observed operons and some details on their ASV alignments.

**Table 5** Observed operons for *Bifidobacterium bifidum* , together with alignment details for the first-listed ASV of each operon

| Operon | Alen | MM | Gap | PID |
|---|---|---|---|---|
| PRL2010 | 5031 | 0 | 0 | 100 |
| S17 | 5033 | 0 | 0 | 100 |
| BGN4 | 5032 | 0 | 0 | 100 |

Note that for S17 the third and fourth listed ASVs have alignment lengths of 4559 and 4696, respectively, and each has 1 mis-match in the alignment while the fourth ASV has 1 gap also.

Figure 18 shows the very clear affect of these 2 ASVs that have poorer alignment.

Figure 19 provides a UMAP2D visualisation of the relation between the observed operons and their associated ASVs. This does not well reflect the distance matrix information in that it has the ASVs associated with operon PRL2010 (A) and BGN4 (C) dispersed almost as much from their operon as the ASVs for strain S17 are from the operons of S17(B).

For this species the evidence is quite clear that we get strain-level resolution. The UMAP2D representation provides limited support for that assertion. That is probably the consequence of the 2 poor ASVs.

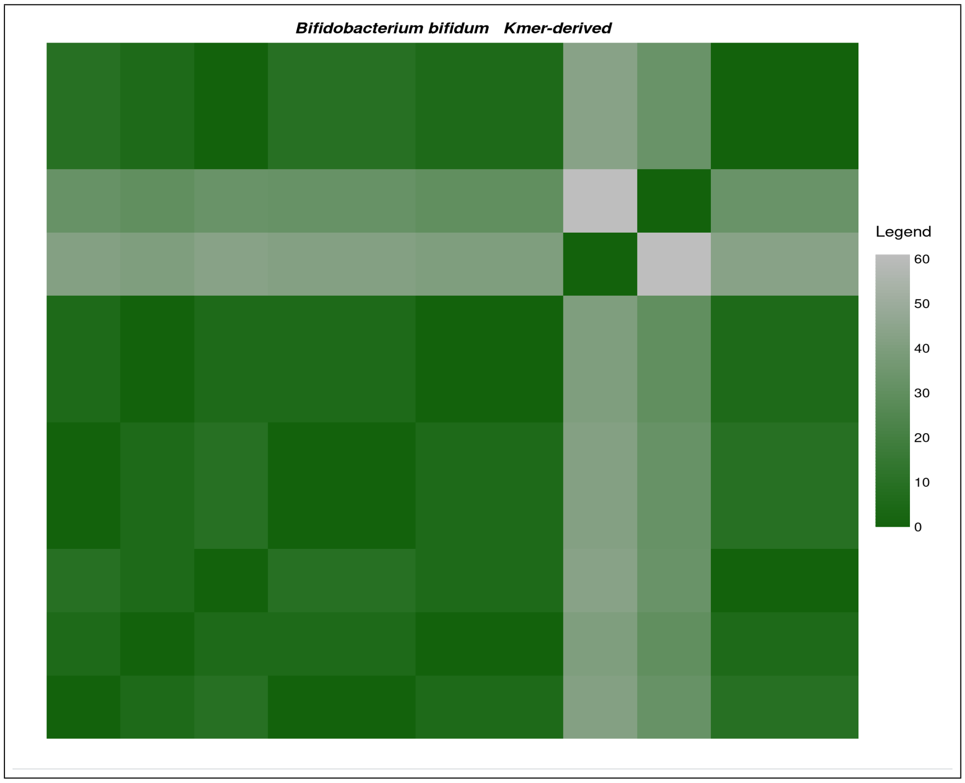


**Fig. 18** A distance matrix between reference operons of those operons observed and the ASVs for which these operons give optimal alignment for *Bifidobacterium bifidum* in the mock microbiome. The distance measure is the approximate kmer measure.

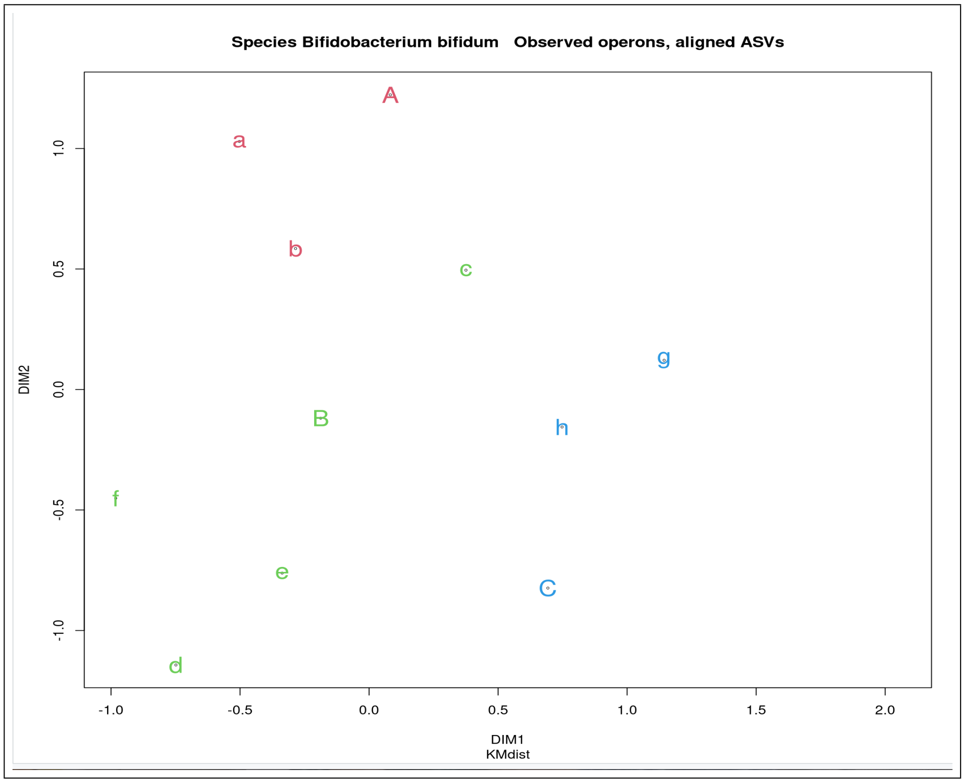


**Fig. 19** A UMAP2D projection plot for a distance matrix between those reference operons observed and the ASVs for which these operons give optimal alignment for *Bifidobacterium bifidum* in the mock microbiome. The distance measure is the approximate kmer measure.

### 5 Delivering improved strain level resolution via operons

Consider a partial re-thinking of the operon-led approach. If a particular strain is present then, if there are multiple operons of that strain that differ, this information could be used to help resolve ambiguity. How might this be done? Suppose a particular operon, O1, occurs in multiple strains in our DB. Note that we are interested in this operon because ASV alignment to the DB has identified multiple DB matches. If a different ASV identifies another operon, O2, from that one of those strains but not from some or all of the strains that had an O1 operon, then we consider the strain with the two operons, O1 and O2, identified from it to be present. Also, if we have ASV alignments to DB entries that do not have actual strain annotation but just subspecies, then, if there are also genuine strain level matches to the ASV, we should reject the subspecies-only matches.

So how would this work in practice?

#### 5.1 *Bifidobacterium longum*

Let us consider our B.longum analysis. There are 12 operons and we have the associated ASV indices for each of these. We are reporting all cases of optimal alignment for the ASVs, omitting non-strain level matches (such as *Bifidobacterium longum subsp. longum* . Thesse items should not have been in the reference DB which was meant to contain only operons with strain level annotation.

**Table 6** Detail on optimally aligned operons for *Bifidobacterium longum* . Note that alignments to operons not annotated to the strain level have been excluded from consideration. UM in the right-hand column means unambiguous match - that is, only 1 DB optimally matched operon occurs. Note the two ASVs , 227 and 290, that have shorter alignment lengths than others. Such alignments are less informative as they may align to more DB entries than ASVs with longer alignments. Column ASV identifies the matched ASV for the operon.

| Operon | ASV | Alen | PID | MM, Gap | Other Ops for this ASV? |
|---|---|---|---|---|---|
| GT15 Ops 1:5 | 54 | 5023 | 100 | 0, 0 | None for any |
| GT15 Ops 1:5 | 57 | 5023 | 100 | 0, 0 | None for any |
| GT15 Ops 1:5 | 227 | 4618 | 100 | 0, 0 | None for any |
| GT15 Ops 1:5 | 290 | 4634 | 100 | 0, 0 | None for any |
| JCM1217 Ops 1,3,4 | 191 | 5023 | 100 | 0, 0 | None for any |
| JCM1217 Ops 1,3,4 | 260 | 5023 | 100 | 0, 0 | None for any |
| DJO10A Op1 | 192 | 5022 | 100 | 0, 0 | None for any - UM |
| DJO10A Op1 | 210 | 5022 | 100 | 0, 0 | None for any - UM |
| DJO10A Op2 | 259 | 5024 | 100 | 0, 0 | None for any - UM |
| DJO10A Op2 | 329 | 5024 | 100 | 0, 0 | None for any - UM |
| DJO10A Op3 | 274 | 5023 | 100 | 0, 0 | None for any - UM |
| DJO10A Op4 | 289 | 5023 | 100 | 0, 0 | None for any - UM |
| JDM301 Op3 | 291 | 5021 | 100 | 0, 0 | None for any - UM |
| JDM301 Op3 | 368 | 5021 | 100 | 0, 0 | None for any - UM |
| JDM301 Op2 | 324 | 5021 | 100 | 0, 0 | None for any - UM |
| JDM301 Op2 | 418 | 5021 | 100 | 0, 0 | None for any - UM |
| JDM301 Op1 | 325 | 5020 | 100 | 0, 0 | None for any - UM |
| JDM301 Op1 | 330 | 5020 | 100 | 0, 0 | None for any - UM |
| ATCC 15697 Ops 1:4 | 367 | 5023 | 100 | 0, 0 | None for any |
| ATCC 15697 Ops 1:4 | 436 | 5023 | 100 | 0, 0 | None for any |
| 157F Ops 3, 4 | 388 | 5023 | 100 | 0, 0 | None for any |
| 157F Op 1 | 431 | 5023 | 100 | 0, 0 | None for any - UM |

We see here 6 strains identified, all of which are in the designed mock microbiome, and there are no missing strains. Note that for all ASVs optimal alignments were error-free, and in no case was there ambiguity of strain for any ASV. Not all operons are identified for every strain, but most are. Strain JDM301 has 3 operons, GT15 has 5 and the others have 4.

#### 5.2 *Bacteroides fragilis*

Let us consider our B.fragilis analysis. There are 11 operons and we have 40 associated ASV indices for each of these. We are reporting all cases of optimal alignment for the ASVs.

There are clearly 2 strains and none have been missed. Of the 5 ASVs that do not give perfect alignments 4 are short. ASVs 141 and 155 are unusually poorly matched to the best-matching operon.

**Table 7** Detail on optimally aligned operons for *Bacteroides fragilis* . Note that alignments to operons not annotated to the strain level have been excluded from consideration. UM in the right-hand column means unambiguous match - that is, only 1 DB optimally matched operon occurs. Note the two ASVs , 141 and 155 have markedly poorer alignments than all other ASVs listed. They also have less than 5% and 10%, respectively of the read count for their associated operon. Column ASV identifies the matched ASV for the operon.

| Operon | ASV | Alen | PID | MM, Gap | Other Ops for this ASV? |
|---|---|---|---|---|---|
| YCH46 Op5 | 13 | 4894 | 100 | 0, 0 | None for any - UM |
| YCH46 Op5 | 31 | 4894 | 100 | 0, 0 | None for any - UM |
| YCH46 Op5 | 121 | 4894 | 99.98 | 1 , 0 | None for any - UM |
| YCH46 Op5 | 133 | 4522 | 100 | 0, 0 | None for any - UM |
| YCH46 Op5 | 194 | 4384 | 100 | 0, 0 | None for any - UM |
| YCH46 Op5 | 277 | 4802 | 100 | 0, 0 | None for any - UM |
| YCH46 Op5 | 313 | 4483 | 100 | 1 , 0 | None for any - UM |
| YCH46 Op3 | 28 | 4971 | 100 | 0, 0 | None for any - UM |
| YCH46 Op3 | 36 | 4971 | 100 | 0, 0 | None for any - UM |
| YCH46 Op3 | 161 | 4347 | 100 | 0, 0 | None for any - UM |
| YCH46 Op4 | 29 | 4896 | 100 | 0, 0 | None for any - UM |
| YCH46 Op4 | 32 | 4896 | 100 | 0, 0 | None for any - UM |
| YCH46 Op4 | 141 | 4687 | 99.72 | 9,4 | None for any - UM |
| YCH46 Ops4,5 | 164 | 4463 | 99.98 | 1, 0 | None for any - UM |
| YCH46 Op2 | 30 | 4902 | 100 | 0, 0 | None for any - UM |
| YCH46 Op2 | 34 | 4902 | 100 | 0, 0 | None for any - UM |
| YCH46 Op2 | 177 | 4350 | 100 | 0, 0 | None for any - UM |
| YCH46 Op2 | 272 | 4630 | 100 | 0, 0 | None for any - UM |
| YCH46 Op1 | 35 | 4974 | 100 | 0, 0 | None for any - UM |
| YCH46 Op1 | 37 | 4974 | 100 | 0, 0 | None for any - UM |
| YCH46 Op1 | 174 | 4405 | 100 | 0, 0 | None for any - UM |
| YCH46 Op1 | 182 | 4710 | 100 | 0, 0 | None for any - UM |
| YCH46 Op1 | 295 | 4931 | 100 | 0, 0 | None for any - UM |
| YCH46 Op1 | 359 | 4737 | 100 | 0, 0 | None for any - UM |
| YCH46 Op1 | 378 | 4543 | 100 | 0, 0 | None for any - UM |
| YCH46 Op6 | 38 | 4894 | 100 | 0, 0 | None for any - UM |
| YCH46 Op6 | 353 | 4352 | 100 | 0, 0 | None for any - UM |
| YCH46 Op6 | 389 | 4686 | 100 | 0, 0 | None for any - UM |
| 638R Ops4,5 | 39 | 4905 | 100 | 0, 0 | None for any - UM |
| 638R Ops4,5 | 40 | 4905 | 100 | 0, 0 | None for any - UM |
| 638R Ops4,5 | 155 | 4524 | 99.76 | 9, 2 | None for any - UM |
| 638R Ops4,5 | 225 | 4548 | 99.98 | 1, 0 | None for any - UM |
| 638R Op1 | 41 | 4896 | 100 | 0, 0 | None for any - UM |
| 638R Op1 | 58 | 4896 | 100 | 0, 0 | None for any - UM |
| 638R Op2 | 60 | 4897 | 100 | 0, 0 | None for any - UM |
| 638R Op2 | 65 | 4897 | 100 | 0, 0 | None for any - UM |
| 638R Op3 | 61 | 4897 | 100 | 0, 0 | None for any - UM |
| 638R Op3 | 188 | 4897 | 100 | 0, 0 | None for any - UM |
| 638R Op6 | 62 | 4904 | 100 | 0, 0 | None for any - UM |
| 638R Op6 | 63 | 4904 | 100 | 0, 0 | None for any - UM |

#### 5.3 *Escherichia coli*

Let us now consider a similar E.coli analysis. There are 35 observed operons and 62 associated ASVs. Most operons have 2 associated ASVs, with 8 having only 1. Table 8 provides key data from the alignment of ASVs to the operons of the reference database.

**Table 8**: Detail on optimally aligned operons for *Escherichia coli*. Note that entries in the rightmost column for BL21-pLyS AG are too long and so have been assigned to labels defined in footnotes 1 to 4. Likewise for some later entries.

| Operon | ASV | Alen | PID | MM, Gap | Other Ops this ASV? |
|---|---|---|---|---|---|
| CFT073 Op4 | 77 | 4800 | 100 | 0, 0 | None for any - UM |
| CFT073 Op7 | 95 | 4892 | 100 | 0, 0 | None for any - UM |
| CFT073 Op7 | 113 | 4892 | 100 | 0, 0 | None for any - UM |
| CFT073 Op1 | 96, | 4892 | 100 | 0, 0 | None for any - UM |
| CFT073 Op1 | 104 | 4892 | 100 | 0, 0 | None for any - UM |
| CFT073 Op5 | 105, | 4800 | 100 | 0, 0 | None for any - UM |
| CFT073 Op5 | 126 | 4892 | 100 | 0, 0 | None for any - UM |
| CFT073 Op2 | 107 | 4800 | 100 | 0, 0 | Nissle 1917 Op3 |
| CFT073 Op2 | 111 | 4800 | 100 | 0, 0 | Nissle 1917 Op3 |
| CFT073 Op6 | 118 | 4800 | 100 | 0, 0 | Nissle 1917 Op5 |
| CFT073 Op6 | 169 | 4800 | 100 | 0, 0 | Nissle 1917 Op5 |
| CFT073 Op3 | 130 | 4892 | 100 | 0, 0 | None for any - UM |
| CFT073 Op3 | 134 | 4892 | 100 | 0, 0 | None for any - UM |
| UM146 Ops4, 6 | 78 | 4800 | 100 | 0, 0 | NU14 Op6, UTI89 Op7 |
| UM146 Ops4, 6 | 341 | 4439 | 100 | 0, 0 | NU14 Op6, UTI89 Op7 |
| SMS-3-5 Ops2, 6 | 80 | 4801 | 100 | 0, 0 | None for any - UM |

**Table 8**: (continued)

| Operon | ASV | Alen | PID | MM, Gap | Other Ops this ASV? |
|---|---|---|---|---|---|
| SMS-3-5 Ops2, 6 | 187 | 4801 | 100 | 0, 0 | None for any - UM |
| SMS-3-5 Ops5 | 224 | 4801 | 100 | 0, 0 | None for any - UM |
| SMS-3-5 Ops3 | 228 | 4890 | 100 | 0, 0 | None for any - UM |
| SMS-3-5 Ops3 | 429 | 4890 | 100 | 0, 0 | None for any - UM |
| SMS-3-5 Ops1 | 221 | 4883 | 100 | 0, 0 | None for any - UM |
| SMS-3-5 Ops1 | 305 | 4883 | 100 | 0, 0 | None for any - UM |
| SMS-3-5 Ops7 | 304 | 4800 | 100 | 0, 0 | None for any - UM |
| SMS-3-5 Ops4 | 321 | 4790 | 100 | 0, 0 | None for any - UM |
| BL21-pLysS AG Ops2,4,7 | 117 | 4793 | 100 | 0, 0 | Blist1[1] |
| BL21-pLysS AG Ops2,4,7 | 181 | 4793 | 100 | 0, 0 | Blist1 |
| BL21-pLysS AG Op3 | 258 | 4887 | 100 | 0, 0 | Blist2[2] |
| BL21-pLysS AG Op3 | 322 | 4887 | 100 | 0, 0 | Blist2 |
| BL21-pLysS AG Op6 | 387 | 4892 | 100 | 0, 0 | Blist3[3] |
| BL21-pLysS AG Op6 | 440 | 4892 | 100 | 0, 0 | Blist3 |
| BL21-pLysS AG Op1 | 407 | 4896 | 100 | 0, 0 | Blist4[4] |
| 1303 Ops4 | 170 | 4799 | 100 | 0, 0 | None for any - UM |
| 1303 Ops4 | 402 | 4799 | 100 | 0, 0 | None for any - UM |
| 1303 Ops2 | 287 | 4877 | 100 | 0, 0 | None for any - UM |
| 1303 Ops2 | 364 | 4877 | 100 | 0, 0 | None for any - UM |
| 1303 Ops5 | 362 | 4892 | 100 | 0, 0 | None for any - UM |
| 1303 Ops5 | 408 | 4892 | 100 | 0, 0 | None for any - UM |
| 1303 Ops6 | 365 | 4885 | 100 | 0, 0 | None for any - UM |
| 1303 Ops6 | 403 | 4885 | 100 | 0, 0 | None for any - UM |
| 1303 Ops3 | 366 | 4892 | 100 | 0, 0 | None for any - UM |

**Table 8**: (continued)

| Operon | ASV | Alen | PID | MM, Gap | Other Ops this ASV? |
|---|---|---|---|---|---|
| 1303 Ops3 | 441 | 4892 | 100 | 0, 0 | None for any - UM |
| 1303 Ops1 | 385 | 4883 | 100 | 0, 0 | None for any - UM |
| 1303 Ops1 | 430 | 4883 | 100 | 0, 0 | None for any - UM |
| UMNK88 Ops3 | 179 | 4894 | 100 | 0, 0 | None for any - UM |
| UMNK88 Ops1,3 | 349 | 4402 | 100 | 0, 0 | None for any - UM |
| UMNK88 Ops5 | 180 | 4894 | 100 | 0, 0 | None for any - UM |
| UMNK88 Ops5 | 186 | 4894 | 99.96 | 2, 0 | None for any - UM |
| UMNK88 Ops6 | 223 | 4894 | 100 | 0, 0 | None for any - UM |
| UMNK88 Ops6 | 347 | 4894 | 100 | 0, 0 | None for any - UM |
| UMNK88 Ops7 | 271 | 4894 | 100 | 0, 0 | None for any - UM |
| UMNK88 Ops1,6,7 | 422 | 4379 | 100 | 0, 0 | None for any - UM |
| UMNK88 Ops1 | 306 | 4894 | 100 | 0, 0 | None for any - UM |
| UMNK88 Ops4 | 346 | 4800 | 100 | 0, 0 | None for any - UM |
| UMNK88 Ops4 | 406 | 4800 | 100 | 0, 0 | None for any - UM |
| Nissle 1917 Ops1,2,5,6 | 337 | 4583 | 100 | 0, 0 | None for any - UM |
| Nissle 1917 Ops6 | 342 | 4488 | 100 | 0, 0 | None for any - UM |
| REL606 Op6 | 345 | 4804 | 100 | 0, 0 | Blist5[5] |
| REL606 Op6 | 442 | 4804 | 100 | 0, 0 | Blist5 |
| NU14 Op3 | 363 | 4892 | 100 | 0, 0 | UTI89 Op3 |
| NU14 Op3 | 401 | 4892 | 100 | 0, 0 | UTI89 Op3 |
| O18:H1 Op5 | 396 | 4487 | 100 | 0, 0 | CFTList[6] |

[1] REL606 Ops2,4,7;B Ops1,4,6; BL21 2,4,7
[2] B Op5; BL21 Op 1 (BL21(DE3) Op 1)
[3] B Op2; REL606 5; BL21 Op5 (BL21(DE3) Op5)
[4] B Op7; REL606 3; BL21 Op3 (BL21(DE3) Op3)
[5] B Op3; BL21(DE3) Op6; BL21 Op6
[6] CFT073 Op6, Nissle 1917 Op7 (Alen 4487)

We see that CFT073, SMS-3-5, 1303, and UMNK88 are cleanly resolved. Nissle 1917 has 2 operons that give length 4800 alignments that are optimal matches to the same ASVs as 2 operons of CFT073. Additionally Nissle 1917 has 1 ASV that aligns unambiguously to operons 1,2,5 and 6 but with a shorter alignment length of 4583, and 1 ASV that aligns unambiguously to operon 6 with alignment length of 4487. A somewhat risky call would be that Nissle 1917 strain is present. Such a call would be based on two relatively short alignments ( $\sim 4500$ bases) and claiming precedence for Nissle 1917 over 2 operons that are more likely from CFT073.

UM146, NU14 and UTI89 compete for a pair of ASVs(78, 341), while NU14 and UTI89 compete for the pair (363, 401) of ASVs. There are 2 identical UM146 operons aligning to ASVs (78,341) with Alen of 4800. The most reasonable decision is that either or both of NU14 and UTI89 are present.

There is clearly considerable overlap between the various B strains, and REL606 is part of this. The ordering of operons - which corresponds to the ordering on the genome - shows that B is the reverse of AG (and all 7 operons are matched) - see Table 9 which brings together the information for this analysis in terms of which operons are identical. It also shows that REL606, BL21 and BL21(DEL) are identical as far as the rrn operons are concerned, and their ordering of operons does not match that of AG or B. Thus we conclude that BL21 and BL21(DEL) are indistinguishable, and since all 7 operons are matched by ASVs for B, BL21 and BL21(DEL) any one of those 3 strains could be the most likely of these present.

The entry for O18:H1 has a relatively short alignment and is ambiguous with CFT073 and Nissle 1917. Given that CFT073 is considered definitely present, this ASV is treated as being associated with operon 6 of CFT073.

**Table 9** Operon indices for identical operons of B-type strains of *Escherichia coli*

| BL21...AG | REL | B | BL21 | BL21(DEL) |
|---|---|---|---|---|
| 1 | 3 | 7 | 3 | 3 |
| 2 | 2 | 6 | 2 | 2 |
| 3 | - | 5 | 1 | 1 |
| 4 | 4 | 4 | 4 | 4 |
| 5 | - | - | - | - |
| 6 | 5 | 2 | 5 | 5 |
| 7 | 7 | 1 | 7 | 7 |
| - | 6 | 3 | 6 | 6 |

Following the above we would consider the following to be the identified E.coli strains:-

- CFT073
- SMS-3-5
- 1303
- UMNK88
- one of (B, BL21, BL21(DE3))
- one of (NU14,UTI89).

The designed strains are *Escherichia coli CFT073, Escherichia coli SMS-3-5, Escherichia coli UMNK88, Escherichia coli 1303, Escherichia coli BL21(DE3),*and *Escherichia coli UTI89.*

Here we see that, even for a discrimination task as difficult as this one, this analysis approach has proven to be quite effective in resolving E.coli strains, except for cases of multiple identical sets. For the B strains we were also able to reduce ambiguity by using the ordering of operons on the genome as implied by their operon numbering in the database. This highlights the importance of having strain databases that include all the operons of that strain, indexed according to genomic location order.

#### 5.4 *Bifidobacterium bifidum*

Let us consider our *Bifidobacterium bifidum* analysis. There are 11 operons and we have 40 associated ASV indices for each of these. We are reporting all cases of optimal alignment for the ASVs. Table 10 presents detail on each ASVs optimal alignments. We see that there is no strain ambiguity at the operon level and for each of the 3 strains all unique operons are identified. Figure 19 does not give such a clear indication of strain resolution, probably because of the short alignment of ASV 340 and its impact on the inter-operon, inter-ASV distance matrix underpinning the UMAP model generation.

**Table 10** Detail on optimally aligned operons for *Bifidobacterium bifidum* . Note that alignments to operons not annotated to the strain level have been excluded from consideration. UM in the right-hand column means unambiguous match - that is, only 1 DB optimally matched operon occurs. Column ASV identifies the matched ASV for the operon.

| Operon | ASV | Alen | PID | MM, Gap | Other Ops for this ASV? |
|---|---|---|---|---|---|
| PRL2010 Ops1,2,3 | 97 | 5031 | 100 | 0, 0 | None for any - UM |
| PRL2010 Ops1,2,3 | 102 | 5031 | 100 | 0, 0 | None for any - UM |
| S17 Ops1,2,3 | 127 | 5033 | 100 | 0, 0 | None for any - UM |
| S17 Ops1,2,3 | 163 | 5033 | 100 | 0, 0 | None for any - UM |
| S17 Ops1,2,3 | 340 | 4559 | 99.98 | 1, 0 | None for any - UM |
| S17 Ops1,2,3 | 372 | 5033 | 99.96 | 1, 1 | None for any - UM |
| BGN4 Op 2 | 190 | 5032 | 100 | 0, 0 | None for any - UM |
| BGN4 Op 2 | 275 | 5032 | 100 | 0, 0 | None for any - UM |

### 6 Anomalous reads and their analysis

The generation of ASVs requires a sufficient number of closely related reads to allow estimation of what might be the common original sequence from which these reads were derived. The default minimum number in the code for the Robust Amplicon Denoising (RAD) code is five. We have not altered this default value in our work. As a result it is expected that reads that are from a strain with very low relative abundance will not have an ASV generated that corresponds to the sequence from which operons of that strain were derived. Such reads may end up being associated with ASVs of other strains. This results in ASVs that have reads from more than one strain associated with them. With simulated reads we can gain insight into this aspect of the processing, and examine the feasibility of further processing that exploits reads that would otherwise be lost.

We first introduce the concept of ASV purity. After identifying a consensus sequence for a cluster of reads and finalising the clustering process, RAD returns these consensus sequences, which ideally are exactly the sequence from which the reads of the associated cluster are derived. Such a sequence is labelled a template by the RAD developers ([8]), and would be an ASV if it aligns exactly with an entry in the database. As noted earlier we use the term ASV to denote the consensus sequence if it gives an alignment of acceptable quality, rather than requiring 100% identity. By including the strain and an operon index of the sequence from which it was generated in the header of each simulated read, it is possible to use the indexing of sequences associated with the reads in an ASV-associated cluster to determine the strain and operon index of any such ASV-associated read. It is therefore possible to specify what strains are represented within the set of reads associated with any ASV. Ideally, if one is seeking strain-level resolution, each ASV would have reads from only one strain in their associated reads. Such an ASV we consider to have an ASV strain purity of 1. If an ASV has more than one strain in its associated reads its purity is defined as the proportion of reads that are from the most common strain.

Now consider what we call anomalous reads. A read is anomalous if it is an outlier in terms of its edit distance from the ASV with which it is associated. Generally this would only occur for reads of a different strain, but that is not always the case. So it can occur that, for instance, 6 Salmonella operons are accepted while the 7th, distant about 100 bases away (edit distance), is treated as anomalous. A kmer-based calculation of edit distance from the ASV itself to each of the reads associated with a particular ASV allows a simple search for outliers. We use a threshold on upward distance from the ASV of the median distance plus 10 times the median absolute deviation from the median of all reads associated with the ASV.

#### 6.1 Processing anomalous reads

All anomalous reads identified across all ASVs were aligned against a chosen reference database. We use a blast database of 25285 ribosomal RNA operons having strain-level information, derived from the GROND Refseq207 database of Walsh et al. ([9]). The best blast hit criterion (BBH) is used to identify a match, subject to that hit having an acceptable length of alignment and alignment score.

All reads having an acceptable alignment are then used to adjust the results from the ASV alignment data. Firstly, the count of anomalous reads identified from a particular ASV is deducted from the count for that ASV. Then, the identified BBH for each read is checked to see if it corresponds to the strain label of any ASV. If so, that read is added to the count of such an ASV. The reads remaining after this are then considered as identifying strains additional to those already observed. The unique strains are identified, counted, and - after determining the number of rRNA operons of the particular strains - strain and species cellular counts are formed. These strains and species counts are then merged with the strain and species data from the adjusted earlier ASV analysis and cellular relative abundances are calculated.

#### 6.2 Results from processing the `mKB_rrn_C11` anomalous reads

The identification process for anomalous reads gave 182 such reads. These reads were processed by blastn to identify the BBH for each. Processing of blastn output identified 25 unique genera, 38 unique species, 44 unique strains and 79 unique unique operons. Because this is a mock microbiome with simulated reads whose headers included information on the operon from which they were generated, we can identify which of the strains identified are actually in the mock microbiome. We identified 40 strains that were valid and 4 that were mis-reports of presence. After removing anomalous reads from strains, and adding them to the already observed strains (valid or invalid) 11 of the 44 strains were found to have their read counts changed. The 6 strains with the largest counts of anomalous reads all had their count reduced (by a maximum of 9 counts for *Streptococcus mitis B6*, from 46 to 35), while the largest increase was from 9 to 13 for *Porphyromonas asaccharolytica DSM 20707*. Proportionately, the impact of non-zero adjustment in counts was greater for the increases than for the decreases.

After the adjustments there were 13 reads that had not been assigned to an observed strain. These best aligned to operons from *Bifidobacterium breve JCM7019*and

*Mageeibacillus indolicus UPII9-5* , both of which actually were in the designed microbiome. Hence the use of anomalous reads, as has been described, allows both improved accuracy for some strains through the re-assignment of reads, as well as, potentially, identification of strains that would otherwise be missed as they were present at too low a level to meet the threshold for ASV formation. Figure 33 provides a scatterplot of observed vs. designed strain cellular relative abundances that incorporates results from processing of anomalous reads.

In the real-world case we do not have the origin of the reads available. However the process above remains valid as it does not require any prior information on the origin of reads. We used such information above only for diagnostic purposes. When there is no ability to reference the origin of any reads there is no way to specify that some reported strains are valid while others are mis-reports. However, some guidance on the reliability of results achieved could be gained from checking the quality of read alignment - if it is an alignment of length similar to the *rrn* length, and has a PID close to 100 then it is probable that identification is at least highly likely to be correct at the species level.

##### 6.2.1 Results from processing the `mKB_rrn_C11` anomalous reads

The results of using this method for the `mKB_rrn_C11` anomalous reads are shown in Figure 20.

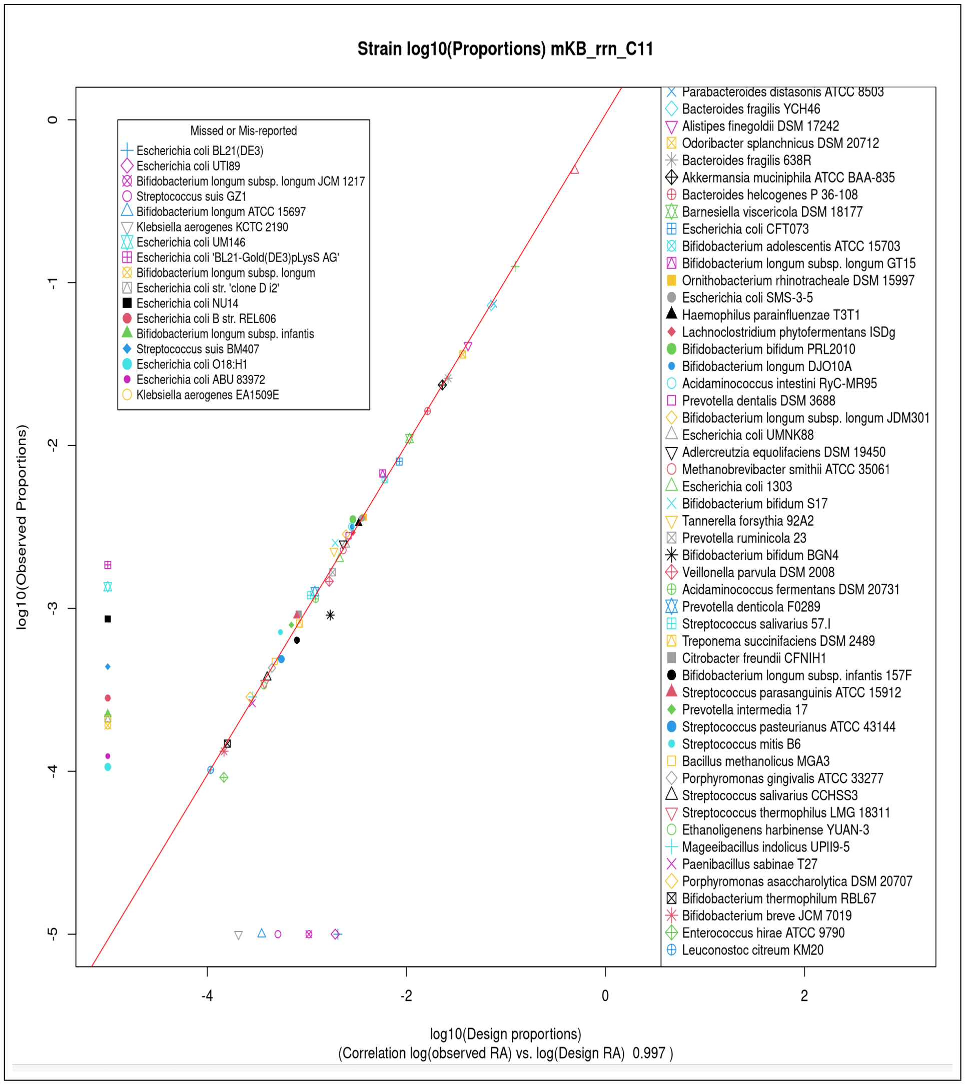


**Fig. 20** Scatterplot of observed vs. designed strain cellular relative abundance for dataset `mKB_rrn_C11` with the anomalous reads results incorporated. The red lines show a linear model fit to the data not including missed or invalid observations.

#### 6.3 UMAP 2D Visualisation of ASVs and their associated reads.

Because of the complexity of a visualisation of the full set of ASVs from the denoising we present both the full set and then multiple subsets of the data. Specifically, 4 of the subsets are of single species that have multiple strains in the mock microbiome. UMAP 2D projection coordinates were computed using the uwot package in R, with training based on all reads associated with any ASV, subject to a threshold of 30 such associated reads. For those reads in excess of the threshold, uwot has an embedding process which was used here to allow representation of all ASV-associated reads. Every ASV has been marked firstly by a black "+" marker. Additionally, a larger marker, with colouring as specified in the legend, is then placed for each ASV. This allows differentiation of strains from this figure.

Figures 21 and 22 show the dominance of the most abundant strain, *Phocaeicola vulgatus* in the microbiome. Those strains that have more than 30 reads associated with ASVs from that strain have embedded reads (green) represented. It is worth noting that only 30-60 of about 140,000 reads are not associated with any ASV. These, of course, may give rise to either additional - typically very low relative abundances - observed strains, or to mis-reported strains.

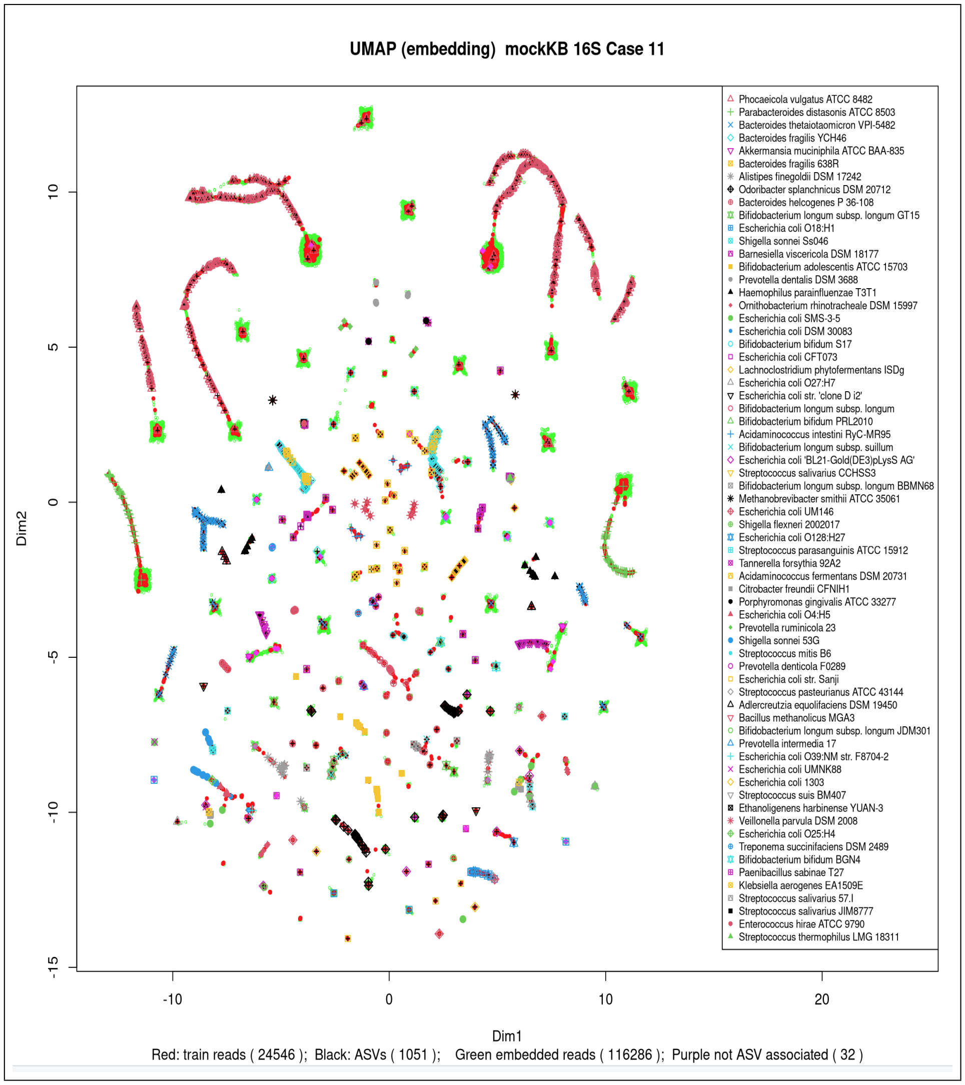


**Fig. 21** 2D UMAP representation of the relationship between ASVs and their associated read clusters for the 16S rRNA gene. Embedding can be seen for each of the first 6 strains listed in the legend - the most abundant strains - dominated by *Phocaeicola vulgatus* .

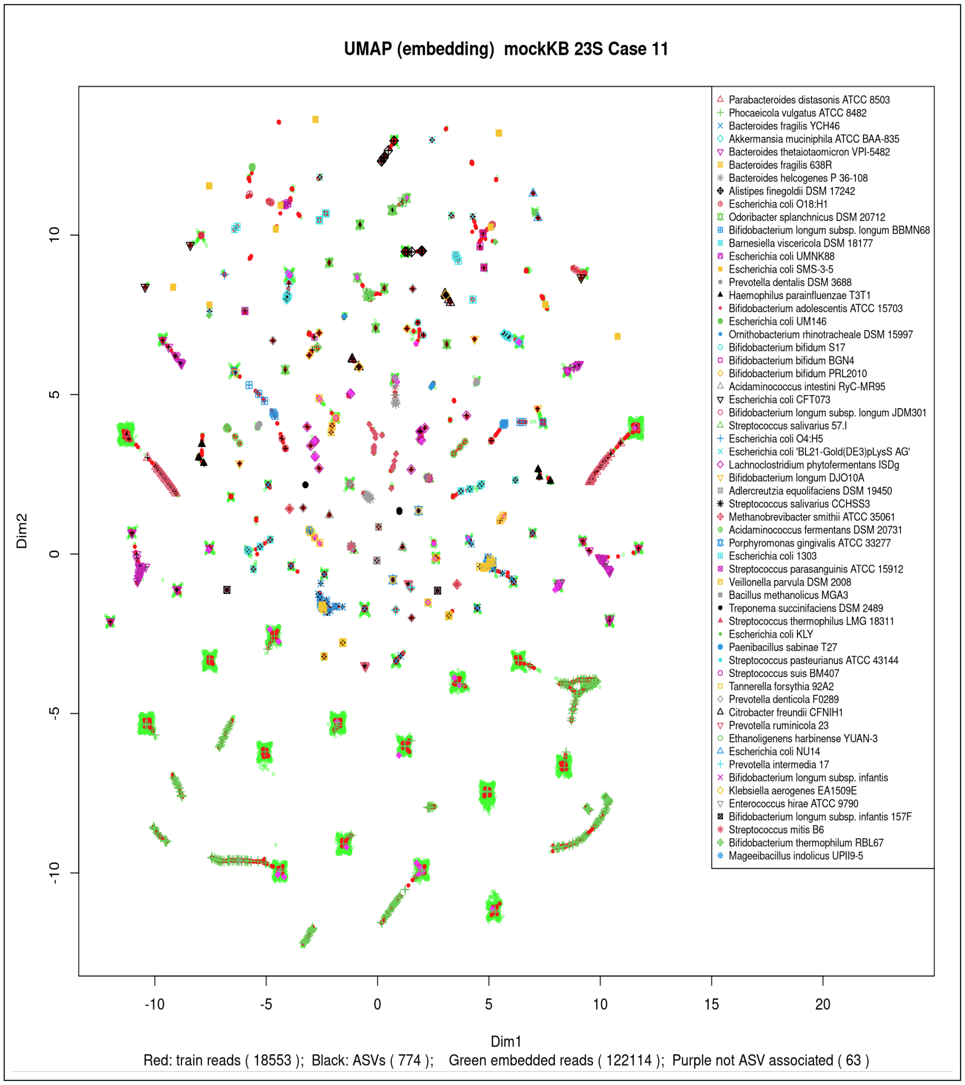


**Fig. 22** 2D UMAP representation of the relationship between ASVs and their associated read clusters for the 23S rRNA gene.

Separate UMAP representations were used for the 6 most abundant strains (Figure 23), and for four species that have multiple strains present *Bifidobacterium longum, Bifidobacterium bifidum, Bacteroides fragilis* and *Escherichia coli* (see Figures 24, 25, 26, 27), . Note, however, that the data for the plot of the top 6 relative abundance strains came from a smaller dataset (C10), generated with the same quality settings but approximately $\frac{2}{3}$ as many reads for denoising. This was because computation with the larger dataset did not complete in an acceptable time (48 hrs with 56 threads).

The patterns for the top 6 relative abundance strains from the 16S and 23S rRNA genes are broadly similar. However note that strain *Alistipes finegoldii DSM17242* occurs in the 16S plot but is replaced by *Eshcerichia coli O18:H1*in the 23S plot. There are also more reads somewhat distant from any ASV in the 16S plot. With a smaller library of reads it is the case that fewer strains and species are observed.

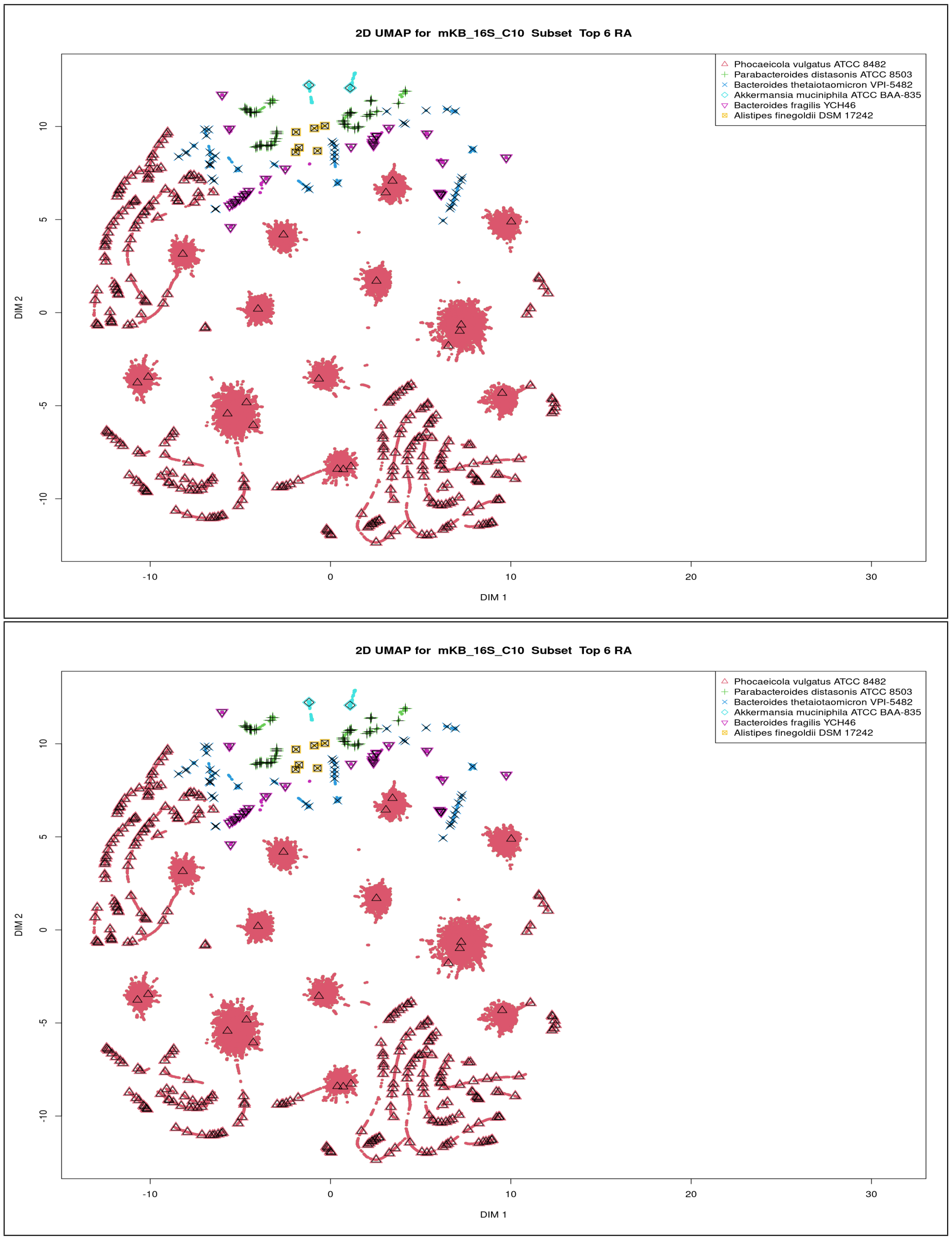


**Fig. 23** 2D UMAP representation of the relationship of the top 6 in RA strains for the 16S (left) and 23S rRNA gene.

Figure 24 shows just the *Bifidobacterium longum* strains in 2D UMAP plots for the 16S and 23S rRNA genes. These plots suggest better discrimination between strains with the 23S rRNA gene than with the 16S rRNA gene..

The *Bifidobacterium longum* species has 6 strains in the mKB mock microbiome. Two of those - *GT15* and *JDM301* - are identified in the 16S plot of Figure 24 and only one - *JDM301* - in the 23S plot. A common mis-reported strain for both 16S and 23S is *BBMN68* . Details from the blast alignment output shows that, for the 23S dataset, there were 9 ASVs that returned *BBMN68* as the best alignment, all these ASVs giving perfect alignments, with the ASVs differing in length (from 2189 to 2817). However operons of this strain and of multiple other operons of *Bifidobacterium longum* , strains also returned perfect alignments. Likewise for the 16S ASVs - 2 ASVs aligned to the *BBMN68* reference operons with PID equal to 100, but also to *DJO10A* (which is in mKB), as well as *NCC2705, sub-species infantis* , strain *infantis 157F*, and *KACC 91563* which are not in mKB. Thus lack of good strain-level resolution for this species is due to the 16S and 23S ribosomal RNA genes not having the diversity to allow resolution, rather than the method of analysis of sequenced rRNA gene data.

The reference database has a considerable number of operons that are not specified with any more taxonomic resolution than *Bifidobacterium longum subsp. longum* or *Bifidobacterium longum subsp. infantis* . Hence we see here for this species that neither the 16S or 23S rRNA gene has sufficient resolution for good strain resolution. It is of interest that this limitation has given rise to concerns over probiotics that claim to have certain beneficial effects due to a particular strain of *Bifidobacterium longum* but the specific strain was found to not be present in some commercial formulations claiming the beneficial effect [10], [11].

For both 16S and 23S rRNA genes the strains reported are identical and all three strains are in mKB. A cluster of *Bifidocbacterium bifidum S17* reads stand distant from the 2 ASVs aligned to that strain in both the 16S and 23S plots, and the 23S plot has 2 read clusters that do not sit close to an ASV. Despite this we are clearly seeing strain level resolution from both 16S and 23S rRNA data.

Two strains of *Bacillus fragilis* are present in mock microbiome mKB. The 16S analysis had one ASV that aligned best to strain NCTC 9343 with PID of 100, and had a single mis-match in alignment to an operon of strain 638R. No such problem arises with the 23S data. There are 14 reads associated with this ASV.

Not surprisingly *Escherichia coli* provides a demanding test for strain level resolution of the 4 species considered here, due to the number of strains actually in the designed microbiome.

The 16S data looks to have serious mis-reporting. However each of the mis-reports arises from ASVs that have PID values of 100, and there are correct strains also with that PID value. Thus the 16S cannot resolve which of multiple strains is present. The 23S has fewer mis-reports - 4 mis-reports and 6 correct identification. Again mis-reports result from tied best matches.

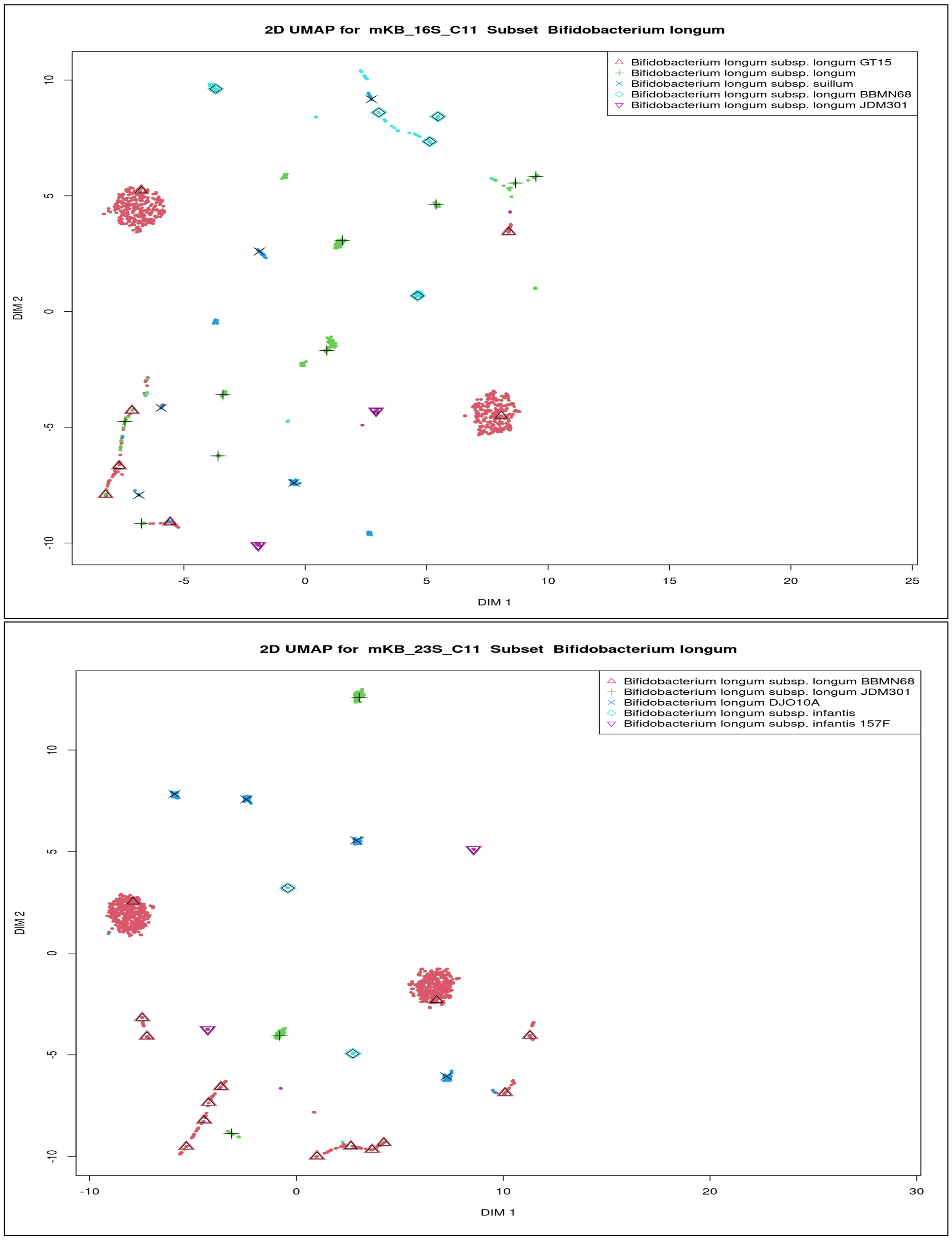


**Fig. 24** 2D UMAP representation of *Bifidobacterium longum* strains for 16S and 23S rRNA genes.

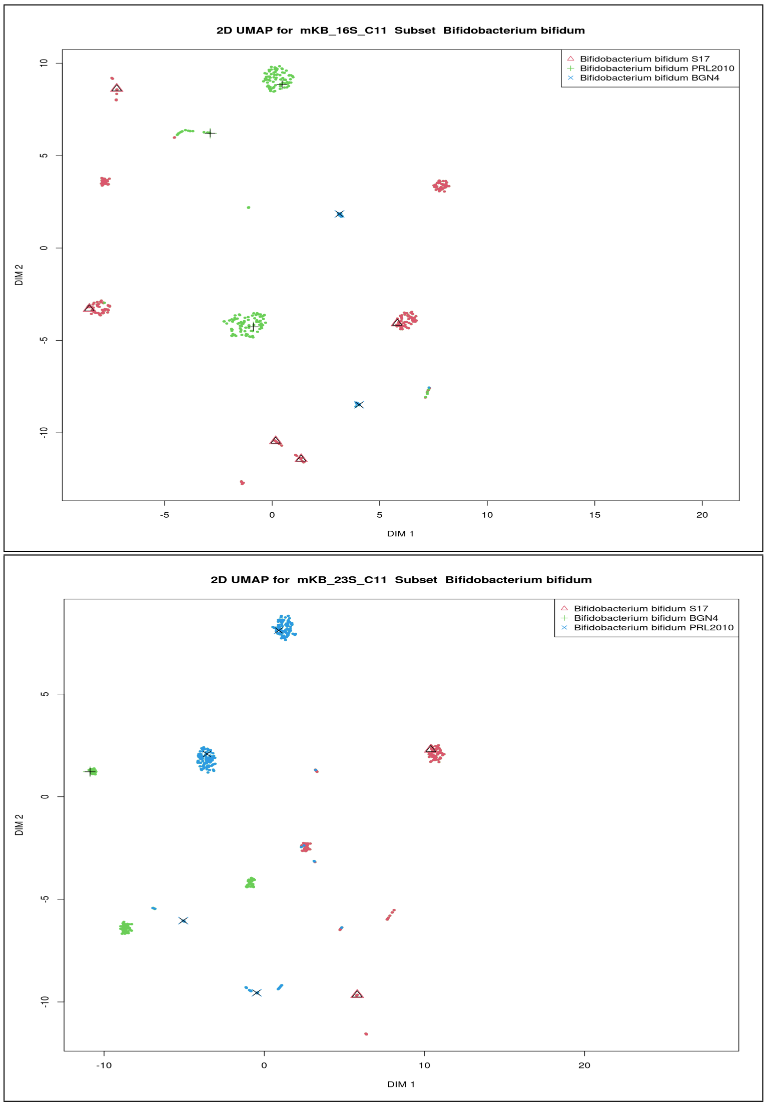


**Fig. 25** 2D UMAP representation of *Bifidobacterium bifidum* strains for 16S and 23S rRNA genes.

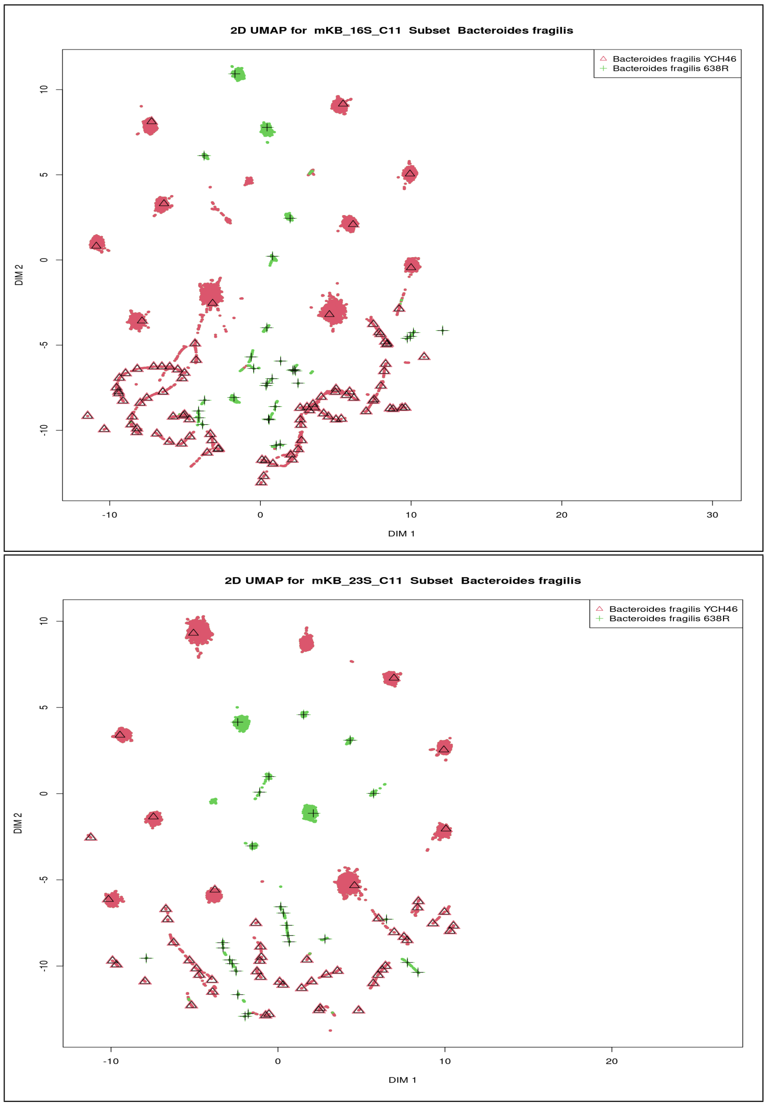


**Fig. 26** 2D UMAP representation of *Bacteroides fragilis* strains for 16S and 23S rRNA genes.

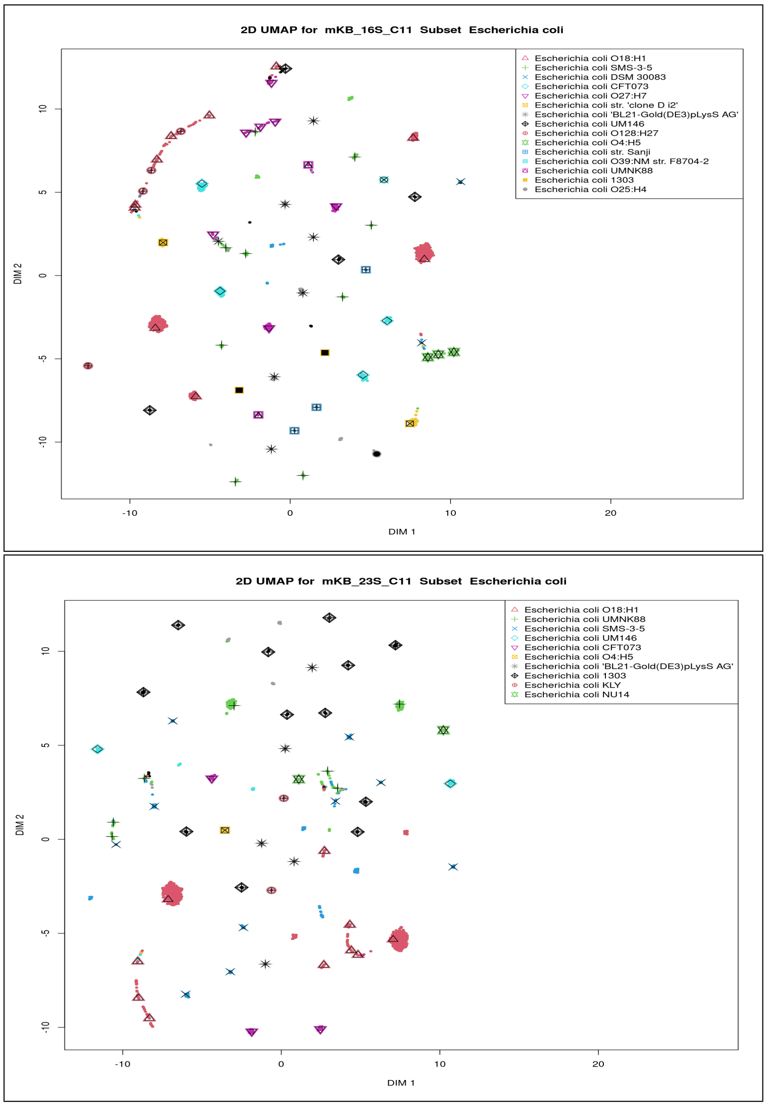


**Fig. 27** 2D UMAP representation of *Escherichia coli* strains for 16S and 23S rRNA genes.

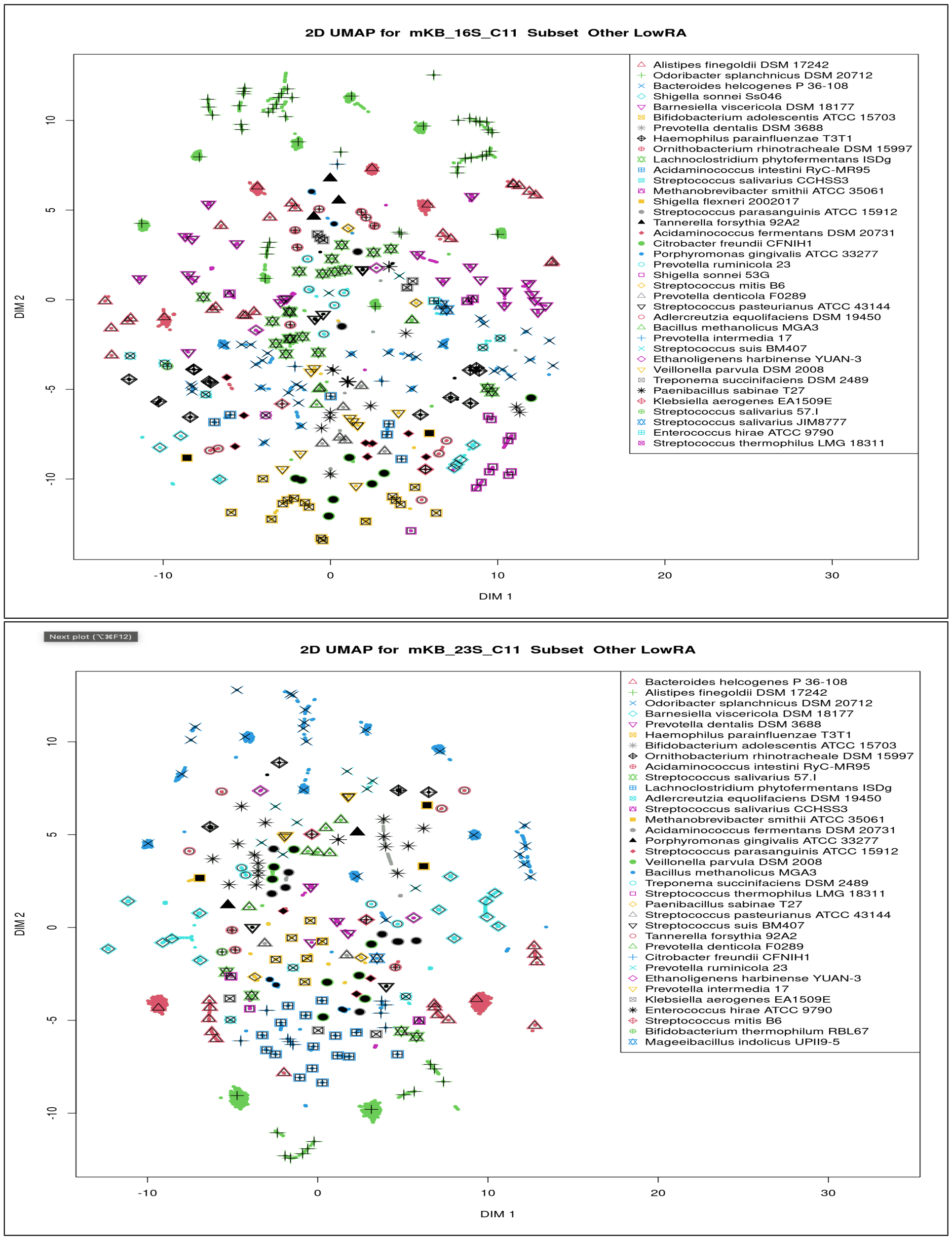


**Fig. 28** 2D UMAP representation, for 16S and 23S rRNA genes, of those strains not in the 6 most abundant, nor from the 4 species that were individually represented in figures 24 - 27.

#### 6.4 Relative Abundance - observed vs. actual

Here we present two slightly different sets of scatterplots. The first set - Figures 29 and 30 - has not included any results from the analysis of anomalous reads, whereas the second set - Figures 31 and 32 - does. Scatterplots are presented of the observed vs. actual relative abundances for the 16S and 23S datasets for the mKB microbiome, in the same manner as in the primary document. Note that the correlation coefficient reported does not include those strains which were either not observed, or were mis-reported as being present in the microbiome. It does include those strains deduced to be present based on the analysis of anomalous reads.

Figure 33 presents the corresponding result from the full ribosomal rRNA operon with anomalous read data included to allow easy comparison with the 16S and 23S ribosomal RNA genes.

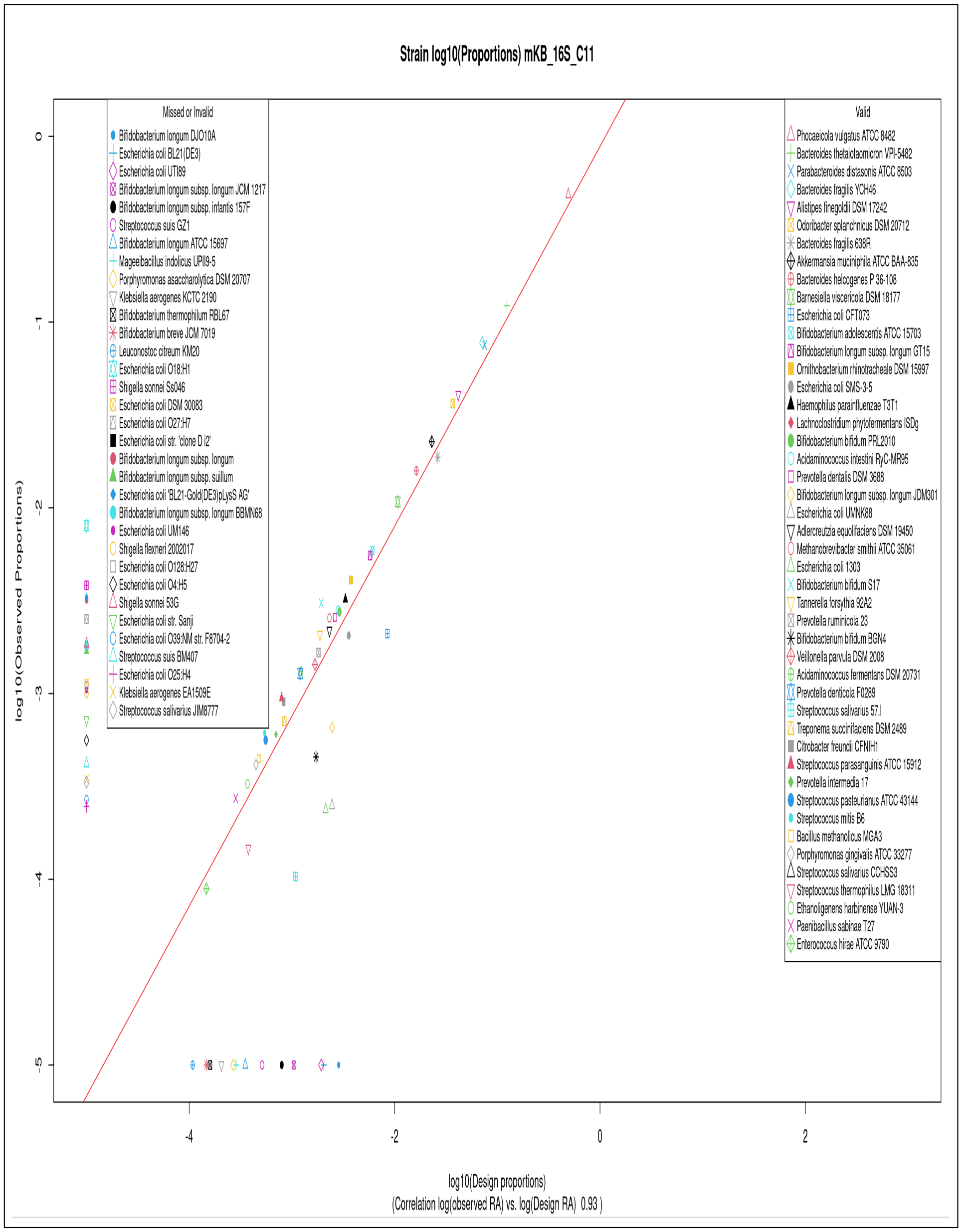


**Fig. 29** log(Observed) vs log(designed) relative abundances of the mKB microbiome as determined from 16S rRNA gene sequence reads from the C11 simulated nanopore-sequenced reads.

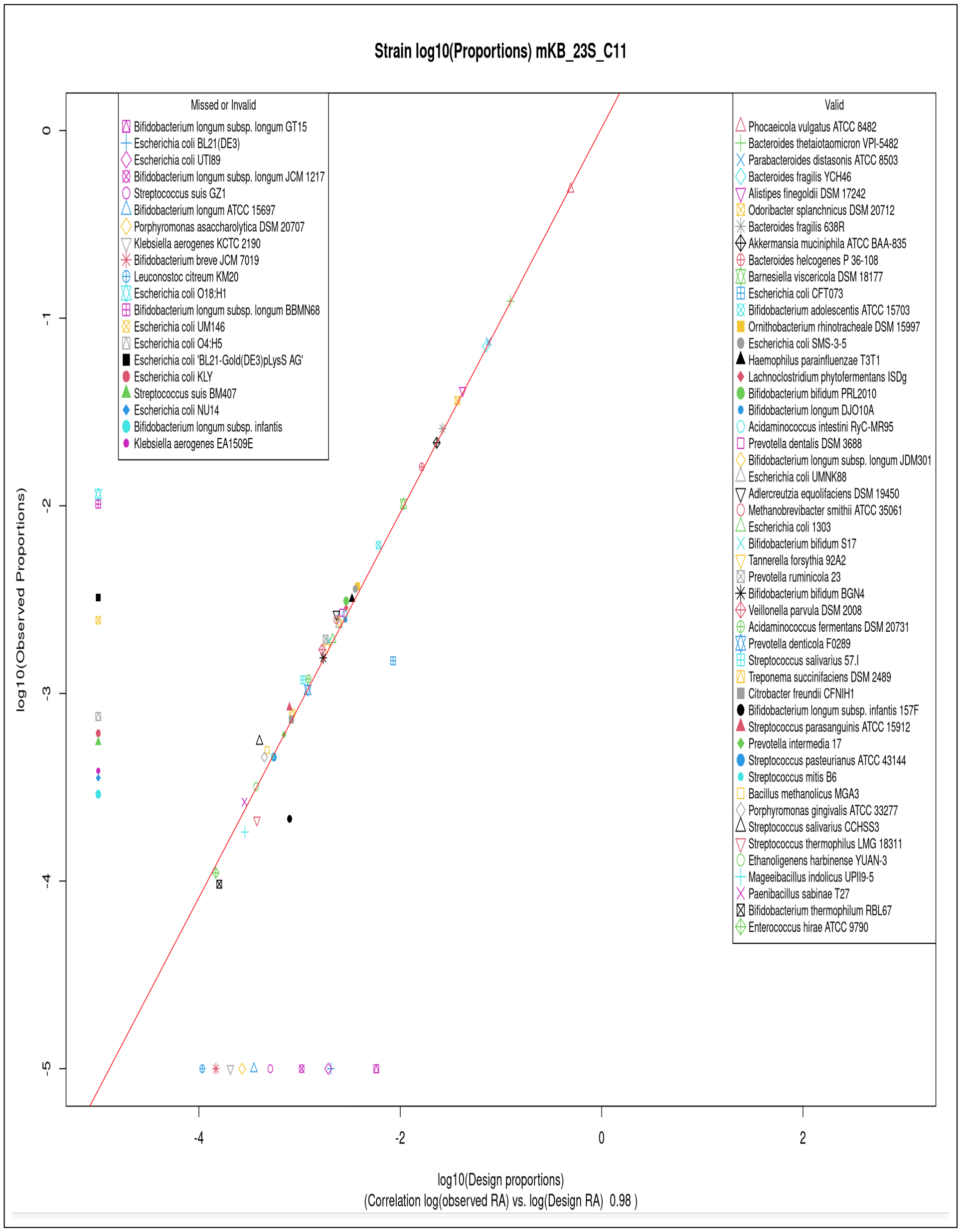


**Fig. 30** log(Observed) vs log(designed) relative abundances of the mKB microbiome as determined from 23S rRNA gene sequence reads from the C11 simulated nanopore-sequenced reads.

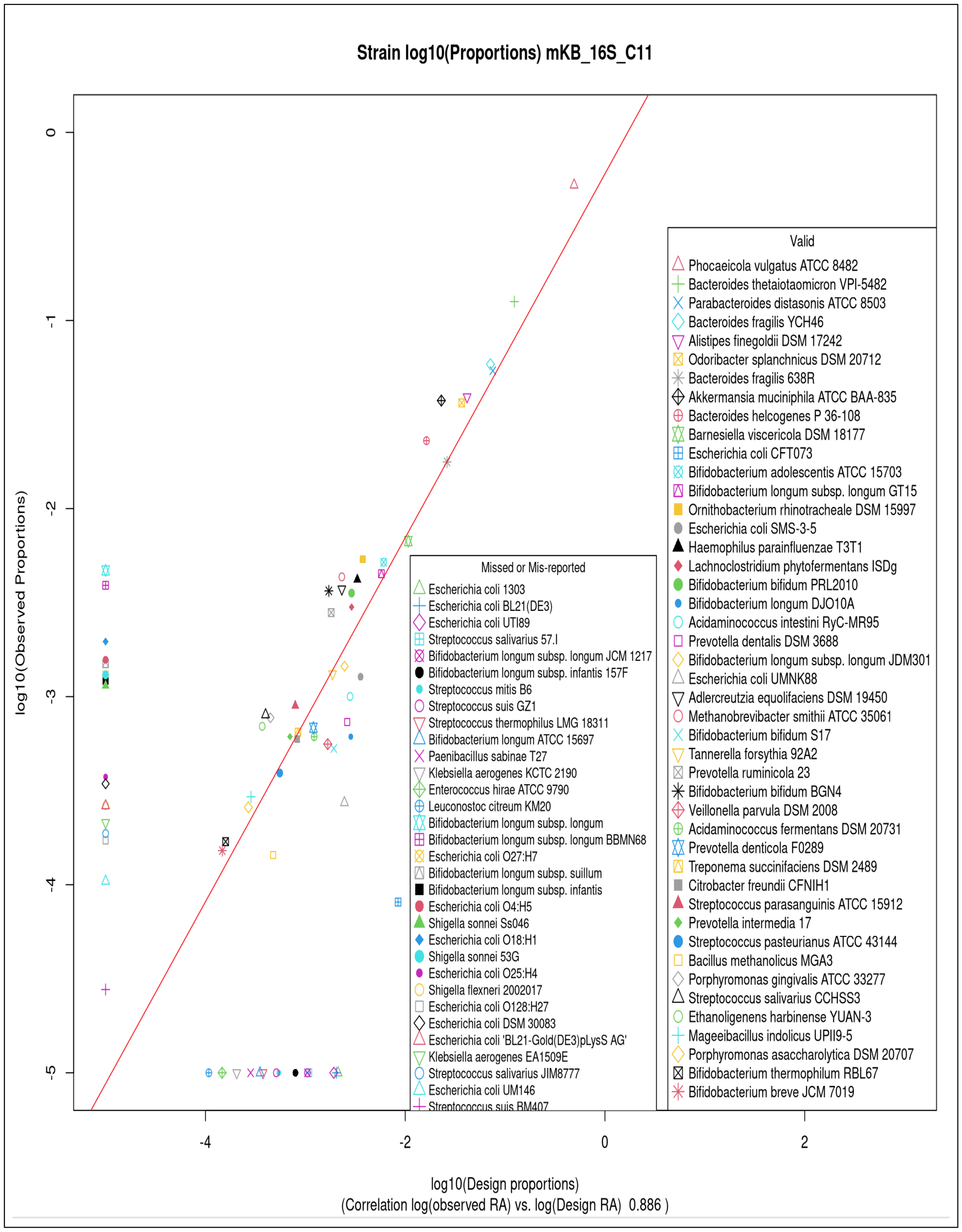


**Fig. 31** log(Observed) vs log(designed) relative abundances of the mKB microbiome as determined from 16S rRNA gene sequence reads from the C11 simulated nanopore-sequenced reads, with anomalous reads analysis incorporated.

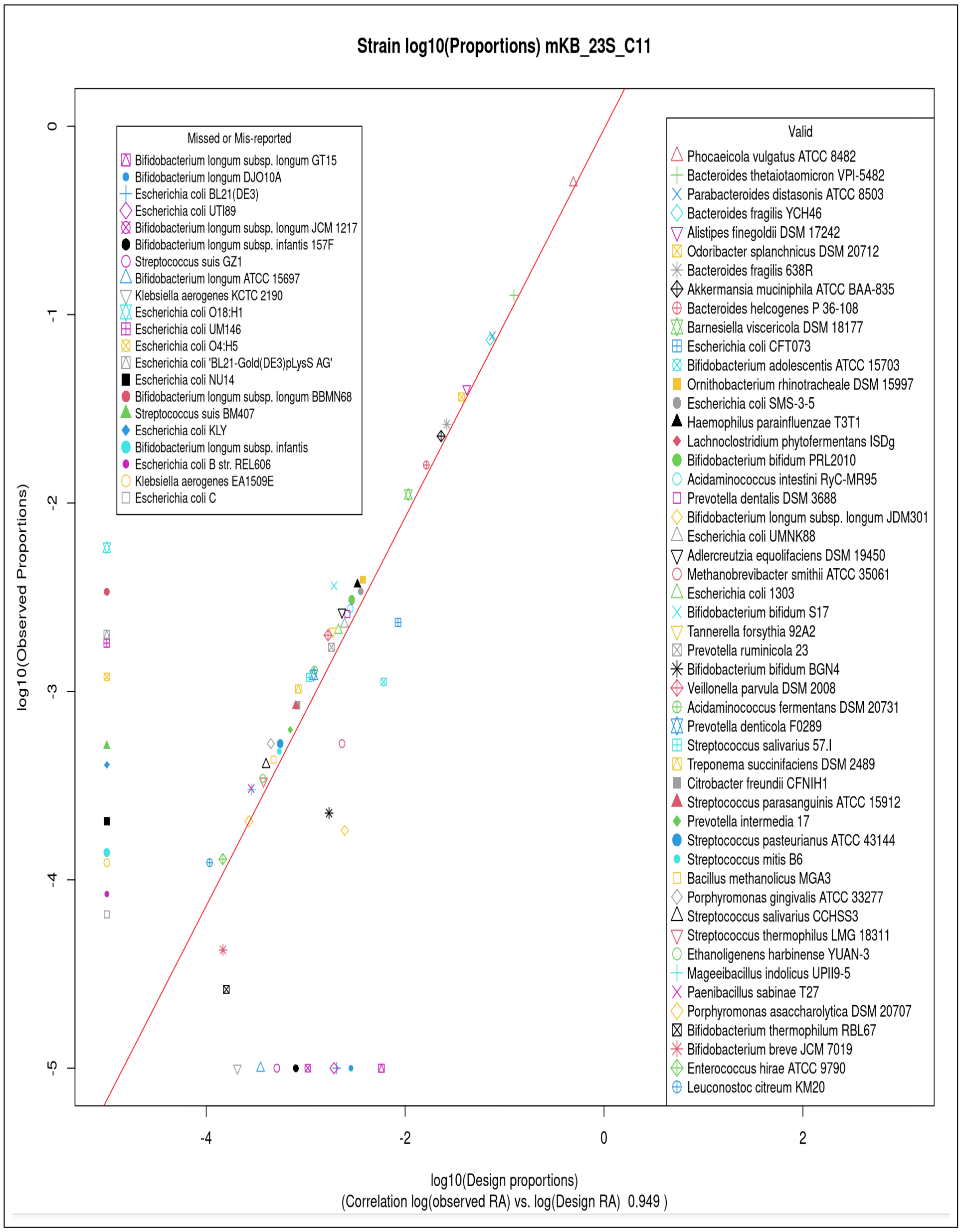


**Fig. 32** log(Observed) vs log(designed) relative abundances of the mKB microbiome as determined from 23S rRNA gene sequence reads from the C11 simulated nanopore-sequenced reads, with anomalous reads analysis incorporated.

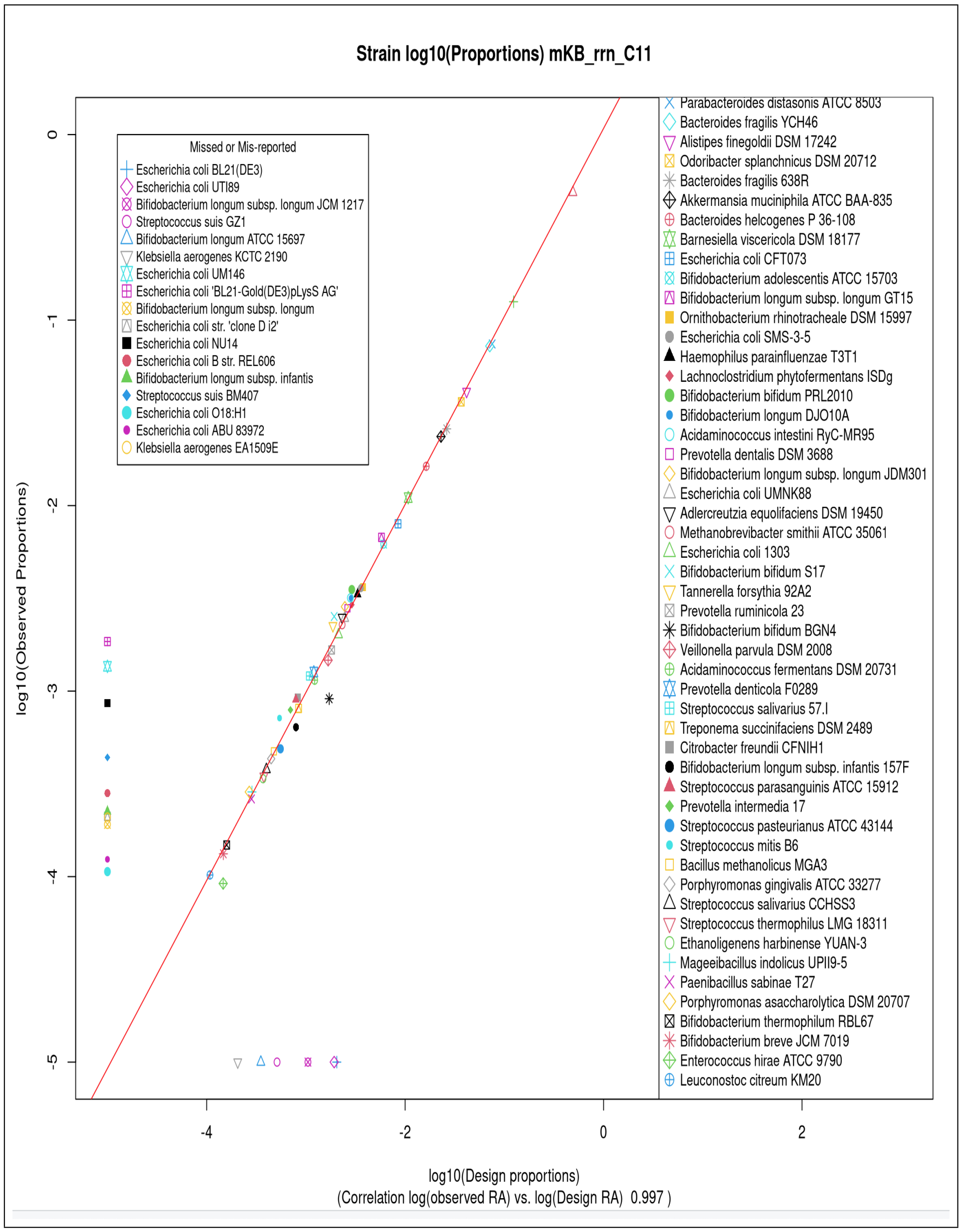


**Fig. 33** log(Observed) vs log(designed) relative abundances of the mKB microbiome as determined from *rrn* operon sequence reads from the C11 simulated nanopore-sequenced reads, with anomalous reads analysis incorporated.

#### 6.5 Results from processing the mKB_16S_C11 and mKB_23S_C11 anomalous reads

The analysis method of section 6.2 was also used for the 16S and 23S rRNA gene datasets. The results are shown in Figure 34.

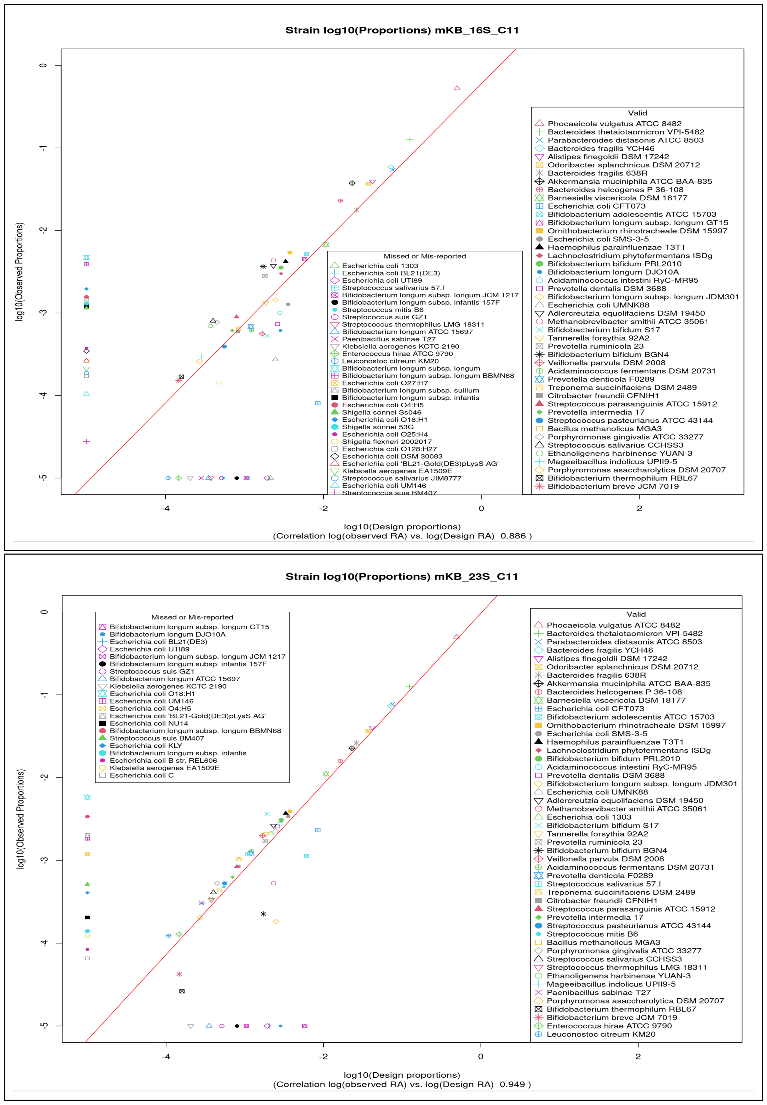


**Fig. 34** Scatterplot of observed vs. designed strain cellular relative abundance for datasets `mKB_16S_C11` and `mKB_23S_C11` with the anomalous reads results incorporated - 16S rRNA gene (upper sub-figure) and 23S rRNA gene sequences (lower sub-figure). The red lines show a linear model fit to the data not including missed or invalid observations.

### 7 Primers used for 16S rRNA gene and 23S rRNA gene identification and extraction

IUPAC codes are used to characterise primers that have variants in the base at particular locations. Most hypervariable regions have multiple variants. The 16S rRNA gene boundaries are not usually coincident between successive hypervariable regions, however the 23S rRNA gene Z regions were deliberately coincident and hence only the 3' boundary is specified for each Z region.

**Table 11** Primers used for boundary identification for 16S rRNA genes

| Type | ID | Sequence | Comment |
|---|---|---|---|
| 16S rRNA gene_5' | 27F | AGAGTTTGATCMTGGCTCAG | |
| V2_5' | 97F_V | GGCGVACGGGTGAGTAA | |
| V2_3' | 338R | TGCTGCCTCCCGTAGGAGT | |
| V3_5' | V3F | CCAGACTCCTACGGGAGGCAG | |
| V3_3' | V3R | CGTATTACCGCGGCTGCTG | |
| V4_5' | 515F | GTGYCAGCMGCCGCGGTAA | Caporaso etal |
| V4_3' | 806R | GGACTACHVGGGTWTCTAAT | Caporaso etal |
| V5_5' | 784F | AGGATTAGATACCCT | |
| V5_3' | 908F_KW | ACTCAAAKGAATWGACGG | probeBase |
| V5_3' | 908R_M | TACGGYTACCTTGTTACGACTT | probeBase |
| V6_5' | V6F | TCGATGCAACGCGAAGAA | Chakravorty et al. |
| V6_3' | V6R | ACATTTCACAACACGAGCTGACGA | Chakravorty et al. |
| V7_5' | 1100.F16 | CAACGAGCGCAACCCT | |
| V7_3' | 1237F | GGGCTACACACGYGCWAC | |
| V9_5' | 1381R | GACGGGCGGTGTGTRCA | |
| V9_3' | 1492R | TACGGYTACCTTGTTACGACTT | |
| 16S rRNA gene_3' | 1492R | TACGGYTACCTTGTTACGACTT | |

Primers for the 16S rRNA gene boundary identification, specified using IUPAC DNA base ambiguity codes.

Table 12 presents results from unpublished original work by Zhang [3]. Multiple sets of quasi-randomly chosen 23S rRNA gene sequences from Refseq were created. Multiple sequence alignment was performed on each set and then per base Shannon entropy plots generated. Based on these, candidate locations for primers (low entropy regions of approximately 15 bases) were selected and the sequences in these regions examined to create a potential primer. Primers were then evaluated across separate random sets of sequences and progressively refined, resulting in the primers documented here. Full details are available in the thesis [3]. Note that in the final analysis this work only used the 129F [12] and 2428R [13] primers.

**Table 12** Primers used for boundary identification for 23S rRNA genes

| Type | ID | Sequence | Comment |
|---|---|---|---|
| 23S rRNA gene_5' | 129F_YV | CYGAATGGGGVAACC | Hunt 2006 |
| Z0_3' | Z0F_84 | GAABTGAAACATCTHAGTA | Zhang 2021 |
| Z1_3' | Z1F_136 | AGTAGYGGCGAGCGAA | Zhang 2021 |
| Z2_3' | Z2F_384 | AGTACYGTGARGGAA | Zhang 2021 |
| Z3_3' | Z3F_728 | WRATAGCTSGTWCTC | Zhang 2021 |
| Z4_3' | Z4F_986 | GTTRGCTYRGAAGCAGC | Zhang 2021 |
| Z5_3' | Z5F_1588 | AGGAAYTMKGCAA | Zhang 2021 |
| Z6_3' | Z6F_1723 | TGAYRCCTGCCCRGTGC | Zhang2021 |
| Z7_3' | Z7F_1821 | TCCTAAGGTAGCGAAATTCCTTG | Zhang 2021 |
| Z8_3' | Z8F_2144 | ACTGGGGYGGTYKCCTCC | Zhang 2021 |
| 23S rRNA gene_3' | 2241R_H | ACCGCCCCAGTHAACT | Hunt 2006 |
| 23S rRNA gene_3' | U2482R_RM | CCRAMCTGTCTCACGACG | Martijn Lind 2019 |

Primers for boundaries of hypervariable regions of the 23S rRNA gene large subunit of bacterial ribosomal RNA operons. All primers labelled Z... were generated and evaluated by Zhengming Zhang, unpublished M.Sc Thesis, University of Melbourne. [3]

It will be noted that Table 12 has two entries for the 23S rRNA gene 3' boundary which are rather distant from each other. In this work the U2482R primer [13] has been used.

### 8 Outline of key computer code and processing pipeline.

There are 4 steps in the processing pipeline for creating a mock microbiome based on the King et al GutFeeling Knowledge Base to visualisations and results output giving identification and quantification of the microbiota of the mock microbiome. These 4 steps can be executed from the ASVpipeline.sh script, one command at a time, or by direct Rscript call from a terminal session. It is necessary that the directory structure under a defined basepath be constructed as detailed in the Github repository, [14] for this study. It is also neessary to download various pre-computed files that need to be located within that directory structure. These are located on Figshare at [15] and [16]. The following sub-section details where the various files need to be located.

#### 8.1 Placing files downloaded from Figshare

Files in the repository [16] consist of the blastn databases labelled blastdb1 and blastndb2. These need to be located in the corresponding folders of ..basepath/GROND and unzipped. The *.xlsx files are text files which are to be located in ..basepath/text

Files in the repository download of [15] consist of 10 datasets from the Sereika source in a single tarred file, together with 8 compressed files giving the 8 datasets from the Srinivas source. All these files are placed into ..basepath/fastq directory.

Below we detail the 4 calls, and then we give more detail on what calculations are done by the various steps.

#### 8.2 The 4 calls

The can be run from bash script ASVpipeline.sh by making minor edits to that script according to which of the calls detailed below you wish to use.

1. Create operon store for simulated reads mock microbiome - e.g. mockKB_23S_rrn

```
Rscript --vanilla make_mock_operon_store.R
             /vast/projects/rrn/ASVtest
               mockKB 23S 11 30 4 291 59 30000 12
```

2. Create the fastq file that is the read library for a simulated reads mock microbiome. Denoise the microbiome created or one that is from real sequencing.

```
Rscript --vanilla make_mock_and_denoise.R <basepath>
           whichMock whichSubunit whichCase whichPair
             uppThreshErrRate totOperons totalStrains numcores

e.g. for mockKB_23S_C11
Rscript --vanilla make_mock_and_denoise.R
            vast/projects/rrn/ASVtest mockKB 23S
             11 0 00.01 291 59 2

e.g. for mSriSA2\_rrn\_C03
```

```
Rscript --vanilla make_mock_and_denoise.R
           /vast/projects/rrn/ASVtest mSriSA2 rrn
             03 0 2 0.015 0 0 2
```

```
e.g. for mSerS3\_16S\_03
Rscript --vanilla make_mock_and_denoise.R
           /vast/projects/rrn/ASVtest mSerS3 16S
             03 3 00.01 0 0 2
```

3. Generate kmer spectra for the ASVs and the reads presented for denoising.

```
sbatch --mem=240GB --time=00:20:00 ASVkmerSpectra.sh <basepath>
          whichMock whichSubunit whichCase whichSubMock
            whichPair uppThreshErrRate numcores
```

```
e.g. sbatch --mem=240GB --time=00:10:00 ASVkmerSpectra.sh
               /vast/projects/rrn/ASVtest mSriSA2 rrn
                 03 0 2 0.015 56
```

4. Run blastn on the ASVs, then analyse the alignment data to identify and quantify the strains, species and genera present in the mock microbiome.

```
Rscript --vanilla blastn_call_ident_quant.R <basepath>
           whichMock whichSubunit whichCase whichSubMock whichPair
```

```
            uppThreshErrRate totalNumberOperons totalNumberStrains numcores
```

```
e.g. for mSriSA2_rrn_03 with error rate threshold 0.0125

Rscript --vanilla blastn_call_ident_quant.R /vast/projects/rrn/ASVtest

            mSriSA2 rrn 03 0 2 0.0125 0 0 2
```

Running this call for mockKB is likely to lead to exceeding the allowed resources on a terminal session. Hence a slurm job is needed, via the shell script blastnCallIDQuant.sh which has the same 10 parameters.

```
sbatch --mem=100GB --time=4:00:00 blastnCallIDQuant.sh  basepath

          whichMock  whichSubunit whichCase whichSubMock

            whichPair uppErrRate maxopnum nmock 56
```

```
e.g. sbatch --mem=240GB --time=48:00:00 blastnCallIDQuant.sh

                /vast/projects/rrn/ASVtest mockKB 23S

                  11 0 0 0.01 291 59 16
```

#### 8.3 Summary of what is done through each call

The 3 items below are executed through `make_mock_store.R`

- Select the operons of the strains that are in the mock microbiome and generate individual fastq files for each one.

- Determine the number of simulated reads to be generated for each operon, based on an approximate number of reads that the library of the pre-filtered mock microbiome is to have for the operons of the most abundant strain.
- For each operon submit a batch job that generates a moderate excess of fastq sequences over the number actually required for the mock microbiome dataset.

The 2 items below are executed through `make_mock_denoise.R`

- On completion of simulated read generation for all operons, randomly select the required number of reads for each operon and store as a single, multifastq file. This is the dataset of the designed mock microbiome.
- Submit this file for quality and length filtering followed by RAD denoising. These are all carried out in Julia code, the core of which is Kumar et al.'s Robust Amplicon Denoising code, RAD. The Julia script used here is a simple elaboration of that provided by Ben Murrell ([2], and private communication).

For anomalous reads analysis a dis-similarity matrix for the associated reads for each ASV is required. The dis-similarity measure used is a kmer-based edit distance calculated from the kmer spectra of each read. This is calculated at this stage. This is computed by code `ASV_reads_kmerSpectra.R` which can be run as a slurm job using the batch script ASVkmerSpectra.sh called as sbatch –mem=128GB –time=8:00:00 ASVcode/ASVkmerSpectra.sh 16S 12 mSerS 2 `Case_03` for dataset `mSerS2_16S_C03` using 12 threads.

The 7 items below are executed through `blastn_call_ident_quant.R`

- Set up characterising data on the mock microbiomes, such as strain names, number of operons for each strain, relative abundances.
- Read the key text files created as output by RAD specifying the sequence of each ASV generated and the reads associated with each ASV.
- Set up a call of blastn to align each ASV against the reference strain database. We call blastn via R's system2() function. Call blastn twice to create the desired text file output for further analysis.
- Parse the blastn output to determine the best match of each ASV sequence to the operons in the database, breaking ties for best alignment by selecting the first listed database entry. Each ASV then has a strain label.
- Save the read counts for each labelled ASV. Using the number of operons for the relevant strain convert operon counts to cellular counts to each strain, and then return the relative cellular abundance of each strain identified, distinguishing between strains that were in the designed mock microbiome and those not in that microbiome.
- Generate 2D UMAP projection plots that illustrate the identification performance being achieved with the analysis.
- Generate scatterplots of observed proportions vs. designed (or expected) proportions - or the logarithm of these quantities - showing the quantification performance of the analysis.

If the analysis is to be extended to anomalous reads then code `anomalous_reads.R` is run. That code generates scatterplots and 2D UMAP projection plots analogous to those of `blastn_call_ident_quant.R` but with the anomalous reads also taken account of.

### References


[1] Mantas Sereika et al. "Oxford Nanopore R10.4 long-read sequencing enables the generation of near-finished bacterial genomes from pure cultures and metagenomes without short-read or reference polishing". In: *Nature Methods* 19 (July 2022), pp. 823–826. DOI: 10.1038/s41592-022-01539-7.

[2] Benjamin Murrell. *seqUMAP*. 2021.

[3] Zhengming Zhang. "Microbiome Analysis using the 23S rRNA Gene with Nanopore Sequencing Data". (unpublished) MC-SCIBF University of Melbourne. 2021.

[4] James Melville. *uwot: The Uniform Manifold Approximation and Projection (UMAP) Method for Dimensionality Reduction*. R package version 0.2.3. 2025. DOI: 10.32614/CRAN.package.uwot. URL: https://CRAN.R-project.org/package=uwot.

[5] Charles H. King et al. "Baseline human gut microbiota profile in healthy people and standard reporting template". In: *PLoS One* 14.9 (Sept. 2019), e0206484. DOI: 10.1371/journal.pone.0206484. URL: https://doi.org/10.1371/journal.pone.0206484.

[6] Ryan R. Wick. "Badread: simulation of error-prone long reads". In: *Journal of Open Source Software* 4.36 (2019), p. 1316. DOI: 10.21105/joss.01316. URL: https://doi.org/10.21105/joss.01316.

[7] Ryan R. Wick. 2024. URL: https://github.com/rrwick/Badread.

[8] Venkatesh Kumar et al. "Long-read amplicon denoising". In: *Nucleic Acids Research* 47.18 (Aug. 2019), e104. URL: https://doi.org/10.1093/nar/gkz657.

[9] Calum J. Walsh et al. "GROND: a quality-checked and publicly available database of full- length 16S-ITS-23S rRNA operon sequences". In: *Microbial Genomics* 10.001255 (June 2024). DOI: 10.1099/mgen.0.001255.

[10] Zachary T Lewis et al. "Validating bifidobacterial species and subspecies identity in commercial probiotic products". In: *Pediatric Research* 79 (Mar. 2016), pp. 445–452. DOI: https://doi.org/10.1038/pr.2015.244.

[11] Ghazal Aziz, Arsalan Zaidi, and Muhammad Tariq. "Compositional quality and possible gastrointestinal performance of marketed probiotic supplements". In: *Probiotics and antimicrobial proteins* 14 (Apr. 2022), pp. 288–312. DOI: https://doi.org/10.1007/s12602-022-09931-7.

[12] Dana E. Hunt et al. "Evaluation of 23S rRNA PCR Primers for Use in Phylogenetic Studies of Bacterial Diversity". In: *Applied and Environmental Microbiology* 72.3 (Mar. 2006), pp. 2221–2225. DOI: 10.1128/AEM.72.3.22212225.2006.

[13] Joran Martijn et al. "Confident phylogenetic identification of uncultured prokaryotes through long read amplicon sequencing of the 16S-ITS-23S rRNA operon". In: *Environmental Microbiology* 21.7 (July 2019), pp. 2485–2498. DOI: 10.1111/1462-2920.14636. URL: https://doi.org/10.1111/1462-2920.14636.

[14] Chris Woodruff. *ASVpaper Rcode and processing pipeline scripts.* (Accessed 28 January 2026). 2026. URL: https://github.com/cjwoodruff50/ASVpaper.

[15] Chris Woodruff. *10 Sereika datasets and 8 Srinivas datasets.* (Accessed 28 January 2026). URL: https://doi.org/10.6084/m9.figshare.31052200.v1.

[16] Chris Woodruff. *Compressed files providing blastn databases used for the ASV paper analyses, together with gut microbiome definition files.* (Accessed 28 January 2026). URL: https://doi.org/10.6084/m9.figshare.31041886.v1.
